# Activity of Carbazole, Aminoguanidine and Diamine Anti-infectives against *Toxoplasma gondii*

**DOI:** 10.1101/2025.02.15.638445

**Authors:** Davinder Singh, Melissa A. Sleda, Akanksha Pandey, Satish R. Malwal, Yiyuan Chen, Ruijie Zhou, Feyisola Adewole, Katie Sadowska, Oluseye K. Onajole, Silvia N.J. Moreno, Eric Oldfield

## Abstract

We report the observation that carbazole anti-infectives developed as antibacterial and antifungal drug leads have activity against the tachyzoite-stage growth of the Apicomplexan parasite *Toxoplasma gondii* with IC_50_ values as low as 2 μM. We show that a phenylthiazole aminoguanidine with antibacterial as well as antifungal activity inhibits growth with an IC_50_ value of 2.1 μM. We also tested a series of 18 analogs of the diamine SQ109, a tuberculosis drug candidate which likewise has both antibacterial and antifungal activity, finding activity as low as 2.3 μM. We tested all compounds for their activity in collapsing the ΔpH component of the promotive force, the results indicating that all compounds acted, at least in part, as protonophore uncouplers. Finally, we also investigated the correlation between the activity of all compounds against the yeast *Saccharomyces cerevisiae* and the bacterium *Mycobacterium smegmatis*, finding significant correlations between the collapse of the proton motive force and anti- fungal/antibacterial activity.

---

Toxoplasmosis, a parasitic infection caused by the protozoan *Toxoplasma gondii*, poses a significant health concern with an estimated 8–22% of people in the USA being infected.^1^ The global prevalence of *T. gondii* infection is high, with estimates suggesting that one-third of the world’s population is infected.^2^ *T. gondii* is a ubiquitous parasite with a complex life cycle involving both definitive and intermediate hosts. Cats are the definitive hosts, excreting oocysts in their feces that can contaminate soil, water, and food. Humans and other animals serve as intermediate hosts, acquiring the parasite through ingestion of undercooked meat, contaminated water, or soil. Additionally, congenital transmission can occur from an infected mother to her fetus.^3^ While most infections are asymptomatic, a substantial proportion of the infected population harbors the parasite in a latent state, potentially reactivating in immunosuppressed patients.^4^ Reactivation of the infection in immunocompromised individuals, such as those with HIV/AIDS, organ transplantation recipients, or individuals undergoing cancer treatment, can progress to severe and life-threatening disease, affecting the central nervous system, eyes, and other organs.^5–7^ Congenital toxoplasmosis can result in severe birth defects, including intellectual disability, visual impairment, and seizures.

The standard treatment for toxoplasmosis involves a combination of pyrimethamine and sulfadiazine,^8^ often with the addition of leucovorin to reduce bone marrow toxicity. While this regimen is effective in many cases, it can have significant side effects, including bone marrow suppression, gastrointestinal disturbances, and skin rashes. There is, therefore, a need for novel drugs to address the limitations of existing therapies. The potential for severe side effects, and the challenges associated with long-term treatment highlight the desirability of developing new therapeutic approaches. Recent years have witnessed significant progress in the identification of novel drug targets and the development of potential therapeutic agents for toxoplasmosis. Several promising drug leads have been identified, including inhibitors of parasite-specific enzymes and host-pathogen interactions. However, progress in targeting the persister-like (bradyzoite) stages,^9^ as well as the development of drugs that increase innate immunity, have been limited.

One approach to finding new drugs to treat toxoplasmosis is to repurpose drugs already in use or in development to treat other infectious diseases^10–12^. In previous work^13, 14^ we investigated a class of drugs called bisphosphonates, used to treat bone resorption disorders. These compounds target the enzymes of the *T. gondii* isoprenoid synthesis pathway^15^ and in past work we found promising results with n-alkyl bisphosphonates in animal models^13,14^. More recent work with lipophilic, nitrogen-containing bisphosphates showed their potent activity against *T. gondii* growth by inhibiting the long prenyl synthase TgCoq1 resulting in inhibition of the formation of ubiquinones,^16^ leading to good activity in an animal model of infection.^17^ We also investigated the tuberculosis drug candidate SQ109, a molecule which targets many enzymes in *Mycobacterium tuberculosis* including MmpL3, MenA, MenG^18^ as well as undecaprenyl diphosphate phosphatase, the *E. coli* analog of the *M. tuberculosis* enzyme decaprenyl diphosphate phosphatase, essential for peptidoglycan biosynthesis.^19^ We found activity against the tachyzoite form of *T. gondii* with an IC_50_ = 1.8 μM as well as an 80% survival in a mouse model of infection.^20^

Additionally, several groups have been investigating the carbazole class of molecules as antibacterials and antifungal agents. Chao and coworkers^21^ investigated 22 carbazoles against *Candida albicans* SC5314. Most compounds were inactive (MIC>64 μg/mL) but CAR-20 (**1**, Figure 1), also known as THCz-1^22^ had an MIC=4 μg/mL and CAR-8 (THCz-2, **2**) had an MIC=2 μg/mL. They proposed that the mechanism of action was by inducing endoplasmic reticulum stress. Interestingly, in earlier work, Reithuber and coworkers^22^ discovered that the same molecules also had activity against bacteria, and they proposed that the mechanism of action involved inhibition of cell wall biosynthesis, with the inhibitors binding to isoprenoid molecules containing diphosphate groups, the analog **3** being inactive. In the yeast study^21^, there were also promising results in the *Galleria mellonella* animal model of infection, as well as the interesting observation that THCz-2 (CAR-8) had potent synergistic activity (FICI = 0.28) with fluconazole in a fluconazole-resistant strain of *Candida albicans*.

**Figure 1.**
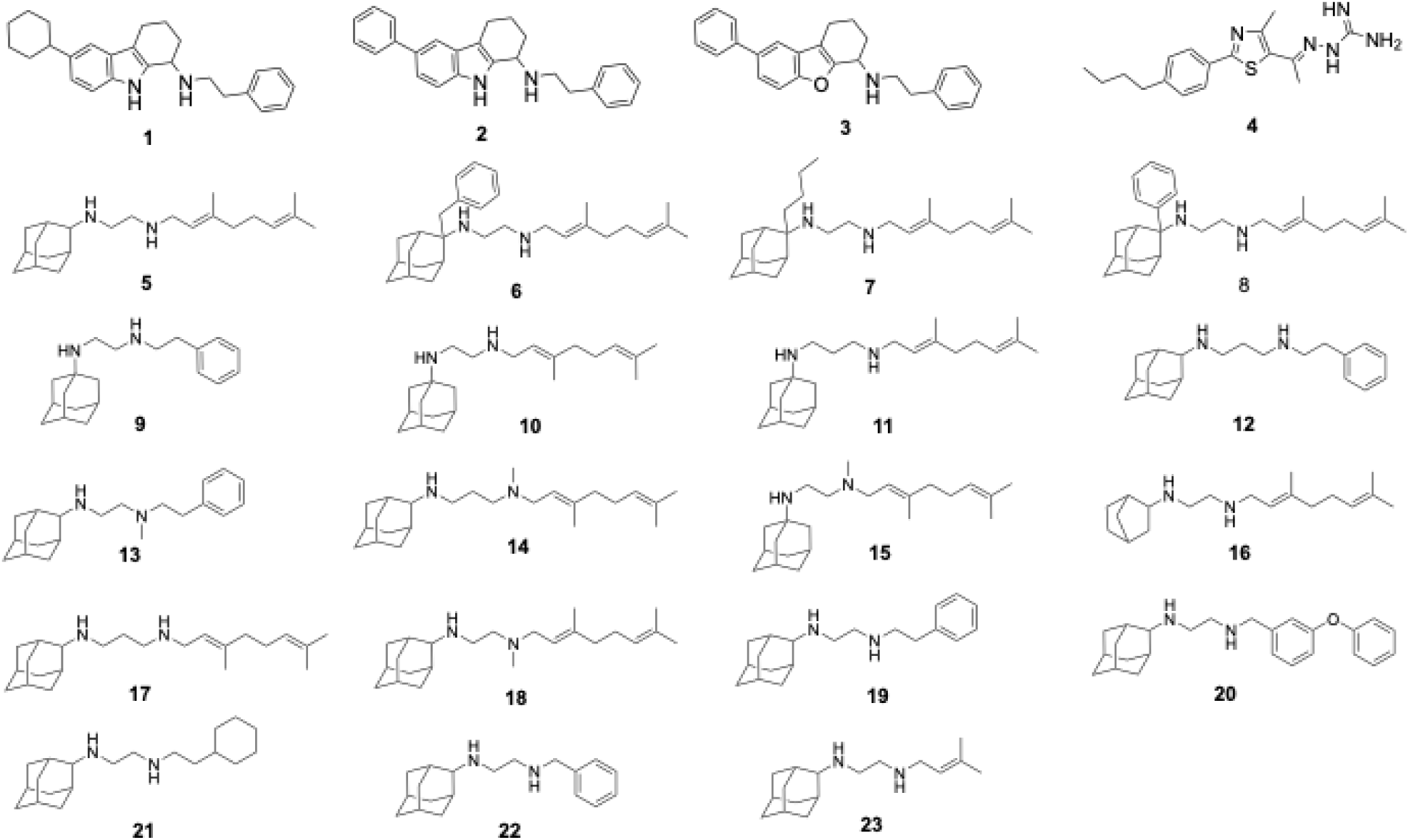
Structures of THCz-1 (**1**), THCz-2 (**2**), THCz-6, (**3**) Thiazole (**4**), SQ109 (**5**), and 18 other SQ109 analogs (**6**-2**3**) investigated in this work.

A third series of molecules of interest are the aminoguanidines, in particular phenylthiazole species such as **4** (Figure 1) developed by Seleem and coworkers, with both antifungal^23^ and antibacterial activity.^24–26^ Chemogenomic profiling against more than 6,000 heterozygous diploid deletion strains of *Saccharomyces cerevisiae* tested, in addition to heterozygous strains of *Candida* encoding homologous genes in the dolichyl phosphate pathway, did not clarify their mechanism of action.^26, 27^ It is thus possible their activity is due to a more “physical” mechanism of action, although it was noted by Seleem et al.^23^ that the phenylthiazole, **4,** did not disrupt either bacterial or fungal cell membranes. Here, we thus investigated the activity of three carbazoles, the phenylthiazole aminoguanidine as well as 18 analogs of SQ109, against the *in vitro* growth of *T. gondii* tachyzoites.

## RESULTS AND DISCUSSION

We first investigated the activity of THCz-1 (**1**, Figure 1), THCz-2 (**2**) and THCz-6 (**3**) (the nomenclature used in Reference 22), and of the phenylthiazole **4** against *T. gondii* tachyzoites, then investigated the activity of SQ109 and a series of 18 SQ109 analogs using the same assay.

As can be seen in Figure 2a, b, the activity of THCz-1 and THCz-2 is ∼2 μM, similar to the activity observed previously with SQ109 (IC_50_ =1.8 μM).^20^ The analog THCz-6 was less active, Figure 2c. (IC_50_ = 7.5 μM). There was no difference in activity between the cyclohexyl (THCz-1) and phenyl (THCz-2) species, so the difference in activity between THCz-1/2 and THCz-6 could be due to replacement of the carbazole nitrogen with oxygen in THCz-6. Based on our results with SQ109 in which we found that SQ109 had activity as a protonophore uncoupler,^28^ we hypothesized that the THCz species would also be uncouplers and that the oxygen might decrease activity due to an electron-withdrawing effect. We thus next tested the effects of THCz-1, THCz- 2 and THC-6 in the *E. coli* inverted membrane vesicle (IMV)/ACMA fluorescence assay^28^ finding that THCz-1 (Figure 2d) and THCz-2 (Figure 2e) both had potent activity in collapsing the ΔpH component of the proton motive force with IC_50_ values of <1 μM whereas THCz-6 was much less active with an IC_50_ of 5.4 μM, Figure 2f.

**Figure 2.**
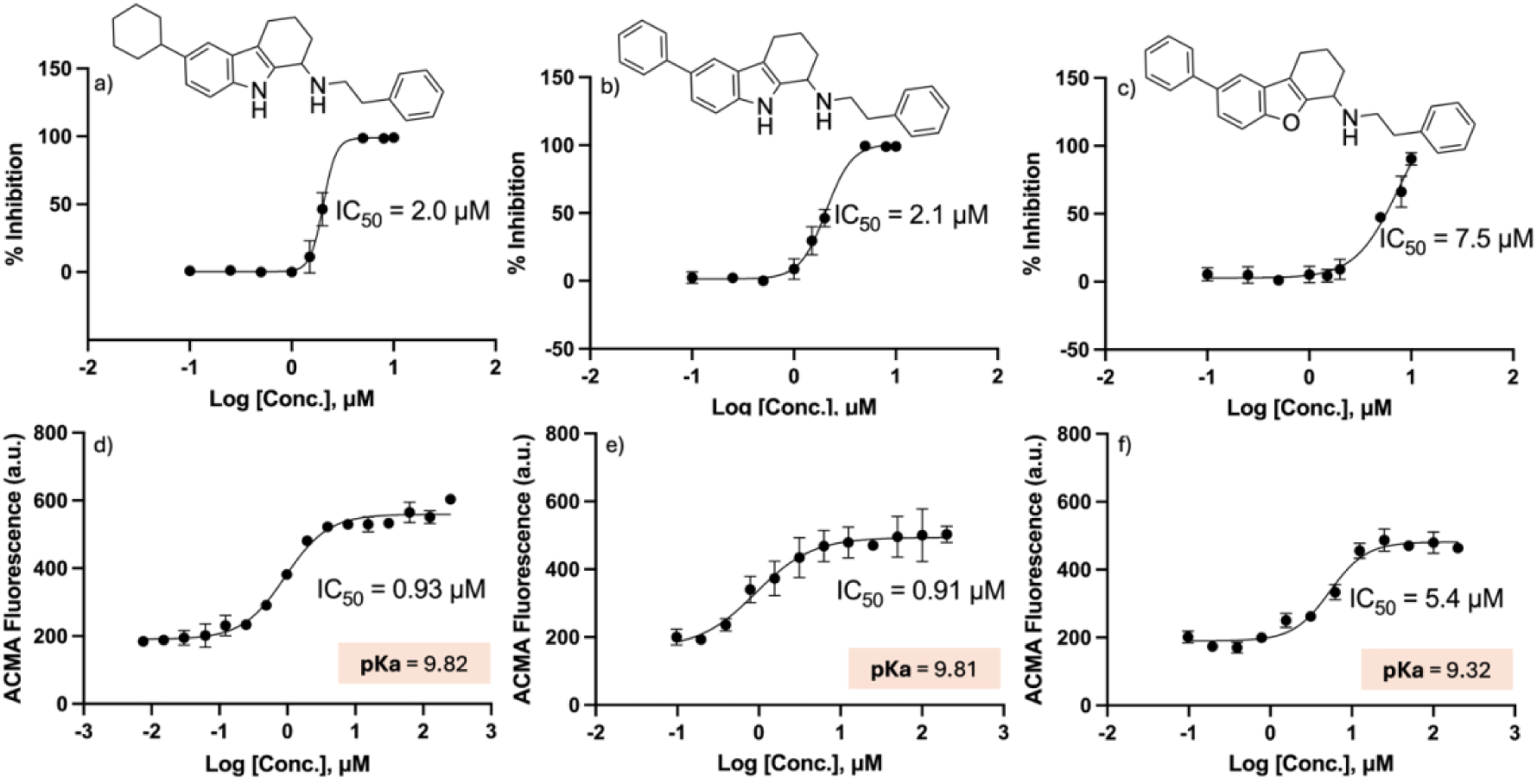
*Toxoplasma gondii* growth inhibition and ΔpH collapse results in *E. coli* inverted membrane vesicles for THCz-1 (**1**), THCz-2 (**2**) and THCz-6 (**3**). (a-c), Growth inhibition results. a) THCz-1. b) THCz-2. c) THCz-6. Each bar represents the standard deviation of the experimental points of at least three independent experiments. (d-f), Dose-response curves for pH gradient collapse using *E. coli* IMVs and ACMA fluorescence. d) THCz-1. e) THCz-2. f) THCz-6. The salmon shaded rectangles show the computed pKa values (chemicalize.com).

We next investigated the activity of the phenylthiazole, which showed an IC_50_ of 2.1 μM in the tachyzoite assay, Figure 3a. There was also an effect on the ΔpH component of the proton motive force with an IC_50_ of 6.7 μM, Figure 3b, although this was much less than the effect of FCCP, Figure 3c. The amino-guanidine sidechain in the phenylthiazole is computed to have a pKa value of ∼9 so is less effective as a protonophore uncoupler. However, since the inner mitochondrial membrane maintains a negative membrane potential, essential for ATP production, cationic lipophilic species, being positively charged and lipid-soluble, can readily cross the membrane and collapse the ΔΨ component of the PMF. This effect is seen in the IMV assay with the lipophilic base SQ109 where both the ΔpH as well as the ΔΨ components are affected.^28^ In addition, SQ109 also collapses the ΔΨ component of the PMF in the protozoan parasite, *T. cruzi*,^29^ making it likely that the ΔΨ is also decreased in *T. gondii* as well.

**Figure 3.**
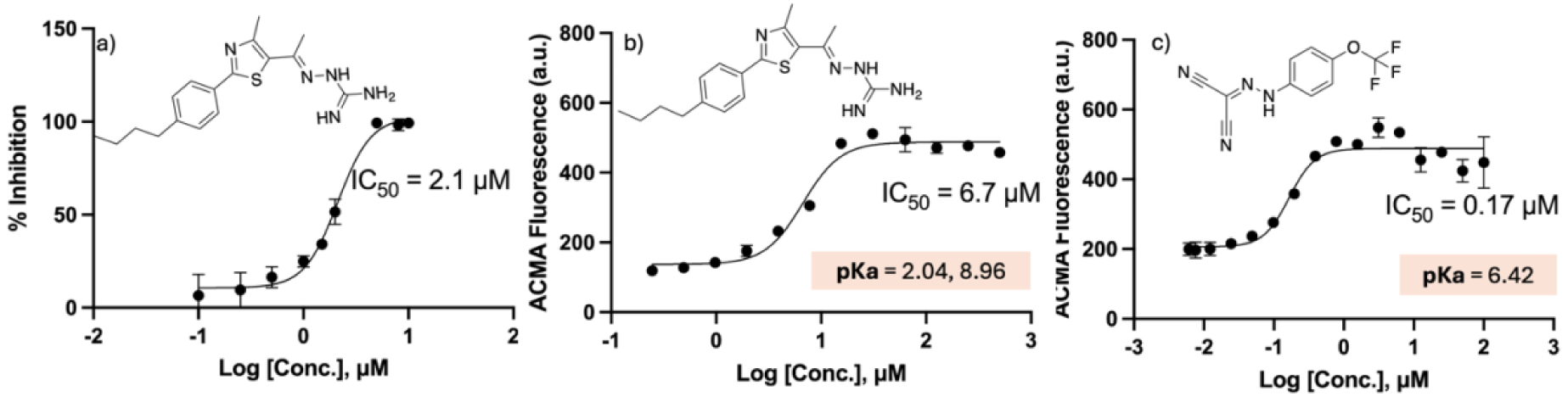
*Toxoplasma gondii* growth inhibition and IMV results. a) Dose-response curve for *T. gondii* growth inhibition by the phenylthiazole **4**. b) pH gradient collapse by **4** using *E. coli* IMVs and ACMA fluorescence. c) pH gradient collapse by FCCP. The salmon shaded rectangles highlight the pKa values (chemicalize.com).

Next, we synthesized a series of SQ109 analogs and tested these as well as a series of other analogs^30, 31^ for activity against tachyzoite growth. Results for SQ109 and two analogs are shown in Figures 4a-c and data for all compounds are shown in Table 1.

**Figure 4.**
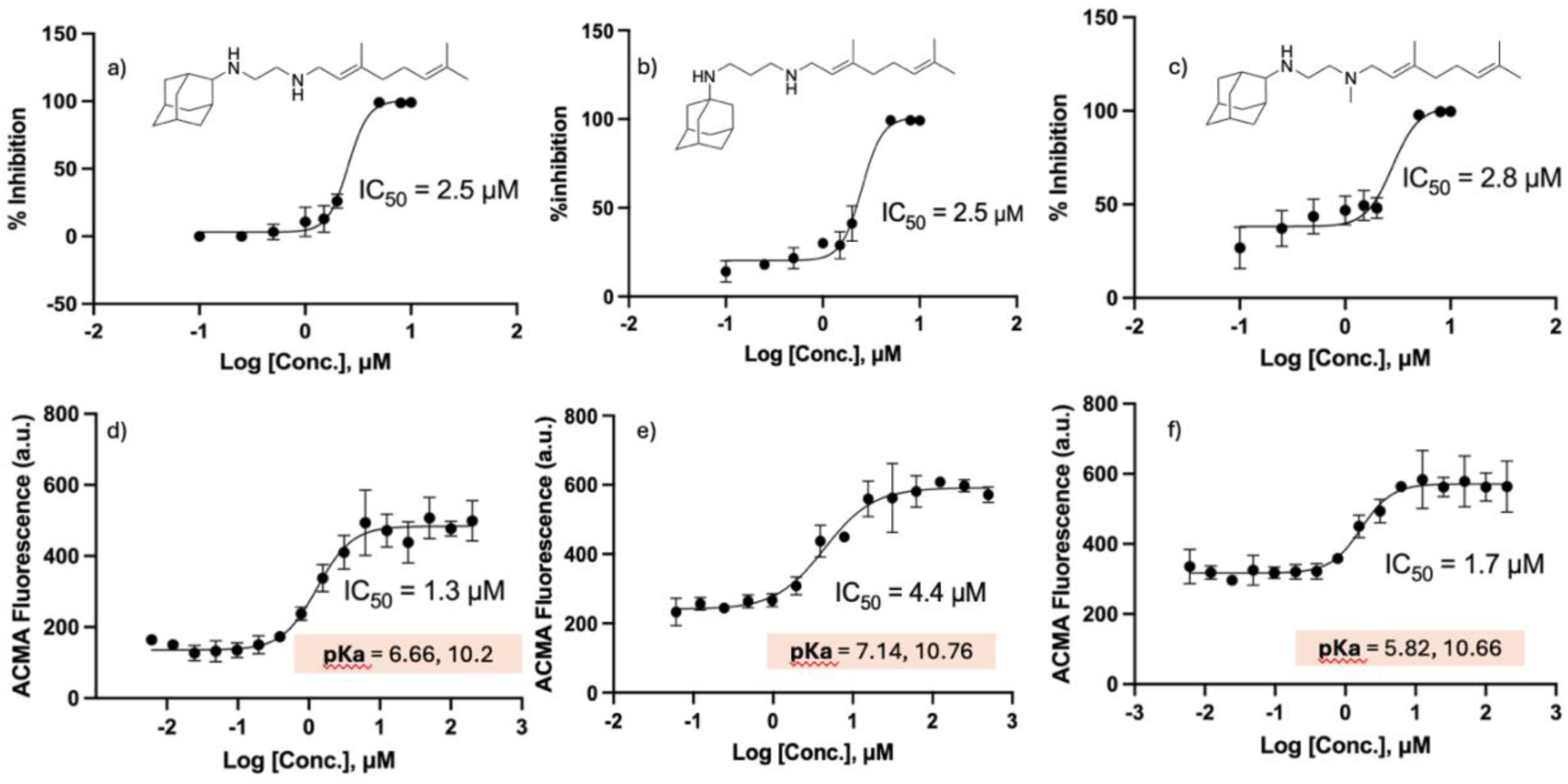
*Toxoplasma gondii* growth inhibition and uncoupling activity of SQ109 (**5**), **11** and **18**. (a-c), Growth inhibition for: a) **1**. b) **11**. c) **18**. (d-f) Dose-response curves for pH gradient collapse using *E. coli* IMVs and ACMA fluorescence. d) **1**. e) **11.** f) **18.** The shaded rectangles show the computed pKa values (chemicalize.com).

**Table 1.**
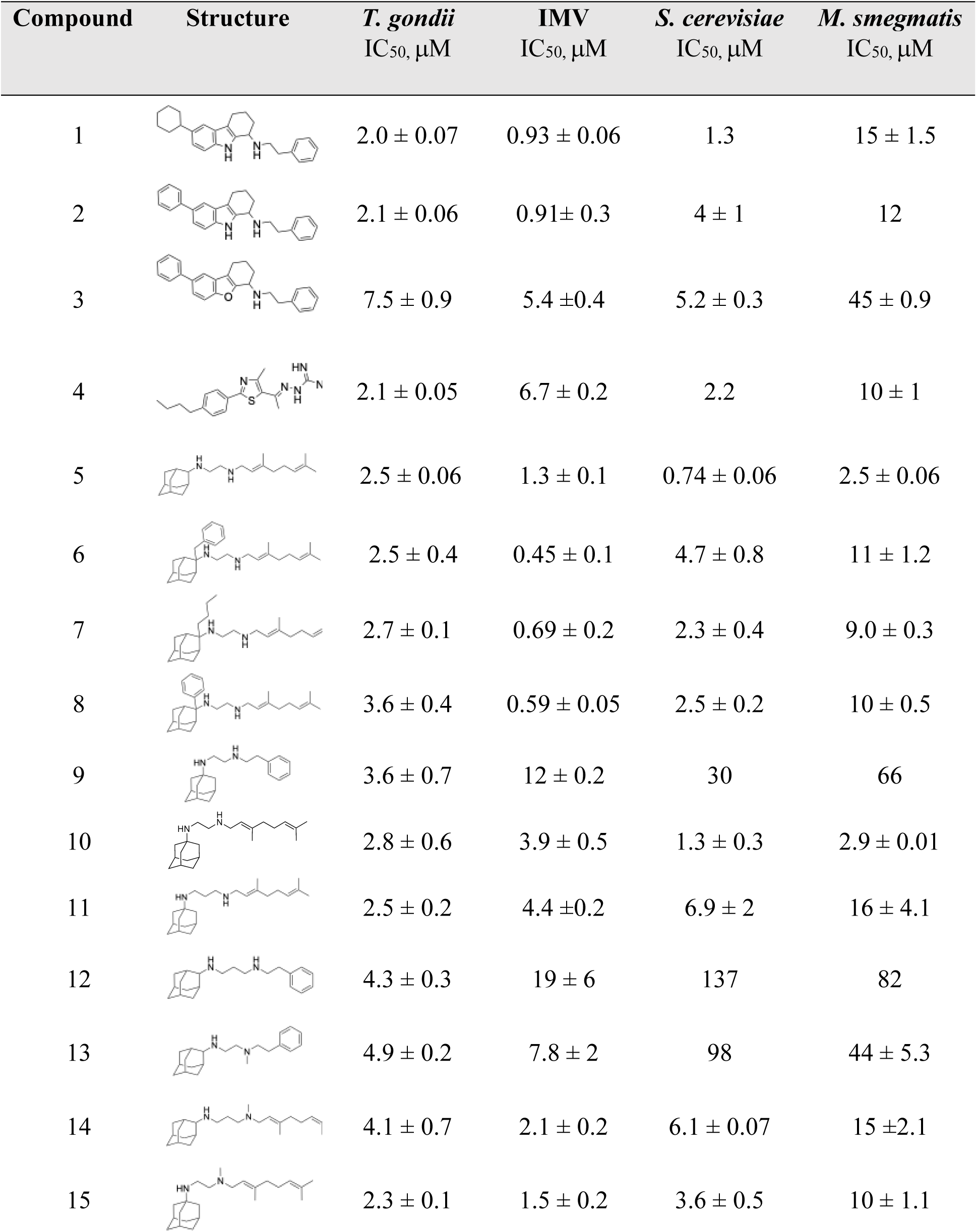
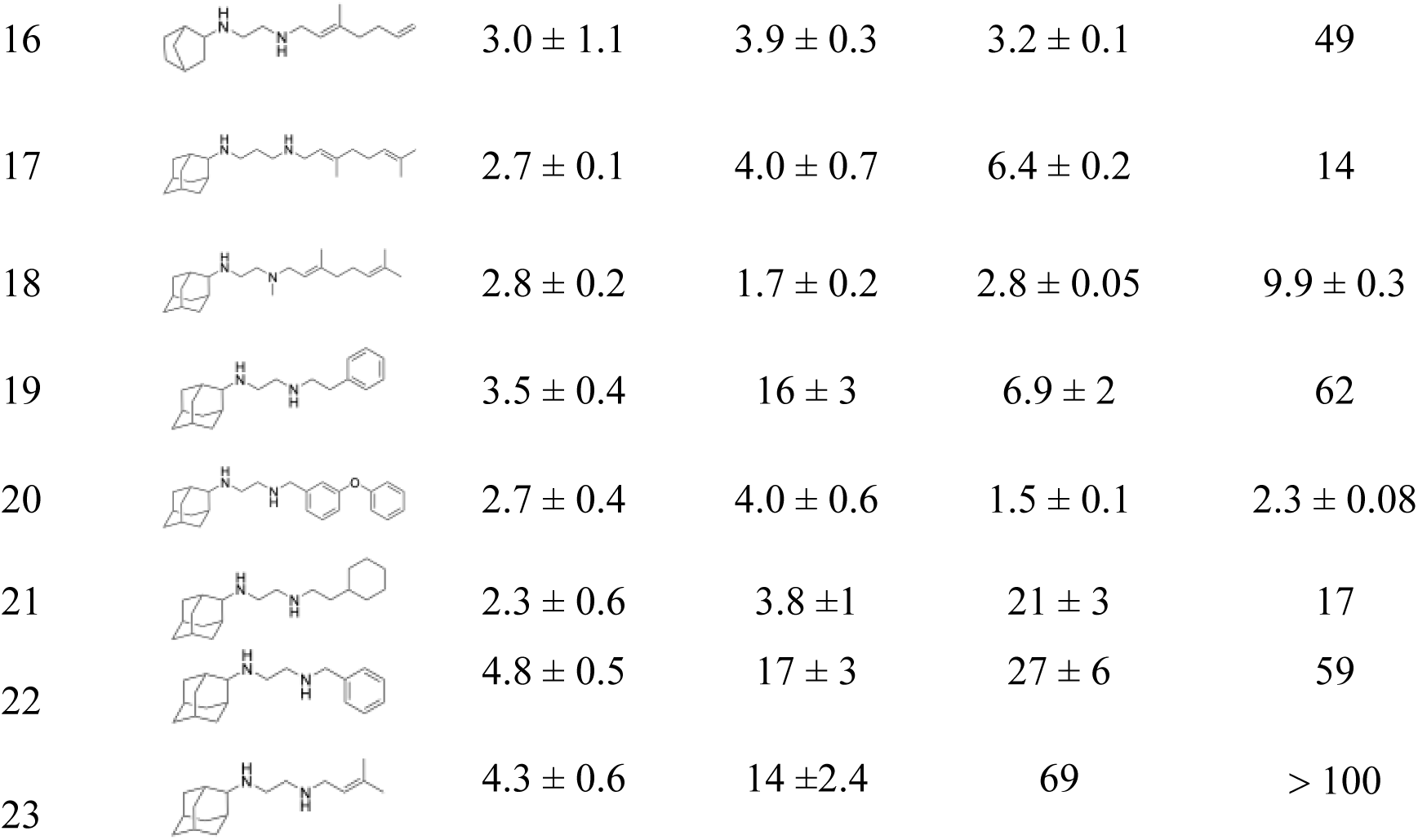
IMV and cell growth inhibition data for THCzs, phenylthiazole, SQ109 and its 18 analogs.

| Compound | Structure | *T. gondii*<br>IC<sub>50</sub>, μM | IMV<br>IC<sub>50</sub>, μM | *S. cerevisiae*<br>IC<sub>50</sub>, μM | *M. smegmatis*<br>IC<sub>50</sub>, μM |
| --- | --- | --- | --- | --- | --- |
| 1 |  | 2.0 ± 0.07 | 0.93 ± 0.06 | 1.3 | 15 ± 1.5 |
| 2 |  | 2.1 ± 0.06 | 0.91 ± 0.3 | 4 ± 1 | 12 |
| 3 |  | 7.5 ± 0.9 | 5.4 ± 0.4 | 5.2 ± 0.3 | 45 ± 0.9 |
| 4 |  | 2.1 ± 0.05 | 6.7 ± 0.2 | 2.2 | 10 ± 1 |
| 5 |  | 2.5 ± 0.06 | 1.3 ± 0.1 | 0.74 ± 0.06 | 2.5 ± 0.06 |
| 6 |  | 2.5 ± 0.4 | 0.45 ± 0.1 | 4.7 ± 0.8 | 11 ± 1.2 |
| 7 |  | 2.7 ± 0.1 | 0.69 ± 0.2 | 2.3 ± 0.4 | 9.0 ± 0.3 |
| 8 |  | 3.6 ± 0.4 | 0.59 ± 0.05 | 2.5 ± 0.2 | 10 ± 0.5 |
| 9 |  | 3.6 ± 0.7 | 12 ± 0.2 | 30 | 66 |
| 10 |  | 2.8 ± 0.6 | 3.9 ± 0.5 | 1.3 ± 0.3 | 2.9 ± 0.01 |
| 11 |  | 2.5 ± 0.2 | 4.4 ± 0.2 | 6.9 ± 2 | 16 ± 4.1 |
| 12 |  | 4.3 ± 0.3 | 19 ± 6 | 137 | 82 |
| 13 |  | 4.9 ± 0.2 | 7.8 ± 2 | 98 | 44 ± 5.3 |
| 14 |  | 4.1 ± 0.7 | 2.1 ± 0.2 | 6.1 ± 0.07 | 15 ± 2.1 |
| 15 |  | 2.3 ± 0.1 | 1.5 ± 0.2 | 3.6 ± 0.5 | 10 ± 1.1 |
| 16 | | $3.0 \pm 1.1$ | $3.9 \pm 0.3$ | $3.2 \pm 0.1$ | 49 |
| 17 | | $2.7 \pm 0.1$ | $4.0 \pm 0.7$ | $6.4 \pm 0.2$ | 14 |
| 18 | | $2.8 \pm 0.2$ | $1.7 \pm 0.2$ | $2.8 \pm 0.05$ | $9.9 \pm 0.3$ |
| 19 | | $3.5 \pm 0.4$ | $16 \pm 3$ | $6.9 \pm 2$ | 62 |
| 20 | | $2.7 \pm 0.4$ | $4.0 \pm 0.6$ | $1.5 \pm 0.1$ | $2.3 \pm 0.08$ |
| 21 | | $2.3 \pm 0.6$ | $3.8 \pm 1$ | $21 \pm 3$ | 17 |
| 22 | | $4.8 \pm 0.5$ | $17 \pm 3$ | $27 \pm 6$ | 59 |
| 23 | | $4.3 \pm 0.6$ | $14 \pm 2.4$ | 69 | > 100 |

We next tested the activity of these compounds as uncouplers using the ACMA fluorescence assay. Results are given in Table 1 and dose-response curves for SQ109, and two analogs are shown in Figures 4d-f from which it can be seen that all compounds have potent activity as protonophore uncouplers.

However, as can be seen in Table 1, there is only a relatively small range in the IC_50_ values for *T. gondii* cell growth inhibition. Since THCzs, phenylthiazoles as well as SQ109 and their analogs have been found to have activity against bacteria and yeasts, we next determined the activity of **1**- **23** against *M. smegmatis* and *S. cerevisiae*. Results are shown in Table 1. We then compared the cell growth inhibition results (log IC_50_) with the ΔpH uncoupling results (log IC_50_ in the IMV assay) to see if there were significant correlations, which would support a role in their activity as anti-infectives. Graphical correlations are shown in Figure S1a-c in the Supporting Information and are summarized in Table 2.

**Table 2.** Summary of correlations between the ΔpH component of the proton motive force (log IC_50_ in the IMV assay) and cell growth inhibition (log IC_50_ values) for 1-23 in *T. gondii*, M. smegmatis *and* S. cerevisiae.

|  | Slope | Pearson R-value | P-value |
| --- | --- | --- | --- |
| *T. gondii* | 0.15 | 0.52 | 0.0106 |
| *M. smegmatis* | 0.57 | 0.60 | 0.0030 |
| *S. cerevisiae* | 0.83 | 0.66 | 0.0006 |

As can be seen in Table 2, the small range in IC_50_ values for cell growth inhibition with *T. gondii* results in a small slope and the smallest R-value and the largest P-value which is, however, statistically significant (p∼0.01). The slope and R-value with *M. smegmatis* increase and P decreases to P=0.003, and with *S. cerevisiae* the slope increases further (to 0.83) as does the R value, and P decreases to P=0.0006. Taken together these results indicate that uncoupling activity plays a larger role in *M. smegmatis* and *S. cerevisiae* cell growth inhibition than with *T. gondii* cell growth inhibition and as expected, *M. smegmatis* and *S. cerevisiae* cell growth inhibition are highly correlated (R=0.77, p=0.00003), as shown in Figure S2 a, b. In the yeast study with THCz-2 (CAR- 8)^21^, it was shown that there was potent synergistic activity with fluconazole in a fluconazole-resistant strain of *Candida albicans* and the potent uncoupling activity of THCz-2 may contribute to decreased ATP synthesis, inhibiting ATP-powered efflux pumps.

Finally, we discuss some other possible mechanisms of action of SQ109 since as noted above, we found promising activity in an *in vivo* mice model of infection with an 80% survival rate.^20^ Here, there are two main points of interest. First, in a recent study of the activity of SQ109 against *M. tuberculosis* it was shown that SQ109 targets murine macrophages, converting cells to an M1 (pro-inflammatory) phenotype, resulting in activation of the mitogen-activated protein kinase (MAPK) pathway, a large (∼25x) increase in iNOS production, and downregulation of the M2- specific marker, arginase.^32^ The same effects were seen with infected peritoneal macrophages treated with SQ109, and SQ109-pretreated macrophages effectively killed *M. tuberculosis*. There are several other targets for SQ109 in *M. tuberculosis*, as shown in Figure 5a and as outlined above.

**Figure 5.**
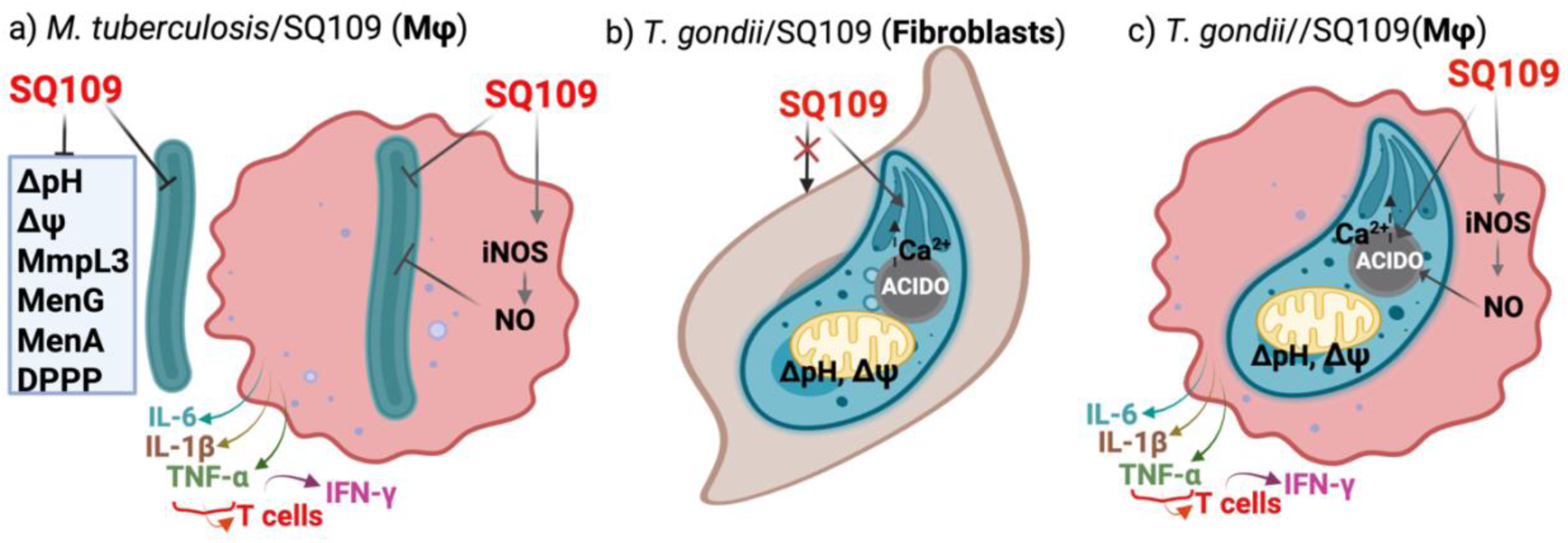
Summary of mechanism of action of SQ109 against *M. tuberculosis* and proposed mechanisms of action against *T. gondii* indicating the potential role of host cells. a) *M. tuberculosis*/SQ109/macrophage (Mφ) host indicating NO/iNOS role and innate immune response. b) *T. gondii*/SQ109/Fibroblast c) *T. gondii*/SQ109/macrophage.

Second, in previous work we reported that SQ109 has much greater activity (a ∼50-80x smaller IC_50_) against intracellular *Leishmania* spp. parasites (amastigotes), which are grown in macrophages, than against the extracellular (promastigote) forms^33–34^ but the differences in amastigote/epimastigote activity were far smaller in *Trypanosoma cruzi*, ∼ 5-20x, which are grown in fibroblasts (either Vero or LLC-MK_2_ cells). With two other cationic drugs, amiodarone and dronedarone, tested against *L. mexicana* and *T. cruzi*, there was a similar pattern of activity with a ∼100-200x increase in activity against the intracellular amastigotes in *L. mexicana* (in macrophages) but only ∼5x in *T. cruzi* (in fibroblasts). These results suggest that macrophage stimulation contributes to the potent activity seen against intracellular *Leishmania* parasites, but not in *T. cruzi*, at least *in vitro* in Vero or LLC-MK_2_ cells and by analogy, in *T. gondii* in animals. In the mice models of *M. tuberculosis* infection, it was also shown that SQ109 had major effects on the immune system that contributed to its efficacy. Specifically, there were large increases in the M1-specific markers IL-6, IL-1β, TNF-α and IFN-γ and given that *T. gondii* in animals also proliferate in macrophages,^35–37^ it is likely that there may be similar immune-related effects that contribute to its efficacy, in mice.

The mechanism of action of SQ109 for extracellular and intracellular *M. tuberculosis* (in macrophages) is shown in Figure 5a and that for SQ109 targeting *T. gondii* grown in hTERT (human telomerase reverse transcriptase-immortalized) fibroblasts is shown in Figure 5b. In the latter, uncoupling activity as well as targeting of acidocalcisomes is proposed, based on the observation of uncoupling activity in the IMV assay and the observation of effects on [Ca^2+^]_in_ in *T*. *cruzi*, *L. donovani* and *L. mexicana*. In Figure 5c we propose a model for *T. gondii* growth inhibition in macrophages and by extension *in vivo* activity, involving an innate immune response.

## CONCLUSIONS

The results we obtained are of interest for several reasons. First, we found that the carbazole anti-infectives THCz-1(CAR-20, **1**) and THCz-2(CAR-8, **2**) that inhibit bacterial and yeast growth also inhibited growth of the Apicomplexan parasite *T. gondii* with IC_50_ values of ∼2 μM. Second, we found that a phenythiazole **4** that likewise inhibits the growth of bacteria and yeasts also had potent activity against *T. gondii*, again with an IC_50_ value of ∼2 μM. Third, we investigated the activity of these compounds together with SQ109 and a series of 18 SQ109 analogs in an inverted membrane vesicle assay to determine their effects as protonophore uncouplers, finding in many cases potent activity, in the ∼1-4 μM range. Fourth, we found that there were correlations between the uncoupling effects and *T. gondii* growth inhibition, as well as with growth inhibition of the bacterium *M. smegmatis* and the yeast, *S. cerevisiae.* However, the correlations were more significant with the bacterium and yeast, suggesting that uncoupling effects play a larger role in these organisms than in *T. gondii*. We also propose that SQ109 (and its analogs) are likely to contribute to the killing of *T. gondii*, resident in macrophages, and *in vivo* via macrophage polarization. Overall, our results have broad general interest since they indicate that compounds that have previously been investigated as antibacterial and antifungal agents also have activity against *T. gondii*.

## METHODS

### General

All chemicals were of reagent grade and were used as received. Moisture-sensitive reactions were performed under an inert atmosphere (dry nitrogen) with dried solvents. Reactions were monitored by TLC using Merck silica gel 60 F-254 thin-layer plates. Flash column chromatography was carried out on Merck silica gel 60 (230–400 mesh). ^1^H NMR and ^13^C NMR spectra were recorded on Varian (Palo Alto, CA) Unity spectrometers at 400 and 500 MHz for ^1^H and at 100 and 125 MHz for ^13^C. Coupling constants (*J*) are reported in Hertz. High-resolution mass spectra (HRMS) were recorded in the University of Illinois Mass Spectrometry Laboratory. qNMR spectra were recorded using Varian (Palo Alto, CA) 500 MHz Unity spectrometers with 1,3,5- trimethoxybenzene (99.96% purity; Sigma-Aldrich TraceCERT, SKU: 74599-1G) as the internal total-spin-count quantitation standard; 60° pulse excitation, 60 s recycle delay, 65,536 data points,

1.0 Hz line-broadening due to exponential multiplication, and 16 accumulations. qNMR data were processed using Mnova NMR software (Mestrelab, Escondido, CA). All NMR spectra (including qNMR spectra) are provided in the supporting information. All tested compounds are >95% pure by qNMR analysis.

### Synthesis and Characterization of Compounds

#### 4-Cyclohexyl phenylhydrazine hydrochloride **(IM-1)**

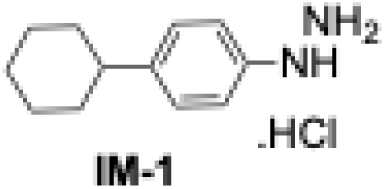

A solution of NaNO_2_ (476 mg, 6.90 mmol) in 5 mL of water was added dropwise to a mixture of 4-cyclohexyl aniline (1.00 g, 5.71 mmol) and concentrated HCl (10 mL) at 0 °C and stirred for 30 min after addition.^38^ Then, this RM was added dropwise into a stirred solution of SnCl_2_•H_2_O (3.84 g, 17.1 mmol) in 10 mL of concentrated HCl at RT. White precipitates formed upon stirring for 1 h were collected, washed thoroughly with water, and dried to afford **IM-1** (1.29 g, 92 %) as a white powder.^1^H NMR (400 MHz, DMSO-d_6_) *δ* 10.15 (s, 3H), 8.16 (s, 1H), 7.17 – 7.05 (m, 4H), 2.64 – 2.50 (m, 1H), 1.97 – 1.76 (m, 5H), 1.44 – 1.30 (m, 5H).

#### 6-Cyclohexyl-2,3,4,9-tetrahydro-1H-carbazol-1-one **(IM-2)**

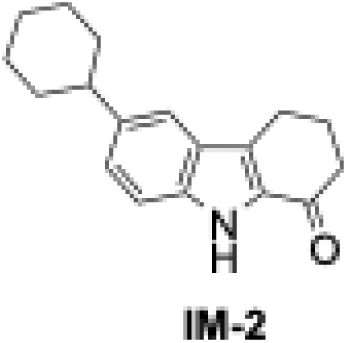

To a stirred solution of 1,2-cyclohexanedione (995 mg, 8.80 mmol) in concentrated HCl, (5 mL) and acetic acid (15 mL), was added dropwise 4-cyclohexyl phenylhydrazine **IM-1** (1.0 g, 4.41 mmol) solution in MeOH (15 mL).^22^ After addition, the reaction was heated at 60 °C for 1 h with continuous stirring. The RM was cooled to RT and subsequently kept in a refrigerator overnight. The bright yellow precipitate formed was filtered out, washed twice with MeOH, and dried *in vacuo* to afford **IM-2** (740 mg, 63 %) as a pale-yellow powder. ^1^H NMR (400 MHz, CDCl_3_) *δ* 8.73 (s, 1H), 7.46 (s, 1H), 7.33 (d, *J* = 8.4 Hz, 1H), 7.27 – 7.24 (m, 1H), 3.00 (t, *J* = 6.0 Hz, 2H), 2.65 (t, *J* = 6.4 Hz, 2H), 2.60 – 2.57 (m, 1H), 2.26 (p, *J* = 6.4 Hz, 2H), 1.94 – 1.85 (m, 4H), 1.77 (d, *J* = 12.8 Hz, 1H), 1.53 – 1.38 (m, 4H), 1.33 – 1.26 (m, 1H).

#### 6-Cyclohexyl-N-phenethyl-2,3,4,9-tetrahydro-1H-carbazol-1-amine **(1)**

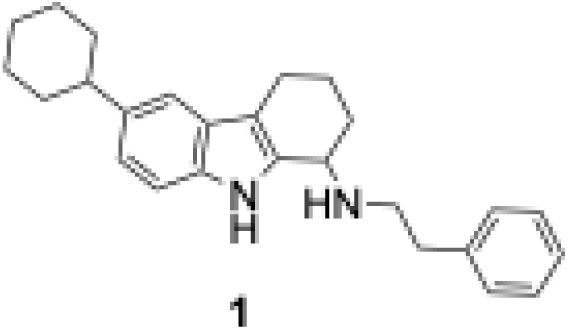

A mixture of 6-cyclohexyl-2,3,4,9-tetrahydro-*1H*-carbazol-1-one, **IM-2** (500 mg, 1.87 mmol), phenethylamine (362 mg, 3.01 mmol) and *p*TsOH (32.0 mg, 0.19 mmol) in dry toluene (10 mL) was heated at 140 °C for 18 h with a Dean-Stark trap under inert conditions. The solvent was removed *in vacuo* and the residue was dissolved in anhydrous MeOH. NaBH_4_ (120 mg) was added to this mixture at 0 °C and then heated to 80 °C for 4 h.^39^ The RM was quenched with H_2_O, extracted with DCM (2 × 20 mL), and the combined organic layer was washed with brine, dried over anhydrous Na_2_SO_4_, and then concentrated *in vacuo*. The crude extract was purified by silica gel flash chromatography (0–20% MeOH/DCM) to afford **1** (400 mg, 57 %) as a yellow semisolid. ^1^H NMR (500 MHz, CDCl_3_) *δ* 8.18 (s, 1H), 7.33 – 7.29 (m, 3H), 7.26 – 7.22 (m, 3H), 7.18 (d, *J* = 8.0 Hz, 1H), 7.02 (d, *J* = 8.5 Hz, 1H), 3.97 (t, *J* = 6.0 Hz, 1H), 3.05 – 2.99 (m, 1H), 2.97 – 2.92 (m, 1H), 2.90 – 2.85 (m, 1H), 2.80 – 2.74 (m, 1H), 2.72 – 2.64 (m, 2H), 2.62 – 2.55 (m, 1H), 2.19 – 2.13 (m, 1H), 2.03 – 1.96 (m, 1H), 1.96 – 1.90 (m, 2H), 1.89 – 1.83 (m, 2H), 1.80 – 1.74 (m, 2H), 1.66 – 1.59 (m, 1H), 1.55 – 1.38 (m, 4H), 1.33 – 1.27 (m, 1H); ^13^C NMR (125 MHz, CDCl_3_) δ 140.16, 139.24, 136.01, 134.43, 128.92, 128.61, 127.53, 126.37, 121.22, 115.64, 111.21, 110.65, 77.41, 77.16, 76.91, 52.42, 50.57, 47.13, 44.95, 37.01, 35.42, 35.41, 30.16, 27.31, 26.46, 21.87, 21.10; HRMS (TOF-ESI) m/z [M + H]^+^ calculated for [C_26_H_33_N_2_]^+^ 373.2644; found 373.2637.

#### 4-Biphenylhydrazine hydrochloride **(IM-3)**

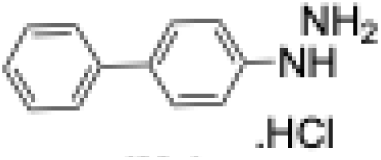

A solution of NaNO_2_ (980 mg, 14.2 mmol) in 5 mL of water was added dropwise to a mixture of 4-phenylaniline (2.00 g, 11.8 mmol) and concentrated HCl (10 mL) at 0 °C and stirred for 30 min after addition. Then, this RM was dropwise added into a stirred solution of SnCl_2_•H_2_O (8.00 g, 35.4 mmol) and 10 mL of concentrated HCl at RT.^38^ White precipitates formed upon stirring for 1 h were collected, washed thoroughly with water, and dried to afford **IM-3** (1.60 g, 73%) as a white crystalline powder. ^1^H NMR (500 MHz, DMSO–d_6_) *δ* 10.31 (s, 3H), 8.41 (s, 1H), 7.62 – 7.59 (m, 4H), 7.43 – 7.41 (m, 2H), 7.31– 7.28 (m, 1H), 7.08 (d, *J* = 8.5 Hz, 2H).

#### 6-Phenyl-2,3,4,9-tetrahydro-1H-carbazol-1-one **(IM-4)**

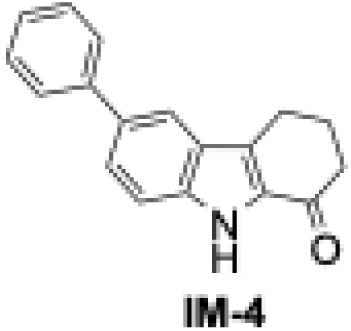

To a stirred solution of 1,2-cyclohexanedione (715 mg, 6.40 mmol) in concentrated HCl (3 mL) and acetic acid (9 mL) was added 4-biphenylhydrazine **IM-3** (700 mg, 3.20 mmol) solution in MeOH (10 mL) dropwise. After addition, the reaction was heated at 60 °C for 1 h with continuous stirring.^22^ The RM was cooled to RT and subsequently kept in a refrigerator overnight. The bright yellow precipitate formed was filtered out, washed with MeOH (2 × 5 mL), and dried *in vacuo* to afford **IM-4** (521 mg, 62%) as an orange powder. ^1^H NMR (500 MHz, DMSO) *δ* 10.75 (s, 1H), 7.04 (s, 1H), 6.79 – 6.73 (m, 3H), 6.55 – 6.42 (m, 4H), 2.16-2.05 (m, 2H), 1.67 – 1.60 (m, 4H).

#### N-phenethyl-6-phenyl-2,3,4,9-tetrahydro-1H-carbazol-1-amine **(2)**

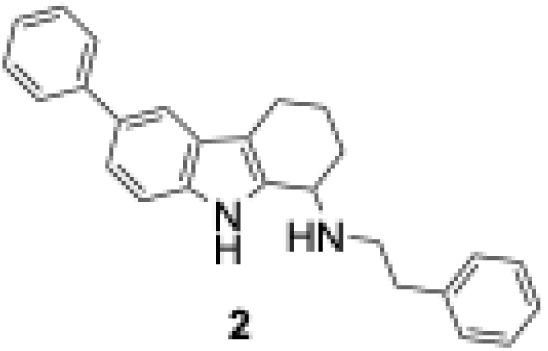

A mixture of ketone **IM-4** (150 mg, 0.58 mmol), phenethylamine (111 mg, 0.92 mmol), and *p*TsOH (10.0 mg, 0.187 mmol) in dry toluene (10 mL) was heated at 140 °C for 18 h with a Dean-Stark trap under inert conditions. The solvent was removed *in vacuo,* and the residue was dissolved in anhydrous MeOH. NaBH_4_ (65 mg, 1.72 mmol) was added to this mixture at 0 °C and then heated to 80 °C for 4 h.^39^ The RM was quenched with H_2_O, extracted twice with DCM (2 × 20 mL), and the combined organic layers were washed with brine, dried over anhydrous Na_2_SO_4_, and then concentrated *in vacuo*. The crude extract was purified by silica gel flash chromatography (0−20% MeOH/DCM) to afford **2** (115 mg, 54%) as a bright orange semisolid. ^1^H NMR (500 MHz, CDCl_3_) *δ* 8.16 (s, 1H), 7.65 – 7.63 (m, 3H), 7.43 – 7.36 (m, 3H), 7.33 – 7.28 (m, 4H), 7.24 – 7.21 (m, 3H), 3.97 (t, *J* = 7.0 Hz, 1H), 3.04 – 2.93 (m, 2H), 2.90 – 2.83 (m, 1H), 2.79 – 2.64 (m, 3H), 2.20 – 2.15 (m, 1H), 2.02 – 1.99 (m, 1H), 1.81 – 1.73 (m, 1H), 1.65 – 1.58 (m, 1H); ^13^C NMR (125 MHz, CDCl_3_) δ 142.96, 140.23, 136.89, 135.36, 132.68, 128.93, 128.82, 128.68, 128.60, 128.08, 127.43, 126.36, 126.22, 121.32, 116.91, 111.69, 111.13, 77.42, 77.16, 76.91, 52.43, 47.18, 37.06, 30.16, 21.87, 21.04.

#### 4-(Cyclohex-2-en-1-yloxy)-1,1’-biphenyl **(IM-5)**

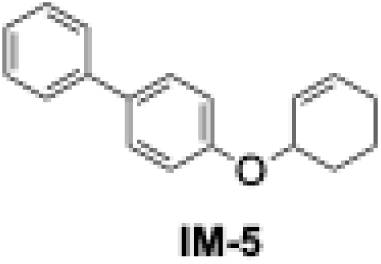

To a stirred solution of 4-phenylphenol (3.90 g, 22.5 mmol) in DMF (40 mL), was added bromocyclohexene (2.90 mL, 22.5 mmol) and K_2_CO_3_ (6.22 g, 45.0 mmol). The RM was stirred at RT for 24 h.^22^ The RM was further extracted with EtOAc (2 × 30 mL), rinsed with H_2_O (20 mL), followed by a brine wash, and dried over Na_2_SO_4_. After filtration, the solvent was evaporated *in vacuo,* and the crude mixture was purified by silica gel flash chromatography (0−40% EtOAc/hexane0 to afford **IM-5** (2.83 g, 48%) as a white solid. ^1^H NMR (500 MHz, CDCl_3_) *δ* 7.57 – 7.52 (m, 4H), 7.44 – 7.40 (m, 2H), 7.32 – 7.29 (m, 1H), 7.01 (d, *J* = 8.5 Hz, 2H), 6.02 – 5.99 (m, 1H), 5.93 – 5.90 (m, 1H), 4.85 (t, *J* = 7.0 Hz, 1H), 2.19 – 2.13 (m, 1H), 2.04 – 1.86 (m, 4H), 1.69 – 1.66 (m, 1H).

#### 1’’,2’’,3’’,6’’-Tetrahydro-[1,1’:3’,1’’-terphenyl]-4’-ol **(IM-6)**

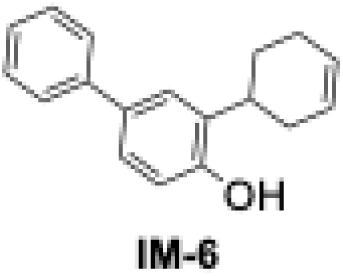

**IM-5** (2.80 g, 11.2 mmol) was dissolved in *N,N*-diethyl aniline (25 mL) and heated at 195 °C for 48 h. The RM was cooled to RT, and HCl (6 M, 30 mL) was added.^22^ The mixture was extracted with EtOAc (2 × 25 mL). The combined organic layer was dried over Na_2_SO_4_, filtered, and concentrated *in vacuo* to afford a brown oil, which after purification by silica gel flash chromatography (0−10% EtOAc/hexane) yielded **IM-6** (1.43 g, 51%) as a clear yellow oil. ^1^H NMR (400 MHz, CDCl_3_) *δ* 7.55 (d, *J* = 8.4 Hz, 2H), 7.43 – 7.40 (m, 2H), 7.36 – 7.34 (m, 2H), 7.32 – 7.28 (m, 1H), 6.88 (d, *J* = 8.4 Hz, 1H), 6.11 – 6.06 (m, 1H), 5.89 – 5.87 (m, 1H), 5.49 (s, 1H), 3.67 – 3.63 (m, 1H), 2.18 – 2.08 (m, 3H), 1.87 – 1.80 (m, 1H), 1.73 – 1.64 (m, 2H).

#### 8-Phenyl-1,2,3,4-tetrahydrodibenzo[b,d]furan-4-ol **(IM-7)**

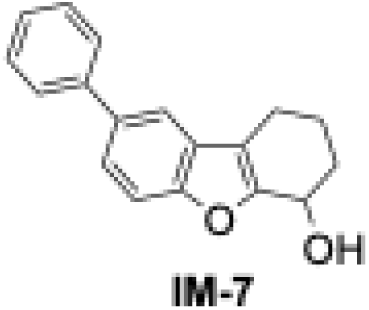

In an oven-dried sealed tube, **IM-6** (1.10 g, 4.43 mmol), *m*CPBA (915 mg, 5.32 mmol) in benzene (20 mL) were added. The RM was heated at 90 °C for 4 h, cooled to RT, washed with a saturated solution of NaHCO_3_ (3 × 15 mL), and dried over Na_2_SO_4_.^22^ After filtration, the solvent was evaporated *in vacuo* to afford a crude yellow oil which after purification by silica gel flash chromatography (EtOAc/hexane, 0−30%) yielded **IM-7** (670 mg, 57%) as a clear yellow oil. ^1^H NMR (500 MHz, CDCl_3_) *δ* 7.54 (d, *J* = 7.0 Hz, 2H), 7.43 – 7.40 (m, 2H), 7.38 – 7.35 (m, 1H), 7.31 – 7.29 (m, 2H), 6.90 (d, *J* = 8.0 Hz, 1H), 4.56 (t, *J* = 8.0 Hz, 1H), 3.74 – 3.68 (m, 2H), 2.18 – 2.14 (m, 1H), 1.90 – 1.83 (m, 2H), 1.71 – 1.68 (m, 1H).

#### 8-Phenyl-2,3-dihydrodibenzo[b,d]furan-4(1H)-one (**IM-8**)

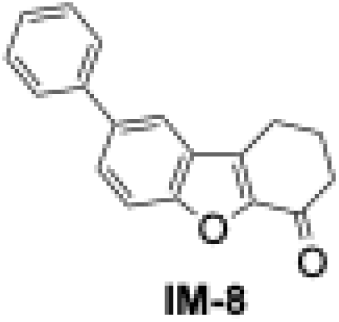

To a stirred solution of cyclic alcohol **IM-7** (500 mg, 1.87 mmol)) in xylene (15 mL), was added DDQ (1.28 g, 5.63 mmol), and the solution was heated in a sealed tube at 150 °C for 24 h.^22^ Then, the dark brown RM was filtered through Celite, and the residue was washed with EtOAc. The filtrate was concentrated *in vacuo* to afford a brown oil which was subsequently purified by silica gel flash chromatography (10−40% EtOAc/heptane) to afford **IM-8** (350 mg, 71%) as a brown solid. ^1^H NMR (500 MHz, CDCl_3_) *δ* 7.82 (d, *J* = 2.0 Hz, 1H), 7.74 – 7.72 (m, 1H), 7.64 – 7.61 (m, 3H), 7.49 – 7.46 (m, 2H), 7.39 – 7.36 (m, 1H), 3.05 (t, *J* = 6.0 Hz, 2H), 2.73 (t, *J* = 6.5 Hz, 2H), 2.32 (m, 2H).

#### N-phenethyl-8-phenyl-1,2,3,4-tetrahydrodibenzo[b, d]furan-4-amine **(3)**

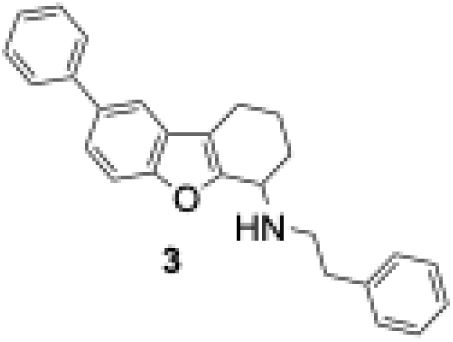

To a stirred solution of ketone **IM-8** (100 mg, 0.37 mmol) in DCE (5 mL), was added phenethylamine (285 μL, 2.27 mmol) sodium triacetoxyborohydride (479 mg, 2.27 mmol), and acetic acid (129 μL, 2.27 mmol). The RM was stirred at RT for 48 h, after that, saturated sodium bicarbonate (2 mL) was added.^22^ The resulting mixture was extracted with DCM (2 × 20 mL), and the combined organic layers were washed with brine, dried over anhydrous Na_2_SO_4_, and then concentrated *in vacuo*. The crude extract was purified by silica gel flash chromatography (0−10% MeOH/DCM to afford **3** (930 mg, 52%) as a clear yellow oil. ^1^H NMR (500 MHz, CDCl_3_) *δ* 7.61 – 7.59 (m, 3H), 7.46 – 7.41 (m, 4H), 7.33 – 7.28 (m, 3H), 7.25 – 7.19 (m, 3H), 3.97 (t, *J* = 5.5 Hz, 1H), 3.15 – 3.10 (m, 1H), 3.08 – 3.03 (m, 1H), 2.87 (t, *J* = 7.0 Hz, 2H), 2.68 – 2.56 (m, 2H), 2.08 – 2.03 (m, 1H), 1.96 – 1.91 (m, 1H), 1.88 – 1.76 (m, 2H); ^13^C NMR (125 MHz, CDCl_3_) δ 155.39, 154.16, 141.83, 139.96, 135.96, 128.74, 128.68, 128.45, 127.39, 126.73, 126.16, 112.32, 117.59, 114.94, 111.28, 51.51, 48.72, 36.74, 29.96, 20.66, 20.17; HRMS (TOF-ESI) m/z [M + H]^+^calculated for [C_26_H_26_NO]^+^ 368.2014; found 368.2021.

#### 4-Butylbenzamide **(IM-9)**

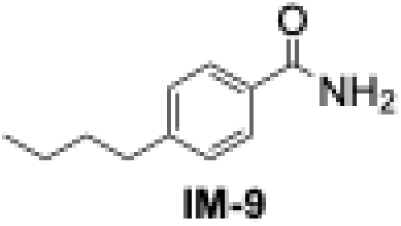

To a stirred solution of 4-butylbenzoic acid (1.78 g, 10.0 mmol) in DCM (20 mL) was added SOCl_2_ (2.38 g, 20.0 mmol) and refluxed for 3 h at RT. The RM was concentrated *in vacuo* and, NH_4_OH (20 mL) solution was added. The RM was stirred for an additional 3 h at RT.^24^ After completion (as monitored by TLC), the reaction was quenched with water and extracted with EtOAc (2 × 25 mL). The combined organic layers were washed with brine, dried over Na_2_SO_4,_ and concentrated *in vacuo*. The dried crude extract was recrystallized in methanol to afford **IM-9** (930 mg, 52%) as a white powder. ^1^H NMR (400 MHz, CDCl_3_) *δ* 7.76 – 7.72 (m, 2H), 7.27 – 7.24 (m, 2H), 6.11 (s, 2H), 2.69 – 2.64 (m, 2H), 1.65 – 1.58 (m, 2H), 1.36 (t, *J* = 7.2 Hz, 2H), 0.96 – 0.91 (m, 3H).

#### 4-Butylbenzothioamide **(IM-10)**

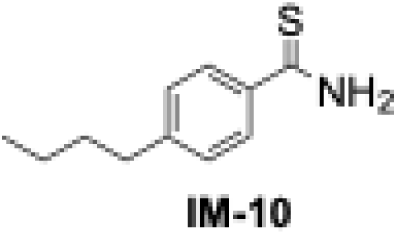

To a stirred solution of benzamide **IM-9** (700 mg, 3.93 mmol) in dry THF (15 mL), was added Lawesson’s reagent (2.38 g, 5.89 mmol) and stirred at RT for 12 h.^24^ The RM was concentrated under reduced pressure, and saturated NaHCO_3_ was added, followed by extraction with EtOAc (2 × 20 mL). The combined organic layers were washed with brine, dried over Na_2_SO_4,_ and concentrated *in vacuo*. The crude extract was purified by silica gel flash chromatography (0−12% EtOAc/Hexane) to afford **IM-10** (650 mg, 85%) as a yellow powder. ^1^H NMR (400 MHz, CDCl_3_) *δ* 7.85 – 7.79 (m, 3H), 7.32 – 7.20 (m, 3H), 2.71 – 2.63 (m, 2H), 1.67 – 1.58 (m, 2H), 1.42 – 1.34 (m, 2H), 0.99 – 0.91 (m, 3H).

#### 1-(2-(4-Butylphenyl)-4-methylthiazol-5-yl)ethan-1-one **(IM-11)**

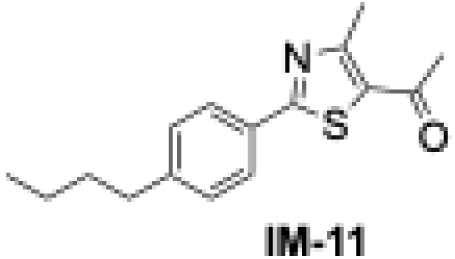

Benzothioamide **IM-10** (500 mg, 2.58 mmol), and 3-chloropentane-2,4-dione (483 mg, 3.61 mmol) were added to ethanol (15 mL) and the resulting mixture was refluxed for 24 h.^24^ Then, the solvent was evaporated *in vacuo*, and a brown residue was collected and purified by silica gel flash chromatography, using EtOAc/hexane (0−10%) to afford **IM-11** (502 mg, 51%) as a colorless oil. ^1^H NMR (400 MHz, CDCl_3_) *δ* 7.93 – 7.88 (m, 2H), 7.31 – 7.26 (m, 2H), 2.82 – 2.79 (m, 3H), 2.71 – 2.64 (m, 2H), 2.60 – 2.57 (m, 3H), 1.68 – 1.61 (m, 2H), 1.44 – 1.35 (m, 2H), 0.99 – 0.93 (m, 3H).

#### (E)-2-(1-(2-(4-butylphenyl)-4-methylthiazol-5-yl)ethylidene)hydrazine-1-carboximidamide **(4)**

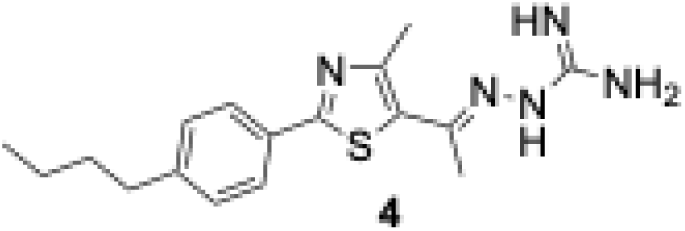

To a solution of ketone **IM-11** (450 mg, 1.65 mmol) in ethanol (20 mL), was added aminoguanidine hydrochloride (181 mg, 1.65 mmol) and a catalytic amount of LiCl (10.0 mg) and refluxed for 24 h.^24^ The solvent was evaporated *in vacuo,* and the obtained crude residue was purified by crystallization in 70% methanol followed by recrystallization in MeOH to afford **4** (230 mg, 42%) as a light-yellow powder. ^1^H NMR (500 MHz, DMSO– d_6_) *δ* 11.07 (s, 1H), 7.83 (d, *J* = 8.0 Hz, 2H), 7.62 (s, 3H), 7.33 (d, *J* = 8.0 Hz, 2H), 2.64 – 2.61 (m, 5H), 2.41 (s, 3H), 1.59 – 1.51 (m, 2H), 1.34 – 1.29 (m, 2H), 0.90 (t, *J* = 7.5 Hz, 3H); ^13^C NMR (125 MHz, DMSO – d_6_) *δ* 165.57, 156.35, 156.33, 152.84, 147.74, 145.85, 130.73, 130.63, 129.66, 126.46, 35.11, 33.28, 22.23, 18.66, 18.59, 14.23. HRMS (TOF-ESI) m/z [M + H]^+^ calculated for [C_17_H_24_N_5_S]^+^ 330.1752; found 330.1747.

#### (1r,3r,5r,7r)-2-Benzyladamantan-2-ol **(IM-12)**

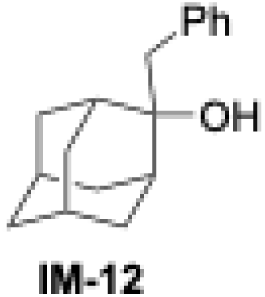

To a stirred solution of (1*r*,3*r*,5*r*,7*r*)-adamantan-2-one (1.00 g, 6.68 mmol) in dry THF (5 mL), was added dropwise BnMgCl (2.0 M in THF, 6.66 mL, 13.32 mmol at 0 °C and stirred overnight at RT.^31^ Then, saturated NH_4_Cl was poured into the RM and extracted in EtOAc (3 × 20 mL), and the combined organic layers were washed with brine, dried over Na_2_SO_4,_ and concentrated *in vacuo*. The crude extract was purified by silica gel flash chromatography (0−5% EtOAc/hexane) as eluents to afford **IM-12** (1.30 g, 81%) as a white powder. ^1^H NMR (400 MHz, CDCl_3_) *δ* 7.34 – 7.31 (m, 2H), 7.27 – 7.24 (m, 3H), 3.01 (s, 2H), 2.20 – 2.10 (m, 4H), 1.96 – 1.92 (m, 1H), 1.82 – 1.79 (m, 3H), 1.74 – 1.69 (m, 4H), 1.55 – 1.46 (m, 3H).

#### N-((1r,3r,5r,7r)-2-Benzyladamantan-2-yl)-2-chloroacetamide **(IM-13)**

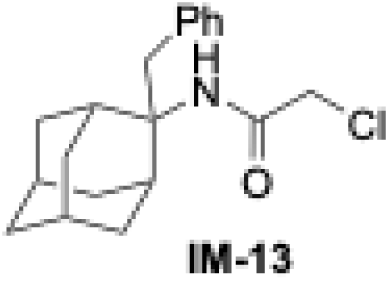

To a stirred solution of *(1r,3r,5r,7r)-*2-benzyladamantan-2-ol (1.00 g, 4.13 mmol) in AcOH (0.7 mL) and H_2_SO_4_, (0.7 mL), was added bromoacetonitrile (991 mg, 8.26 mmol).^31^ The RM was stirred at RT for 2 h, and the reaction completion was monitored by TLC. Then, The RM was poured into ice-cold water and neutralized using solid Na_2_CO_3_. The aqueous phase was extracted with EtOAc (2 × 20 mL), and the combined organic layer was washed with brine, dried over Na_2_SO_4,_ and evaporated *in vacuo.* The crude extract was purified by silica gel flash chromatography (0−5% EtOAc/hexane) to afford **IM-13** (900 mg, 68%) as a pale yellow powder. ^1^H NMR (400 MHz, CDCl_3_) *δ* 7.28 – 7.21 (m, 3H), 7.11 – 7.08 (m, 2H), 5.79 (s, 1H), 3.79 (s, 2H), 3.35 (s, 2H), 2.26 – 2.22 (m, 4H), 1.96 – 1.62 (m, 10H).

#### N-((1r,3r,5r,7r)-2-Benzyladamantan-2-yl)-2-(((E)-3,7-dimethylocta-2,6-dien-1-yl)amino) acetamide **(IM-14)**

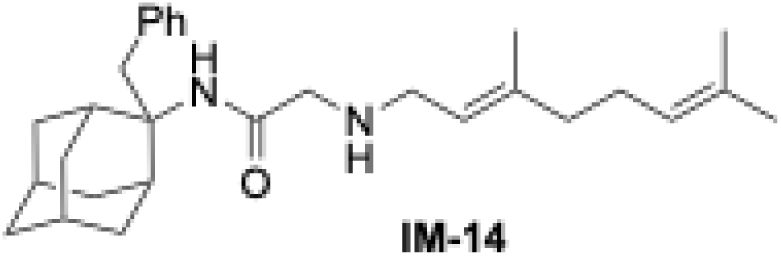

To a solution of chloroacetamide **IM-13** (500 mg, 1.38 mmol) and (*E*)-3,7-dimethylocta-2,6-dien- 1-amine (422 mg, 2.76 mmol) in DMF (5 mL), was added K_2_CO_3_ (476 mg, 3.45 mmol) and stirred at 70 °C for 20 h.^19^ The RM was cooled to RT, diluted with water (30 mL), and extracted with EtOAc (2 × 25 mL). Then, the combined organic layers were dried over Na_2_SO_4_, filtered, and concentrated *in vacuo.* The crude extract was purified by silica gel flash chromatography (10−40% EtOAc/hexane) to afford **IM-14** (500 mg, 83%) as a pale-yellow oil. ^1^H NMR (400 MHz, CDCl_3_) *δ* 7.22 (d, *J* = 7.6 Hz, 2H), 7.18 (d, *J* = 6.8 Hz, 1H), 7.11 (d, *J* = 7.2 Hz, 2H), 6.86 (s, 1H), 5.07 (t, 2H), 3.39 (s, 2H), 3.18 (s, 2H), 3.07 (d, *J* = 6.8 Hz, 2H), 2.30 – 2.24 (m, 4H), 2.06 – 2.02 (m, 2H), 1.99 – 1.90 (m, 6H), 1.82 – 1.79 (m, 2H), 1.75 (s, 3H), 1.67 (s, 3H), 1.64 (s, 1H), 1.61 – 1.59 (m, 4H), 1.55 (s, 3H).

#### N^1^-((1r,3r,5r,7r)-2-Benzyladamantan-2-yl)-N^2^-((E)-3,7-dimethylocta-2,6-dien-1-yl)ethane-1,2- diamine **(6)**

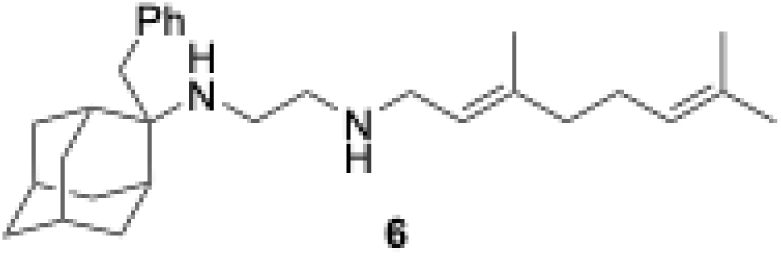

To a solution of acetamide **IM-14** (500 mg, 1.15 mmol) in 5 mL Et_2_O, was added LiAlH_4_ (218 mg, 5.74 mmol), and RM was refluxed for 48 h under N_2_ atmosphere.^19^ The RM was cooled to RT and quenched by adding 0.5 mL of NH_4_OH solution, filtered through celite, and concentrated *in vacuo.* The crude product was purified by silica gel flash chromatography (DCM/MeOH/NH_4_OH; 90:9:1) as an eluent to afford **6** (152 mg, 31%) as a yellow oil and converted into the HCl salt (an off-white powder) using a 2 M HCl solution in diethyl ether. Free base; ^1^H NMR (500 MHz, CDCl_3_) *δ* 7.37 – 7.34 (m, 2H), 7.29 – 7.24 (m, 3H), 5.34 (t, *J* = 7.0 Hz, 1H), 5.18 (d, *J* = 7.0 Hz, 1H), 3.33 (d, *J* = 7.0 Hz, 2H), 3.07 (s, 2H), 2.88 (d, *J* = 6.0 Hz, 2H), 2.78 (t, *J* = 6.0 Hz, 2H), 2.34 (d, *J* = 12.5 Hz, 2H), 2.27 (d, *J* = 13.0 Hz, 2H), 2.17 (q, *J* = 7.5 Hz, 2H), 2.11 – 2.08 (m, 2H), 2.03 (s, 1H), 1.88 (s, 1H), 1.82 – 1.76 (m, 9H), 1.72-168 (m, 8H), 1.51 (d, *J* = 12 Hz, 2H); ^13^C NMR (125 MHz, CDCl_3_) δ 139.36, 138.41, 132.25, 130.99, 128.76, 126.68, 124.90, 123.53, 78.06, 77.80, 77.55, 59.12, 50.73, 48.04, 40.39, 40.24, 39.83, 37.43, 34.83, 34.47, 33.46, 28.84, 28.35, 27.28, 26.45, 18.44, 17.09; HRMS (TOF-ESI) m/z [M + H]^+^ calculated for [C_29_H_45_N_2_]^+^ 421.3583; found 421.3570.

#### (1r,3r,5r,7r)-2-Butyladamantan-2-ol **(IM-15)**

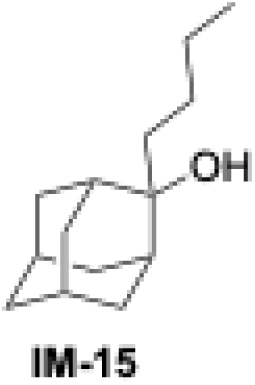

To a stirred solution of (1*r*,3*r*,5*r*,7*r*)-adamantan-2-one (500 mg, 3.34 mmol) in dry THF (5 mL), was added dropwise *n*BuLi (1.6 M in hexane, 6.25 mL, 10.0 mmol) at 0 °C ^24^ and stirred overnight at RT. Then, saturated NH_4_Cl was poured into the RM and extracted in EtOAc (3 × 20 mL). The combined organic layer was washed with brine, dried over Na_2_SO_4,_ and concentrated *in vacuo*. The crude extract was purified by silica gel flash chromatography (0−5% EtOAc/hexane) as systems solvents to afford **IM-15** (601 mg, 83%) as a white powder. ^1^H NMR (500 MHz, CDCl_3_) *δ* 2.17 (d, *J* = 10.0 Hz, 2H), 1.86 – 1.81 (m, 4H), 1.72 – 1.68 (m, 6H), 1.66 – 1.63 (m, 2H), 1.56 – 1.53 (m, 3H), 1.35 – 1.30 (m, 4H), 0.92 (t, *J* = 6.5 Hz, 3H).

#### N-((1r,3r,5r,7r)-2-Butyladamantan-2-yl)-2-chloroacetamide **(IM-16).**

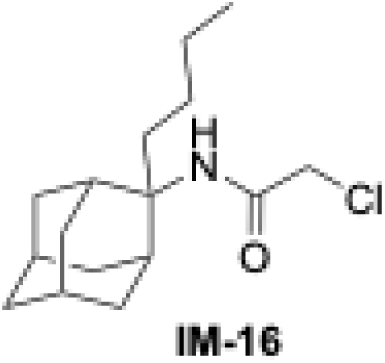

To a stirred solution of 2-adamantanol **IM-15** (1.00 g, 4.80 mmol) in AcOH (0.7 mL) and H_2_SO_4_ (0.7 mL), was added bromoacetonitrile (460 mg, 3.84 mmol).^24^ The RM was stirred at RT for 2 h, and the reaction completion was monitored by TLC. Then, The RM was poured into ice-cold water and neutralized using solid Na_2_CO_3_. The aqueous phase was extracted with EtOAc (2 × 20 mL), and the combined organic layer was washed with brine, dried over Na_2_SO_4,_ and evaporated *in vacuo.* The crude extract was purified by silica gel flash chromatography (0−5% EtOAc/hexane) to afford **IM-16** (860 mg, 63%) as a white powder.

#### N-((1r,3r,5r,7r)-2-Butyladamantan-2-yl)-2-(((E)-3,7-dimethylocta-2,6-dien-1-yl)amino) acetamide **(IM-17)**

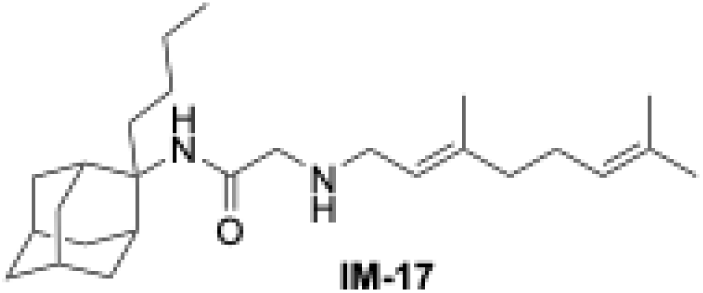

To a solution of chloroacetamide **IM-16** (600 mg, 1.83 mmol) and (*E*)-3,7-dimethylocta-2,6-dien- 1-amine (419 mg, 2.75 mmol) in DMF (5 mL), was added K_2_CO_3_ (632 mg, 4.58 mmol), and RM was stirred at 70 °C for 20 h.^19^ The RM was cooled to RT, diluted with water (30 mL), and extracted with EtOAc (2 × 25 mL). The combined organic layer was dried over Na_2_SO_4_, filtered, and concentrated *in vacuo.* The crude extract was purified by silica gel flash chromatography (10−40% EtOAc/hexane) to afford **IM-17** (480 mg, 65%) as a pale-yellow oil. ^1^H NMR (500 MHz, CDCl_3_) *δ* 7.29 – 7.27 (m, 1H), 5.23 (t, *J* = 7.0 Hz, 1H), 5.09 (t, *J* = 7.0 Hz, 1H), 3.33 – 3.19 (m, 4H), 2.30 – 2.27 (m, 2H), 2.12 – 1.94 (m, 12H), 1.84 (s, 3H), 1.72 – 1.60 (m, 16H), 1.33 – 1.29 (m, 2H), 1.20 – 1.12 (m, 2H).

#### N^1^-((1r,3r,5r,7r)-2-Butyladamantan-2-yl)-N^2^-((E)-3,7-dimethylocta-2,6-dien-1-yl)ethane-1,2- diamine (7)

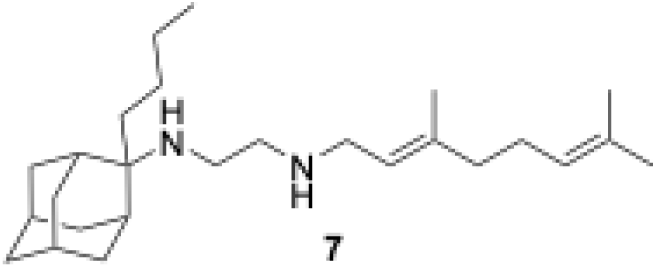

To a solution of acetamide **IM-17** (100 mg, 0.250 mmol) in Et_2_O (3 mL), was added LiAlH_4_ (47.0 mg, 1.25 mmol), and RM was refluxed for 48 h under N_2_ atmosphere.^19^ The RM was cooled to RT and quenched by adding 0.5 mL of NH_4_OH solution, filtered through celite, and concentrated *in vacuo.* The crude product was purified by silica gel flash chromatography (DCM/MeOH/NH_4_OH, 90:9:1) to afford **7** (42 mg, 43%) as a yellow oil and was converted into the HCl salt (an off-white solid) using a 2 M HCl solution in diethyl ether. Free base; ^1^H NMR (500 MHz, CDCl_3_) *δ* 5.25 (t, *J* = 6.0 Hz, 1H), 5.09 (t, *J* = 6.5 Hz, 1H), 3.24 – 3.22 (m, 2H), 2.70 – 2.67 (m, 2H), 2.54 – 2.50 (m, 2H), 2.18 – 2.15 (m, 2H), 2.10 – 2.04 (m, 2H), 2.02 – 1.99 (m, 2H), 1.92 (d, *J* = 13.0 Hz, 2H), 1.81 – 1.77 (m, 2H), 1.68 – 1.56 (m, 21H), 1.45 (d, *J* = 13.0 Hz, 2H), 1.31z – 1.24 (m, 3H), 1.17 – 1.13 (m, 2H); ^13^C NMR (125 MHz, CDCl_3_) δ 137.65, 131.61, 124.29, 123.15, 77.41, 77.16, 76.91, 56.97, 50.32, 47.26, 39.77, 39.43, 39.27, 37.44, 34.20, 33.73, 32.76, 31.52, 28.02, 27.88, 26.64, 25.82, 24.14, 23.60, 17.80, 16.42; HRMS (TOF-ESI) m/z [M + H]^+^ calculated for [C_26_H_47_N_2_]^+^ 387.3739; found 387.3720.

#### (1r,3r,5r,7r)-2-phenyladamantan-2-ol **(IM-18)**

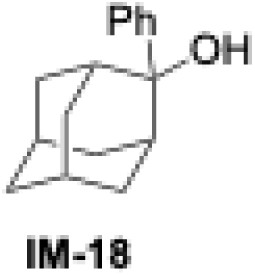

To a stirred solution of (1*r*,3*r*,5*r*,7*r*)-adamantan-2-one (1.00 g, 6.68 mmol) in dry THF (4 mL), was added dropwise PhMgBr (3.0 M in hexane, 4.45 mL, 13.4 mmol) was added dropwise at 0 °C and the mixture was the stirred overnight at RT.^31^ To the obtained RM was added saturated NH_4_Cl followed by extraction with EtOAc (3 × 20 mL). The combined organic layer was washed with brine, dried over Na_2_SO_4,_ and concentrated *in vacuo*. The crude extract was purified by silica gel flash chromatography (0−2% EtOAc/hexane) to afford **IM-18** (1.30 g, 85%) as a white powder. ^1^H NMR (400 MHz, CDCl_3_) *δ* 7.55 (d, *J* = 8.0 Hz, 2H), 7.41 – 7.35 (m, 2H), 7.29 (d, *J* = 8.0 Hz, 1H), 2.58 (s, 2H), 2.41 (d, *J* = 12.4 Hz, 2H), 1.91 (s, 1H), 1.75 – 1.67 (m, 10H).

#### 2-chloro-N-((1r,3r,5r,7r)-2-phenyladamantan-2-yl) acetamide **(IM-19)**

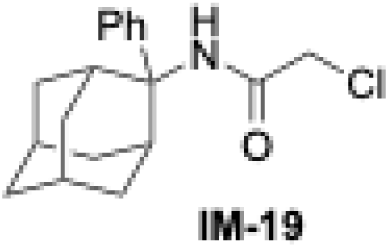

To a stirred solution of 2-adamantanol **IM-18** (1.00 g, 4.38 mmol) in AcOH (0.7 mL) and H_2_SO_4_ (0.7 mL), was added bromoacetonitrile (1.05 g, 8.77 mmol) and the resulting mixture was stirred at RT for 4 h.^31^ The reaction progress was monitored by TLC. The RM was poured into ice-cold water and neutralized using solid Na_2_CO_3_. The aqueous phase was extracted twice with EtOAc (3 × 20 mL), and the combined organic layer was washed with brine, dried over Na_2_SO_4,_ and evaporated *in vacuo.* The crude extract was purified by silica gel flash chromatography (0−10% EtOAc/hexane) to afford **IM-19** (1.02 g, 76%) as a white powder. ^1^H NMR (400 MHz, CDCl_3_) δ 7.57 (d, *J* = 7.2 Hz, 2H), 7.36 – 7.32 (m, 2H), 7.26 – 7.23 (m, 1H), 6.51 (s, 1H), 3.67 (s, 2H), 2.14 (d, *J* = 13.2 Hz, 2H), 1.95 – 1.60 (m, 12H).

#### 2-(((E)-3,7-dimethylocta-2,6-dien-1-yl)amino)-N-((1r,3r,5r,7r)-2-phenyladamantan-2-yl) acetamide **(IM-20)**

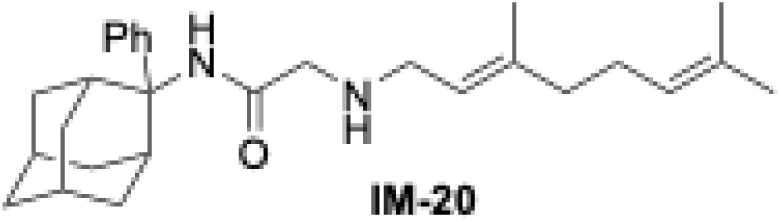

To a solution of chloroacetamide **IM-19** (500 mg, 1.43 mmol) and (*E*)-3,7-dimethylocta-2,6-dien- 1-amine (437 mg, 2.86 mmol) in DMF (10 mL), was added K_2_CO_3_ (495 mg, 3.59 mmol) and RM was stirred at 70 °C for 20 h.^19^ The RM was cooled to RT, and diluted with water (30 mL) and extracted with EtOAc (2 ×25 mL). The combined organic layer was dried over Na_2_SO_4_, filtered, and concentrated *in vacuo.* The crude extract was purified by silica gel flash chromatography (10−40% EtOAc/hexane) to afford **IM-20** (370 mg, 63%) as a pale-yellow oil. ^1^H NMR (400 MHz, CDCl_3_) *δ* 7.59 (d, *J* = 8.0 Hz, 2H), 7.42 (s, 1H), 7.34 – 7.30 (m, 2H), 7.21 – 7.17 (m, 1H), 5.14 (d, *J* = 7.2 Hz, 1H), 5.07 (d, *J* = 7.2 Hz, 1H), 3.13 (s, 2H), 3.07 (d, *J* = 7.2 Hz, 2H), 2.99 (s, 2H), 2.15 (d, *J* = 13.2 Hz, 2H), 2.08 – 2.04 (m, 2H), 2.00 – 1.96 (m, 2H), 1.91 (s, 1H), 1.83 – 1.80 (m, 4H), 1.73 (s, 3H), 1.68 – 1.66 (m, 5H), 1.60 (s, 3H), 1.55 (s, 3H).

#### 2-(((E)-3,7-dimethylocta-2,6-dien-1-yl)amino)-N-((1r,3r,5r,7r)-2-phenyladamantan-2-yl) acetamide **(8)**

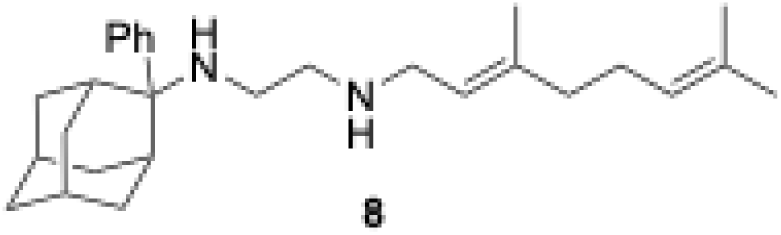

To a solution of acetamide **IM-20** (500 mg, 1.19 mmol) in dry DCM (10 mL) at 0 °C, was added TMSCl (192 mg, 1.78 mmol), and the reaction was stirred for 15 min under N_2_ atmosphere. Then, a suspension of LiAlH_4_ (90 mg, 2.38 mmol) in THF (1 mL) was added into RM at –10 °C, and stirring continued for an additional 3 h at −10 to 0 °C.^31^ The reaction was quenched with 10% NaOH solution, the inorganic precipitate was filtered off, and the organic layer was separated. The aqueous phase was extracted with DCM (2 × 20 mL), and the combined organic layer was dried over Na_2_SO_4_, filtered, and concentrated *in vacuo.* The crude product was purified by silica gel flash chromatography (DCM:MeOH:NH_4_OH; 90:8: 2) to afford **8** (158 mg, 32%) as a colorless oil and converted into the HCl salt (an off-white solid) using a 2 M HCl solution in diethyl ether. Free base; ^1^H NMR (500 MHz, CDCl_3_) *δ* 7.36 – 7.30 (m, 4H), 7.19 – 7.16 (m, 1H), 5.16 (t, *J* = 7.0 Hz, 1H), 5.08 (t, *J* = 6.5 Hz, 1H), 3.05 (d, *J* = 6.5 Hz, 2H), 2.49 – 2.46 (m, 4H), 2.42 (d, *J* = 12.5 Hz, 2H), 2.21 (t, *J* = 6.0 Hz, 2H), 2.08– 2.04 (m, 2H), 1.99 – 1.96 (m, 2H), 1.89 – 1.88 (m, 1H), 1.75 – 1.73 (m, 3H), 1.68 – 1.64 (m, 10H), 1.59 (s, 3H), 1.57 (s, 3H); ^13^C NMR (125 MHz, CDCl_3_) *δ* 144.15, 131.98, 128.54, 126.64, 126.59, 124.16, 123.88, 119.63, 77.42, 77.16, 76.91, 61.48, 47.16, 45.18, 39.72, 38.20, 34.15, 32.97, 32.78, 27.98, 27.00, 26.49, 25.85, 17.86, 16.52; HRMS (TOF-ESI) m/z [M + H]^+^ calculated for [C_28_H_43_N_2_]^+^ 407.3426; found 407.3414.

#### N-((3s,5s,7s)-adamantan-1-yl)-2-chloroacetamide **(IM-21)**

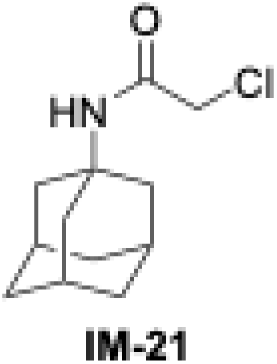

To a stirred solution of 1-adamantylamine (2.00 g, 13.2 mmol) in DCM (15 mL), was added an aqueous solution (10 mL) of K_2_CO_3_ (2.70 g, 3.59 mmol) at 0 °C. To the resulting mixture was added dropwise solution of chloroacetyl chloride (1.63 g, 14.5 mmol) dissolved in DCM (10 mL) over 30 min and stirred for 24 h. The RM was extracted with DCM (2 × 20 mL), and the combined organic layers were washed with Na_2_CO_3_ followed by 3% v/v HCl and brine, dried over Na_2_SO_4_, filtered, and concentrated *in vacuo.* The crude product was purified by silica gel flash chromatography (0−20 % EtOAc/hexane) to afford **IM-21** (1.84 g, 61%) as an off-white powder.

^1^H NMR (400 MHz, CDCl_3_) *δ* 6.22 (s, 1H), 3.92 (s, 2H), 2.09 (s, 3H), 2.02 – 2.01f (m, 6H), 1.69 (d, *J* = 3.2 Hz, 6H).

#### N-((3s,5s,7s)-adamantan-1-yl)-2-(phenethylamino) acetamide **(IM-22)**

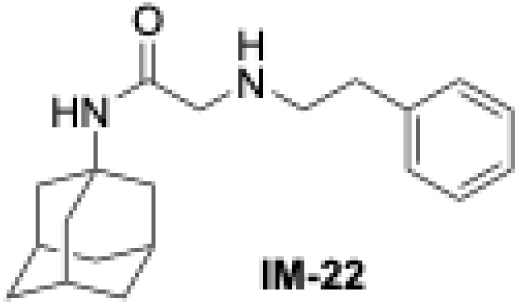

To a solution of chloroacetamide **IM-21** (250 mg, 1.10 mmol) and phenethylamine (200 mg, 1.65 mmol) in DMF (5 mL), was added K_2_CO_3_ (380 mg, 2.75 mmol), and stirred at 70 °C for 24 h.^19^ The RM was cooled to RT, diluted with water (30 mL), and extracted with EtOAc (2 × 25 mL). The combined organic layers were washed with brine, dried over Na_2_SO_4_, filtered, and concentrated *in vacuo.* The crude extract was purified by silica gel flash chromatography (0−5% EtOAc/hexane) to afford **IM-22** (280 mg, 81%) as a yellow oil. ^1^H NMR (400 MHz, CDCl_3_) *δ* 7.37 – 7.33 (m, 2H), 7.28 – 7.23 (m, 3H), 6.94 (s, 1H), 3.16 (s, 2H), 2.90 (t, *J* = 6.8 Hz, 2H), 2.82 – 2.78 (m, 2H), 2.09 (s, 3H), 2.01 – 1.93 (m, 6H), 1.92 (s, 1H), 1.73 – 1.68 (m, 6H).

#### N^1^-((3s,5s,7s)-adamantan-1-yl)-N^2^-phenethylethane-1,2-diamine **(9)**

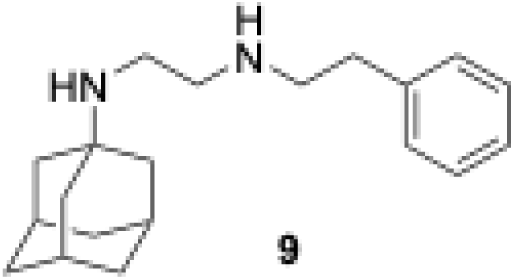

To a solution of acetamide **IM-22** (250 mg, 0.80 mmol) in dry DCM (5 mL) at 0 °C, was added TMSCl (173 mg, 1.60 mmol) and stirred for 15 min under N_2_ atmosphere. Then, a suspension of LiAlH_4_ (46 mg, 1.20 mmol) in THF (1 mL) was added to the RM at –10 °C, and stirring continued for another 3 h at –10 to 0 °C.^31^ The reaction was quenched with 10% NaOH solution, the inorganic precipitate was filtered off, and the organic layer was separated. The aqueous phase was extracted with DCM (2 × 20 mL) and the combined organic layer was dried over Na_2_SO_4_, filtered, and concentrated *in vacuo.* The crude product was purified by silica gel flash chromatography (DCM/MeOH/NH_4_OH; 90:8: 2) to afford **9** (110 mg, 46%) as a colorless oil and converted into the HCl salt (an off-white solid) using a 2 M HCl solution in diethyl ether. Free base; ^1^H NMR (500 MHz, CDCl_3_) *δ* 7.31 – 7.27 (m, 2H), 7.23 – 7.19 (m, 3H), 2.86 (t, *J* = 7.0 Hz, 2H), 2.80 (t, *J* = 6.5 Hz, 2H), 2.74 (t, *J* = 5.5 Hz, 2H), 2.70 – 2.68 (m, 2H), 2.06 – 2.05 (m, 3H), 1.93 (s, 2H), 1.67 – 1.58 (m, 12H); ^13^C NMR (125 MHz, CDCl_3_) δ 140.21, 128.83, 128.49, 126.17, 51.02, 50.68, 49.91, 42.55, 39.71, 36.72, 36.51, 29.58. HRMS (TOF-ESI) m/z [M + H]^+^ calculated for [C_20_H_31_N_2_]^+^ 299.2487; found 299.2482.

#### N-((3s,5s,7s)-adamantan-1-yl)-2-(((E)-3,7-dimethylocta-2,6-dien-1-yl)amino)acetamide **(IM-23)**

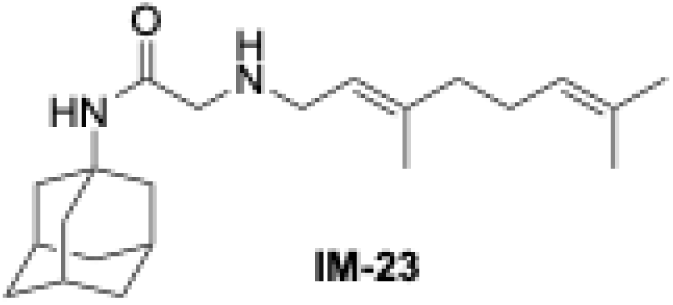

To a solution of chloroacetamide **IM-21** (400 mg, 1.76 mmol) and (*E*)-3,7-dimethylocta-2,6-dien- 1-amine (404 mg, 2.64 mmol) in DMF (5 mL), was added K_2_CO_3_ (607 mg, 4.40 mmol), and stirred at 70 °C for 20 h.^19^ The RM was cooled to RT, diluted with water (30 mL), and extracted with EtOAc (2 × 25 mL). The combined organic layer was washed with brine, dried over Na_2_SO_4_, filtered, and concentrated *in vacuo.* The crude extract was purified by silica gel flash chromatography (0−5 % MeOH/DCM) to afford **IM-23** (400 mg, 66%) as a yellow oil. ^1^H NMR (400 MHz, CDCl_3_) *δ* 6.97 (s, 1H), 5.19 (t, *J* = 7.2 Hz, 1H), 5.08 (t, *J* = 6.8 Hz, 1H), 3.17 (d, *J* = 6.8 Hz, 2H), 3.11 (s, 2H), 2.11 – 2.06 (m, 5H), 2.02 – 1.97 (m, 8H), 1.72 – 1.68 (m, 10H), 1.63 (s, 3H), 1.60 (s, 3H).

#### N^1^-((3s,5s,7s)-adamantan-1-yl)-N^2^-((E)-3,7-dimethylocta-2,6-dien-1-yl)ethane-1,2-diamine **(10)**

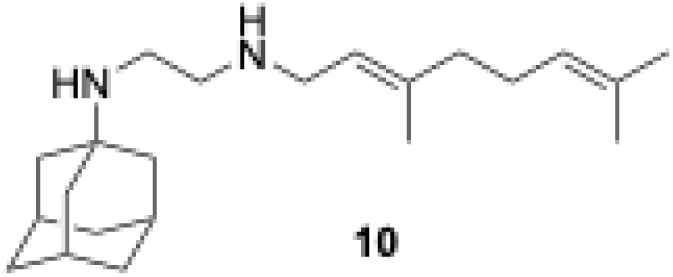

To a solution of acetamide **IM-23** (500 mg, 1.45 mmol) in dry DCM (10 mL) at 0 °C, was added TMSCl (235 mg, 2.18 mmol) and stirred for 15 min under N_2_ atmosphere. A suspension of LiAlH_4_ (110 mg, 2.90 mmol) in THF (1 mL) was added to the RM at −10 °C, and stirring continued for another 3 h at −10 to 0 °C.^31^ The reaction was quenched with 10% NaOH solution, the inorganic precipitate was filtered off, and the organic layer was separated. The aqueous phase was extracted with DCM (2 × 20 mL) and the combined organic layer was dried over Na_2_SO_4_, filtered, and concentrated *in vacuo.* The crude product was purified by silica gel flash chromatography (DCM:MeOH:NH_4_OH; 90:8:2) to afford **10** (152 mg, 31%) as a colorless oil and converted into the HCl salt (white solid) using a 2 M HCl solution in diethyl ether. Free base; ^1^H NMR (500 MHz, CDCl_3_) *δ* 5.23 (t, *J* = 7.0 Hz, 1H), 5.07 (d, *J* = 7.0 Hz, 1H), 3.20 (d, *J* = 7.0 Hz, 2H), 2.70 (s, 4H), 2.07 – 2.04 (m, 5H), 2.00 – 1.97 (m, 2H), 1.91 (s, 2H), 1.65 – 1.57 (m, 21H); ^13^C NMR (125 MHz, CDCl_3_) δ 137.87, 131.61, 124.22, 122.81, 77.42, 77.16, 76.91, 50.71, 49.78, 47.09, 42.69, 39.93, 39.74, 36.79, 29.65, 26.60, 25.80, 17.79, 16.42; HRMS (TOF-ESI) m/z [M + H]^+^ calculated for [C_22_H_39_N_2_]^+^ 331.3113; found 331.3117.

#### N-((3s,5s,7s)-adamantan-1-yl)-3-bromopropanamide **(IM-24)**

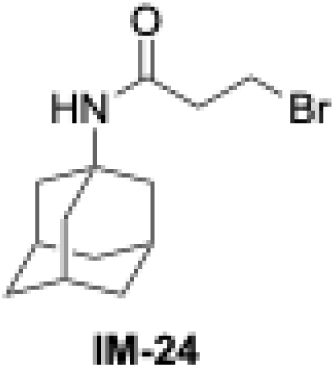

To a stirred solution of 1-adamantylamine (250 mg, 1.66 mmol) in DCM (5 mL), was added an aqueous solution (5 mL) of K_2_CO_3_ (342 mg, 2.48 mmol) at 0 °C. After this, 3-bromopropionyl chloride (312 mg, 1.83 mmol) dissolved in DCM (5 mL) was added dropwise over 30 min and the RM was stirred for 24 h at 30 °C.^19^ The resulting mixture was extracted with DCM (2 × 20 mL), and the combined organic layers were washed with NaHCO_3_ followed by 3% v/v HCl and brine, dried over Na_2_SO_4_, filtered, and concentrated *in vacuo.* The crude product was purified by silica gel flash chromatography (0−5% MeOH/DCM) to afford **IM-24** (280 mg, 60%) as an off-white powder. ^1^H NMR (400 MHz, CDCl_3_) *δ* 5.21 (s, 1H), 3.61 (t, *J* = 6.8 Hz, 2H), 2.65 (t, *J* = 6.8 Hz, 2H), 2.08 (s, 3H), 2.01 – 1.98 (m, 6H), 1.74 – 1.63 (m, 6H).

#### N-((3s,5s,7s)-adamantan-1-yl)-3-(((E)-3,7-dimethylocta-2,6-dien-1-yl)amino) propanamide **(IM-25)**

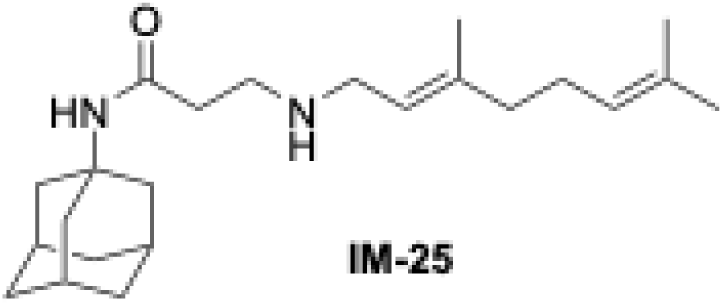

To a solution of 3-bromopropanamide **IM-24** (250 mg, 0.871 mmol) and (*E*)-3,7-dimethylocta- 2,6-dien-1-amine (200 mg, 1.31 mmol) in DMF (5 mL), was added K_2_CO_3_ (300 mg, 2.19 mmol), and the RM was stirred at 70 °C for 20 h.^19^ The RM was cooled to RT, diluted with water (30 mL), and extracted with EtOAc (2 × 25 mL). The combined organic layers were washed with brine, dried over Na_2_SO_4_, filtered, and concentrated *in vacuo.* The crude extract was purified by silica gel flash chromatography (0−5% EtOAc/hexane) to afford **IM-25** (165 mg, 53%) as a yellow oil. ^1^H NMR (400 MHz, CDCl_3_) *δ* 6.99 (s, 1H), 5.24 (t, *J* = 7.2 Hz, 1H), 5.08 (d, *J* = 7.2 Hz, 1H), 3.26 (d, *J* = 6.8 Hz, 2H), 2.85 (d, *J* = 6.4 Hz, 2H), 2.32 (d, *J* = 5.6 Hz, 2H), 2.12 – 1.98 (m, 13H), 1.71 – 1.65 (m, 12H), 1.59 (s, 3H).

#### N^1^-((3s,5s,7s)-adamantan-1-yl)-N^3^-((E)-3,7-dimethylocta-2,6-dien-1-yl)propane-1,3-diamine**(11)**

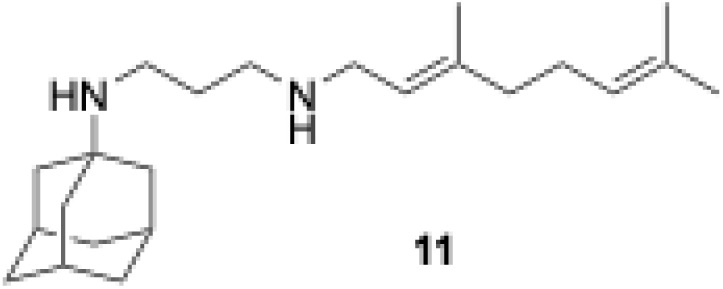

To a solution of propanamide **IM-25** (140 mg, 0.392 mmol) in dry DCM (10 mL) at 0 °C, was added TMSCl (84.0 mg, 0.781 mmol) and stirred for 15 min under N_2_ atmosphere. A suspension of LiAlH_4_ (37.0 mg, 0.971 mmol) in THF (1 mL) was added into RM –10 °C, and stirring continued for another 3 h at –10 to 0 °C.^31^ The reaction was quenched with 10% NaOH solution, the inorganic precipitate was filtered off, and the organic layer was separated. The aqueous phase was extracted with DCM (2 × 20 mL), and the combined organic layer was dried over Na_2_SO_4_, filtered, and concentrated *in vacuo*. The crude product was purified by silica gel flash chromatography (DCM/MeOH/NH_4_OH; 90:8:2) to afford **11** (100 mg, 74%) as a colorless oil and converted into the HCl salt (an off-white solid) using a 2 M HCl solution in diethyl ether. Free base; ^1^H NMR (500 MHz, CDCl_3_) *δ* 5.19 (d, *J* = 7.0 Hz, 1H), 5.03 (d, *J* = 7.0 Hz, 1H), 3.17 (d, *J* = 7.0 Hz, 2H), 2.65 – 2.63 (m, 4H), 2.51 (s, 1H), 2.05 – 2.01 (m, 5H), 1.97 – 1.94 (m, 2H), 1.67 – 1.54 (m, 24H); ^13^C NMR (125 MHz, CDCl_3_) δ 138.15, 131.64, 124.16, 122.48, 77.42, 77.16, 76.91, 51.10, 48.40, 47.19, 42.41, 39.71, 39.42, 36.72, 30.47, 29.58, 26.57, 25.79, 17.77, 16.41; HRMS (TOF-ESI) m/z [M + H]^+^ calculated for [C_23_H_41_N_2_]^+^ 345.3270; found 345.3268.

#### N-((1r,3r,5r,7r)-adamantan-2-yl)-3-bromopropanamide **(IM-26)**

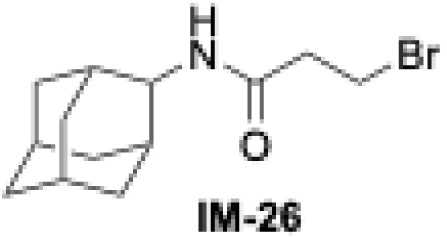

To a stirred solution of 2-adamantylamine (800 mg, 5.29 mmol) in DCM (10 mL), an aqueous solution (5 mL) of K_2_CO_3_ (1.10 g, 7.94 mmol) was added at 0 °C. To this, 3-bromopropionyl chloride (1.09 g, 6.35 mmol) dissolved in DCM (10 mL) was added dropwise over 30 min and the RM was stirred for 24 h at 30 °C.^19^ Then, the RM was extracted with DCM (2 × 20 mL), and the combined organic layer was washed with NaHCO_3_ followed by 3% v/v HCl and brine, dried over Na_2_SO_4_, filtered, and concentrated *in vacuo.* The crude product was purified by silica gel flash chromatography (0−2% MeOH/DCM) to afford **IM-26** (1.12 g, 73%) as an off-white powder. ^1^H NMR (400 MHz, CDCl_3_) *δ* 5.90 (s, 1H), 4.09 (s, 1H), 3.68 – 3.64 (m, 2H), 2.80 – 2.76 (m, 2H), 1.94 – 1.63 (m, 14H).

#### N-((1r,3r,5r,7r)-adamantan-2-yl)-3-(phenethylamino)propanamide **(IM-27)**

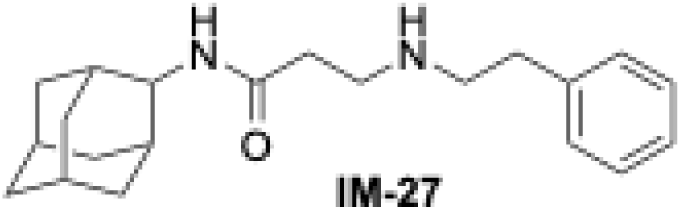

To a solution of 3-bromopropanamide **IM-26** (300 mg, 1.04 mmol) and phenethylamine (188 mg, 1.56 mmol) in DMF (5 mL), was added K_2_CO_3_ (361 mg, 2.62 mmol), and stirred at 70 °C for 20 h.^19^ The RM was cooled to RT, diluted with water (30 mL) and extracted with EtOAc (2 × 25 mL).

The combined organic layers were washed with brine, dried over Na_2_SO_4_, filtered, and concentrated *in vacuo.* The crude extract was purified by silica gel flash chromatography (0−3% MeOH/DCM) to afford **IM-27** (175 mg, 51%) as a yellow oil. ^1^H NMR (400 MHz, CDCl_3_) *δ* 8.48 (s, 1H), 7.33 – 7.30 (m, 2H), 7.26 – 7.18 (m, 3H), 4.05 (d, *J* = 8.4 Hz, 1H), 2.96 – 2.91 (m, 4H), 2.84-2.82 (m, 2H), 2.38 (t, *J* = 6.0 Hz, 2H), 1.93 – 1.81 (m, 11H), 1.75 (s, 2H), 1.65 (d, *J* = 12.8 Hz, 2H).

#### N^1^-((1r,3r,5r,7r)-adamantan-2-yl)-N^3^-phenethylpropane-1, 3-diamine **(12)**

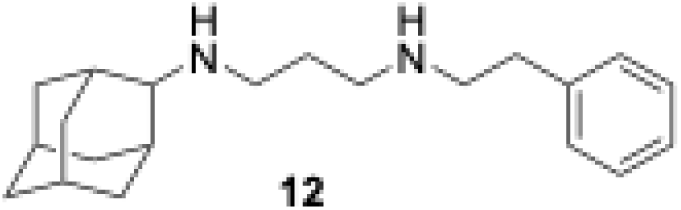

To a solution of propanamide, **IM-27** (120 mg, 0.370 mmol) in dry DCM (5 mL) at 0 °C, was added TMSCl (80.0 mg, 0.741 mmol) and stirred for 15 min under N_2_ atmosphere. A suspension of LiAlH_4_ (35 mg, 0.92 mmol) in THF (1 mL) was added to RM at −10 °C, and stirring continued for another 3 h at −10 to 0 °C.^31^ The reaction was quenched with 10% NaOH solution, the inorganic precipitate was filtered off, and the organic layer was separated. The aqueous phase was extracted with DCM (2 × 20 mL), and the combined organic layer was dried over Na_2_SO_4_, filtered, and concentrated *in vacuo.* The crude product was purified by silica gel flash chromatography (DCM:MeOH:NH_4_OH; 90:8:2) to afford **12** (90 mg, 78%) as a colorless oil and converted into the HCl salt (an off-white solid) using a 2 M HCl solution in diethyl ether. Free base; ^1^H NMR (500 MHz, CDCl_3_) *δ* 7.30 – 7.27 (m, 2H), 7.21 – 7.18 (m, 3H), 2.88 (t, *J* = 7.0 Hz, 2H), 2.82 – 2.79 (m, 2H), 2.71 (t, *J* = 7.0 Hz, 2H), 2.68 (s, 1H), 2.64 (t, *J* = 7.0 Hz, 2H), 1.91 (d, *J* = 12.5 Hz, 2H), 1.84 – 1.83 (m, 6H), 1.77 (s, 1H), 1.70 – 1.66 (m, 7H), 1.48 (d, *J* = 12.5 Hz, 2H); ^13^C NMR (125 MHz, CDCl_3_) δ 139.97, 128.53, 128.28, 125.94, 77.16, 76.91, 76.65, 61.66, 51.21, 48.62, 45.50, 37.80, 37.41, 36.30, 31.78, 31.20, 30.33, 27.67, 27.45; HRMS (TOF-ESI) m/z [M + H]^+^ calculated for [C_21_H_33_N_2_]^+^ 313.2644; found 313.2641.

#### N-((1r,3r,5r,7r)-adamantan-2-yl)-2-chloroacetamide **(IM-28)**

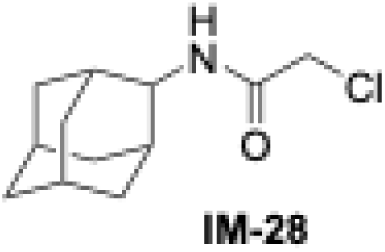

To a stirred solution of 2-adamantylamine (800 mg, 5.30 mmol) in DCM (10 mL), was added an aqueous solution (10 mL) of K_2_CO_3_ (1.09 g, 7.95 mmol) at 0 °C. Chloroacetyl chloride (712 mg, 6.36 mmol) dissolved in DCM (10 mL) was added dropwise over 30 min to the reaction mixture and stirred for 24 h.^12^ The RM was extracted with DCM (2 × 20 mL), and the combined organic layer was washed with Na_2_CO_3_ followed by 3% v/v HCl and brine, dried over Na_2_SO_4_, filtered, and concentrated *in vacuo.* The crude product was purified by silica gel flash chromatography (0−20% EtOAc/hexane) to afford **IM-28** (850 mg, 70%) as an off-white powder. ^1^H NMR (400 MHz, CDCl_3_) *δ* 6.99 (s, 1H), 4.07 (s, 3H), 1.94 – 1.66 (m, 14H).

#### N-((1r,3r,5r,7r)-adamantan-2-yl)-2-(methyl(phenethyl)amino) acetamide **(IM-29)**

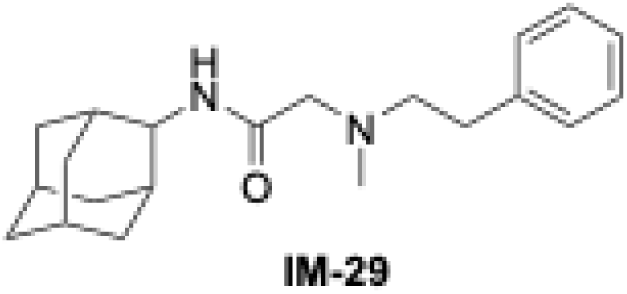

To a solution of chloroacetamide **IM-28** (300 mg, 1.32 mmol) and *N-*Methylphenethylamine (270 mg, 1.98 mmol) in DMF (5 mL), was added K_2_CO_3_ (455 mg, 3.30 mmol), and stirred at 70 °C for 20 h.^19^ The RM was cooled to RT, diluted with water (30 mL), and extracted with EtOAc (2 × 25 mL). The combined organic layers were washed with brine, dried over Na_2_SO_4_, filtered, and concentrated *in vacuo.* The crude extract was purified by silica gel flash chromatography (0−4% MeOH/DCM) to afford **IM-29** (410 mg, 95%) as a colorless oil. ^1^H NMR (400 MHz, CDCl_3_) *δ* 7.54 (s, 1H), 7.30 – 7.26 (m, 2H), 7.21 – 7.17 (m, 3H), 4.01 (d, *J* = 8.4 Hz, 1H), 3.05 (s, 2H), 2.79 – 2.75 (m, 4H), 2.36 (s, 3H), 1.85 – 1.71 (m, 10H), 1.57 (s, 4H).

#### N^1^-((1r,3r,5r,7r)-adamantan-2-yl)-N^2^-methyl-N^2^-phenethylethane-1,2-diamine **(13)**

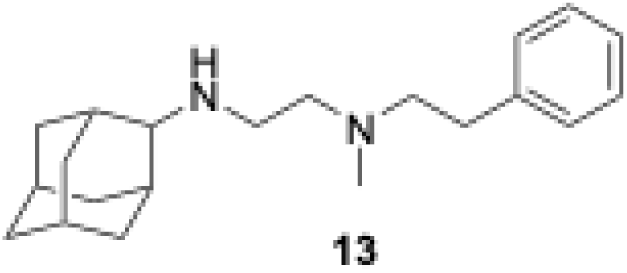

To a solution of acetamide **IM-29** (150 mg, 0.46 mmol) in dry DCM (5 mL) at 0 °C, was added TMSCl (100 mg, 0.922 mmol) and stirred for 15 min under N_2_ atmosphere. A suspension of LiAlH_4_ (44.0 mg, 1.15 mmol) in THF (1 mL) was added to the RM at −10 °C and stirred continuously overnight.^31^ The reaction was quenched with 10% NaOH solution, the inorganic precipitate was filtered off, and the organic layer was separated. The aqueous phase was extracted with DCM (2 × 20 mL), and the combined organic layer was dried over Na_2_SO_4_, filtered, and concentrated *in vacuo.* The crude product was purified by silica gel flash chromatography (DCM:MeOH:NH_4_OH; 90:9:1) to afford **13** (95.0 mg, 66%) as a colorless oil and converted into the HCl salt (white crystalline powder) using a 2 M HCl solution in diethyl ether. Free base; ^1^H NMR (500 MHz, CDCl_3_) *δ* 7.29 – 7.26 (m, 2H), 7.21 – 7.18 (m, 3H), 2.78 (t, *J* = 7.5 Hz, 2H), 2.72 – 2.70 (m, 3H), 2.67 – 2.60 (m, 4H), 2.30 (s, 3H), 1.88 – 1.83 (m, 8H), 1.76 (s, 1H), 1.71 – 1.68 (m, 4H), 1.48 (d, *J* = 13.0 Hz, 2H); ^13^C NMR (125 MHz, CDCl_3_) δ 140.60, 128.83, 128.46, 126.05, 77.41, 77.16, 76.90, 62.08, 59.58, 56.98, 44.23, 42.06, 37.95, 37.65, 33.85, 31.78, 31.31, 27.82, 27.55; HRMS (TOF-ESI) m/z [M + H]^+^ calculated for [C_21_H_33_N_2_]^+^ 313.2644; found 313.2646.

#### N-((1r,3r,5r,7r)-adamantan-2-yl)-3-(((E)-3,7-dimethylocta-2,6-dien-1-yl)amino)propanamide **(IM-30)**

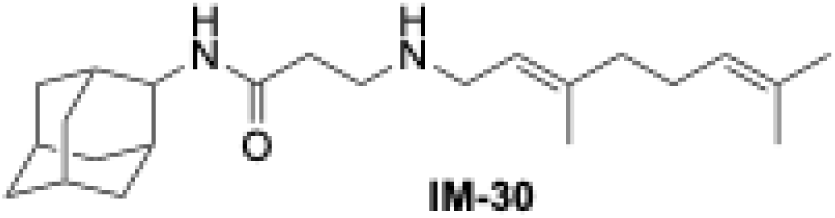

To a solution of 3-bromopropanamide **IM-26** (500 mg, 1.74 mmol) and (*E*)-3,7-dimethylocta-2,6- dien-1-amine (401 mg, 2.63 mmol) in DMF (5 mL), was added K_2_CO_3_ (603 mg, 4.37 mmol), and the RM was stirred at 70 °C for 20 h.^19^ The RM was cooled to RT, diluted with water (30 mL) and extracted with EtOAc (2 × 25 mL). The combined organic layer was washed with brine, dried over Na_2_SO_4_, filtered, and concentrated *in vacuo.* The crude extract was purified by silica gel flash chromatography (0−5% MeOH/DCM) to afford **IM-30** (310 mg, 49%) as a yellow oil. ^1^H NMR (500 MHz, CDCl_3_) *δ* 8.64 (s, 1H), 5.23 (t, *J* = 7.0 Hz, 1H), 5.07 (d, *J* = 7.0 Hz, 1H), 4.12 – 4.06 (m, 1H), 3.27 – 3.23 (m, 2H), 2.90 (t, *J* = 6.0 Hz, 2H), 2.38 (t, *J* = 6.0 Hz, 2H), 2.08 – 1.99 (m, 4H), 1.88 – 1.81 (m, 10H), 1.76 – 1.60 (m, 14H).

#### N-((1r,3r,5r,7r)-adamantan-2-yl)-3-(((E)-3,7-dimethylocta-2,6-dien-1-yl)(methyl)amino) propanamide **(IM-31)**

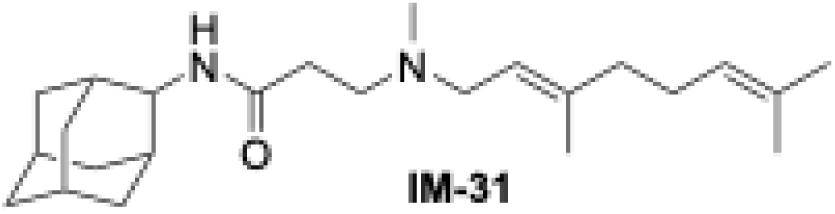

To a stirred solution of propanamide **IM-30** (150 mg, 0.420 mmol) in MeOH (5 mL) was added 37% formaldehyde in H_2_O (0.200 mL, 2.50 mmol) and stirred at RT for 1 h followed by heating at 50 °C for an additional 1 h under N_2_. The RM was then cooled to RT, and NaBH_4_ (110 mg, 2.93 mmol) was added slowly and stirred overnight.^30^ The RM was concentrated *in vacuo*, and the crude extract was dissolved in EtOAc (20 mL) and washed successively with water (3 × 20 mL) and brine (20 mL). The organic layer was dried over Na_2_SO_4_ and concentrated *in vacuo,* and the crude extract was purified via silica gel flash chromatography (0−5% MeOH/DCM) to afford **IM-31** (100 mg, 64%) as a colorless oil. ^1^H NMR (400 MHz, CDCl_3_) *δ* 9.18 (s, 1H), 5.21 (t, *J* = 6.8 Hz, 1H), 5.06 (t, *J* = 6.8 Hz, 1H), 4.04 (d, *J* = 8.4 Hz, 1H), 3.05 (d, *J* = 6.8 Hz, 2H), 2.64 (t, *J* = 6.0 Hz, 2H), 2.41 (t, *J* = 6.0 Hz, 2H), 2.25 (s, 3H), 2.09 – 2.00 (m, 4H), 1.87 – 1.79 (m, 10H), 1.73 (s, 2H), 1.67 – 1.59 (m, 11H).

#### N^1^-((1r,3r,5r,7r)-adamantan-2-yl)-N^3^-((E)-3,7-dimethylocta-2,6-dien-1-yl)-N^3^-methylpropane- 1,3-diamine **(14)**

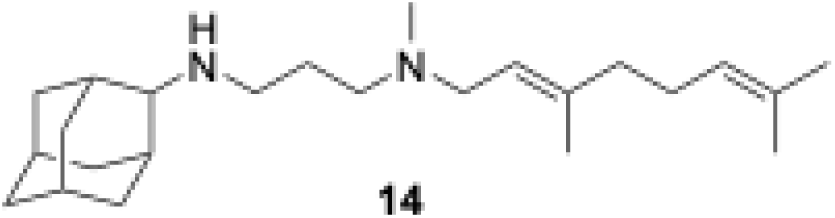

To a solution of propanamide **IM-31** (100 mg, 0.27 mmol) in dry DCM (5 mL) at 0 °C, was added TMSCl (58.0 mg, 0.54 mmol) and stirred for 15 min under N_2_ atmosphere. A suspension of LiAlH_4_ (26.0 mg, 0.670 mmol) in THF (0.5 mL) was added to the RM at −10 °C and stirred overnight.^31^ The reaction was quenched with 10% NaOH solution, the inorganic precipitate was filtered off, and the organic layer was separated. The aqueous phase was extracted with DCM (2 × 15 mL), and the combined organic layer was dried over Na_2_SO_4_, filtered, and concentrated *in vacuo.* The crude product was purified by silica gel flash chromatography (DCM:MeOH:NH_4_OH; 90:9:1) to afford **14** (75.0 mg, 77%) as a colorless oil and converted into the HCl salt (an off-white solid) using 2 M HCl solution in diethyl ether. Free base; ^1^H NMR (500 MHz, CDCl_3_) ^1^H NMR (500 MHz, CDCl_3_) *δ* 5.21 (t, *J* = 7.0 Hz, 1H), 5.07 (t, *J* = 7.0 Hz, 1H), 2.95 (d, *J* = 7.0 Hz, 2H), 2.73 (s, 1H), 2.66 (d, *J* = 7.0 Hz, 2H), 2.40 (d, *J* = 7.0 Hz, 2H), 2.20 (s, 3H), 2.10 – 2.05 (m, 2H), 2.02 – 1.99 (m, 2H), 1.93 – 1.90 (m, 4H), 1.84 – 1.82 (m, 3H), 1.77 – 1.66 (m, 11H), 1.61 (s, 3H), 1.58 (s, 3H), 1.50 (d, *J* = 12.5 Hz, 2H); ^13^C NMR (125 MHz, CDCl_3_) δ 138.53, 131.62, 124.25, 121.50, 121.47, 121.44, 77.42, 77.16, 76.91, 61.93, 56.33, 55.52, 45.95, 42.12, 39.92, 37.99, 37.63, 31.77, 31.40, 27.86, 27.64, 26.55, 25.83, 17.78, 16.49; HRMS (TOF-ESI) m/z [M + H]^+^ calculated for [C_24_H_43_N_2_]^+^ 359.3426; found 359.3425.

#### N-((3s,5s,7s)-adamantan-1-yl)-2-(((E)-3,7-dimethylocta-2,6-dien-1-yl)(methyl)amino)acetamide **(IM-32)**

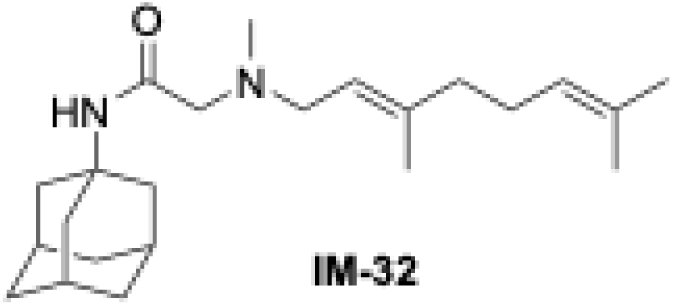

To a stirred solution of acetamide **IM-23** (200 mg, 0.58 mmol) in MeOH (5 mL) was added 37% formaldehyde in H_2_O (0.330 mL, 3.49 mmol) and stirred at RT for 1 h followed by heating at 50 °C for an additional 1 h under N_2_. The RM was then cooled to RT, and NaBH_4_ (172 mg, 4.65 mmol) was added slowly and stirred overnight.^30^ The RM was concentrated *in vacuo*, and the crude extract was dissolved in EtOAc (20 mL) and washed successively with water (3 × 20 mL) and brine (20 mL). The organic layer was dried over Na_2_SO_4_ and concentrated *in vacuo,* and the crude extract was purified via silica gel flash chromatography (0−5% MeOH/DCM) to afford **IM-32** (135 mg, 65%) as a colorless oil. ^1^H NMR (400 MHz, CDCl_3_) δ 6.96 (s, 1H), 5.17 (d, *J* = 6.8 Hz, 1H), 5.07 (t, *J* = 7.2 Hz, 1H), 2.98 (d, *J* = 6.8 Hz, 2H), 2.84 (s, 2H), 2.22 (s, 3H), 2.10 – 2.00 (m, 12H), 1.71 – 1.65 (m, 10H), 1.62 (s, 3H), 1.60 (s, 3H).

#### N^1^-((3s,5s,7s)-adamantan-1-yl)-N^2^-((E)-3,7-dimethylocta-2,6-dien-1-yl)-N^2^-methylethane-1,2- diamine **(15)**

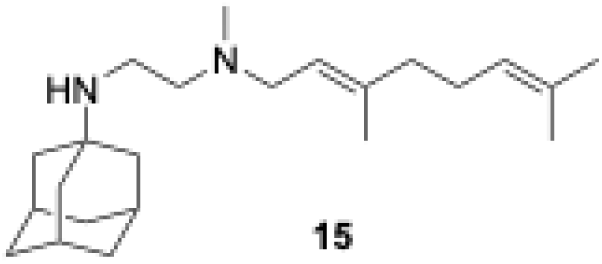

To a solution of acetamide **IM-32** (110 mg, 0.291 mmol) in dry DCM (5 mL) at 0 °C, was added TMSCl (63.0 mg, 0.591 mmol) and stirred for 15 min under N_2_ atmosphere. A suspension of LiAlH_4_ (28.0 mg, 0.74 mmol) in THF (0.5 mL) was added to the RM at −10 °C and stirred overnight.^19^ The reaction was quenched with 10% NaOH solution, the inorganic precipitate was filtered off, and the organic layer was separated. The aqueous phase was extracted with DCM (2 × 15 mL), and the combined organic layer was dried over Na_2_SO_4_, filtered, and concentrated *in vacuo.* The crude product was purified by silica gel flash chromatography (DCM:MeOH:NH_4_OH, 90:9:1) to afford **15** (80.0 mg, 80%) as an off-yellow oil and converted into the HCl salt (an off-white solid) using a 2 M HCl solution in diethyl ether. Free base; ^1^H NMR (500 MHz, CDCl_3_) *δ* 5.23 (t, *J* = 7.0 Hz, 1H), 5.08 (t, *J* = 7.0 Hz, 1H), 2.95 (d, *J* = 7.0 Hz, 2H), 2.69 (d, *J* = 6.0 Hz, 2H), 2.47 (d, *J* = 6.0 Hz, 2H), 2.16 (s, 3H), 2.13 – 1.96 (m, 8H), 1.69 – 1.57 (m, 21H); ^13^C NMR (125 MHz, CDCl_3_) δ 138.43, 131.59, 124.30, 121.51, 77.41, 77.16, 76.90, 57.52, 55.32, 50.44, 42.70, 41.93, 39.94, 37.83, 36.87, 29.70, 26.62, 25.85, 17.82, 16.49; HRMS (TOF-ESI) m/z [M + H]^+^ calculated for [C_23_H_41_N_2_]^+^ 345.3270; found 345.3269.

### Cell Growth Inhibition Assays

#### IMV assay

Proton translocation into IMVs was measured by the decrease of ACMA (9-amino-6- chloro-2-methoxyacridine) fluorescence, as previously reported. The excitation and emission wavelengths were 410 and 480 nm, respectively. IMVs (2 mg/mL membrane protein) were preincubated at 37 °C in 10 mM MOPS-NaOH (pH 7.0), 100 mM KCl, 5 mM MgCl_2_ containing 2 μM ACMA, and the baseline was monitored for 2 min. The reaction was then initiated by adding 1 mM ATP. When the signal had stabilized (∼6 min), compounds were added in a suitable concentration range, and proton translocation was measured fluorometrically.

#### *T. gondii* cell growth inhibition assay

*T. gondii* tachyzoites expressing a td-Tomato red fluorescent protein (RFP) were purified by passing them through a 25-gauge needle, followed by filtration through a 5 μm filter^16^. Human fibroblasts were cultured in 96 well black plates for 48 hours prior to the addition of 4000 tachyzoites/well plus the compounds. Fluorescence values were measured for up to 7 days and both excitation (544 nm) and emission (590 nm) were read from the bottom of the plates in a BioTek Synergy H1 plate reader. The EC_50_s were calculated using Prism software.

#### *S. cerevisiae* cell growth inhibition assay

*S. cerevisiae* (ATCC208352, grown for 48 h at 30 °C) was diluted 50-fold in YPD media and grown to an OD_600_ of ∼0.4, which occurred after about 6.5 hrs. The culture was then diluted 500-fold into a fresh YPD medium to give a working solution. This working solution (180 μL) was transferred into every well in a flat-bottomed 96-well plate except for the first well of columns of interest. Inhibitors were added at specific starting concentrations with a total volume of 360 μL (diluted with working solution) to the first well of the columns. The inhibitors were then sequentially diluted 2-fold across 12 wells; 180 μL of autoclaved Milli-Q water was added to empty peripheral wells to prevent water evaporation from the plate. Plates were incubated at 30 °C, with shaking at 200 rpm for 48 h. The OD_600_ values were then measured and used to determine cell growth inhibition, by using GraphPad Prism software (version 7.04). Experiments were carried out in duplicate or triplicate. IC_50_ values were determined using a four-parameter variable-slope function in the Prism program.

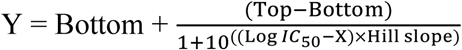

#### *S. cerevisiae* (RH6829) cell growth inhibition assay

These assays were conducted using the same procedure as described for S. cerevisiae cell growth inhibition, with the exception that cholesterol mutant S. cerevisiae (RH6829) cells were used.

#### *M. smegmatis* Growth Inhibition Assay

*M. smegmatis* (grown for 36–48 h) was diluted 1000- fold in fresh Middlebrook 7H9 (plus 10% ADC supplement, Sigma: M0553-1VL; 0.5% glycerol; 0.05% Tween 80) media to generate a working solution. This working solution (180 μL) was then transferred into every well in a flat-bottomed 96-well plate except for the second column and peripheral wells. Inhibitors were added at specific starting concentrations (100 μM – 1 mM) with a total volume of 360 μL (diluted with working solution) to the second column. The inhibitors were then sequentially diluted 2-fold across 1-12 wells; 180 μL of autoclaved Milli-Q water was added to each peripheral well to prevent water evaporation from the plate. Plates were incubated at 37 °C, with shaking at 200 rpm for 48 h. The OD_600_ values were then measured to bacterial growth inhibition using GraphPad Prism software (version 7.04). Experiments were carried out in duplicate or triplicate. IC_50_ values were determined using a four-parameter variable-slope function in the Prism program.

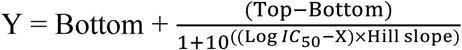

## ASSOCIATED CONTENT

### Supporting Information

The Supporting Information is available free of charge at http://pubs.acs.org.

Synthesis of carbazoles and aminoguanidine analogs **1-4** (Figure S1); Synthesis of SQ109 analogs **6-15** (Figure S2); Correlations between IMV logIC_50_ and cell growth inhibition logIC_50_ values for all 23 compounds: a) *T. gondii*. b) *S. cerevisiae*. c) *M. smegmatis* (Figure S3); Correlation matrix between R and p-values for IMV, *T. gondii*, *S. cerevisiae*, and *M. smegmatis* cell growth inhibition logIC_50_ (μM) results based on data for all compounds: a) R-values. b) p-values (Figure S4); Spectral data (^1^H NMR and ^13^CNMR); qNMR spectra; compounds smiles (Table S1).

## AUTHOR INFORMATION

### Corresponding Author

**Eric Oldfield** - *Department of Chemistry, University of Illinois at Urbana–Champaign, Urbana, IL 61801, United States.*

**Silvia N.J. Moreno -** Center for Tropical and Emerging Global Diseases and Department of Cellular Biology, University of Georgia, Athens, GA 30602, United States.

### Authors

**Davinder Singh** - Department of Chemistry, University of Illinois at Urbana-Champaign, Urbana, Illinois 61801, United States.

**Melissa A. Sleda -** Center for Tropical and Emerging Global Diseases, University of Georgia, Athens, GA 30602, United States.

**Akanksha M. Pandey-** Department of Chemistry, University of Illinois at Urbana–Champaign, Urbana, IL 61801, United States.

**Satish R. Malwal -** Department of Chemistry, University of Illinois at Urbana–Champaign, Urbana, IL 61801, United States.

**Yiyuan Chen -** Department of Chemistry, University of Illinois at Urbana-Champaign, Urbana, Illinois 61801, United States.

**Ruijie Zhou -** Department of Chemistry, University of Illinois at Urbana-Champaign, Urbana, Illinois 61801, United States.

**Feyisola Adewole** - Department of Biological, Physical and Health Sciences, College of Science, Health & Pharmacy, Roosevelt University, 425 South Wabash Avenue, Chicago, Illinois 60605, United States.

**Katie Sadowska -** Department of Biological, Physical and Health Sciences, College of Science, Health & Pharmacy, Roosevelt University, 425 South Wabash Avenue, Chicago, Illinois 60605, United States.

**Oluseye K. Onajole** - Department of Biological, Physical and Health Sciences, College of Science, Health & Pharmacy, Roosevelt University, 425 South Wabash Avenue, Chicago, Illinois 60605, United States.

## Notes

The authors declare no competing financial interests.

## ACKNOWLEDGMENTS

Some images in Figure 5 and the TOC graphic were created by the authors using BioRender.com.

## ABBREVIATIONS

MmpL3: Mycobacterial membrane protein Large 3
MenA: isoprenyl diphosphate:1,4-dihydroxy- 2-naphthoate (DHNA) isoprenyltransferase
MenG: demethylmenaquinone methyl transferase
DPPP: decaprenyl diphosphate phosphatase
ROS: reactive oxygen species
AA: ascorbic acid
RA: retinoic acid
GSH: reduced glutathione
NAC: *N*-acetyl cysteine
SNARE: SNAP Receptors
SNAP: soluble *N*-ethylmaleimide-sensitive factor attachment proteins (Sec17p in yeast
SQ109: *N*^1^-(adamantan-2-yl)-*N*^2^-[(2*E*)-3,7-dimethylocta-2,6-dien-1-yl]ethane-1,2-diamine
MIC: minimum inhibitory concentration
MFC: minimum fungicidal concentration
FICI: fractional inhibitory concentration index
FCCP: carbonyl cyanide-*p*-trifluoromethoxyphenylhydrazone
AO: acridine orange, *N,N,N′,N*′-tetramethylacridine-3,6-diamine
RM: reaction mixture
RT: room temperature
TLC: thin layer chromatography.

