## Supplementary information for "Activity of Carbazole, Aminoguanidine and Diamine Anti-infectives against *Toxoplasma gondii*"

**Table of Contents**

| **Figure S1.** Correlations between IMV logIC_50_ and cell growth inhibition logIC_50_ values for all 23 compounds. a) *T. gondii.* b) *S. cerevisiae*. c) *M. smegmatis*. | S3 |
| --- | --- |
| **Figure S2.** Correlation matrix between R and p-values for IMV, *T. gondii, S. cerevisiae*, *and M. smegmatis* cell growth inhibition logIC_50_ (μM) results based on data for all compounds. a) R-values. b) p-values. | S5 |
| **Figure S3**. Synthesis of carbazoles and aminoguanidine analogs **1−4**. | S6 |
| **Figure S4.** Synthesis of SQ109 analogs **6-15** | S7 |
| Spectral data (^1^H NMR and ^13^ CNMR) | S8 |
| qNMR | S38 |
| **Table S1.** SMILES of the compounds | S46 |

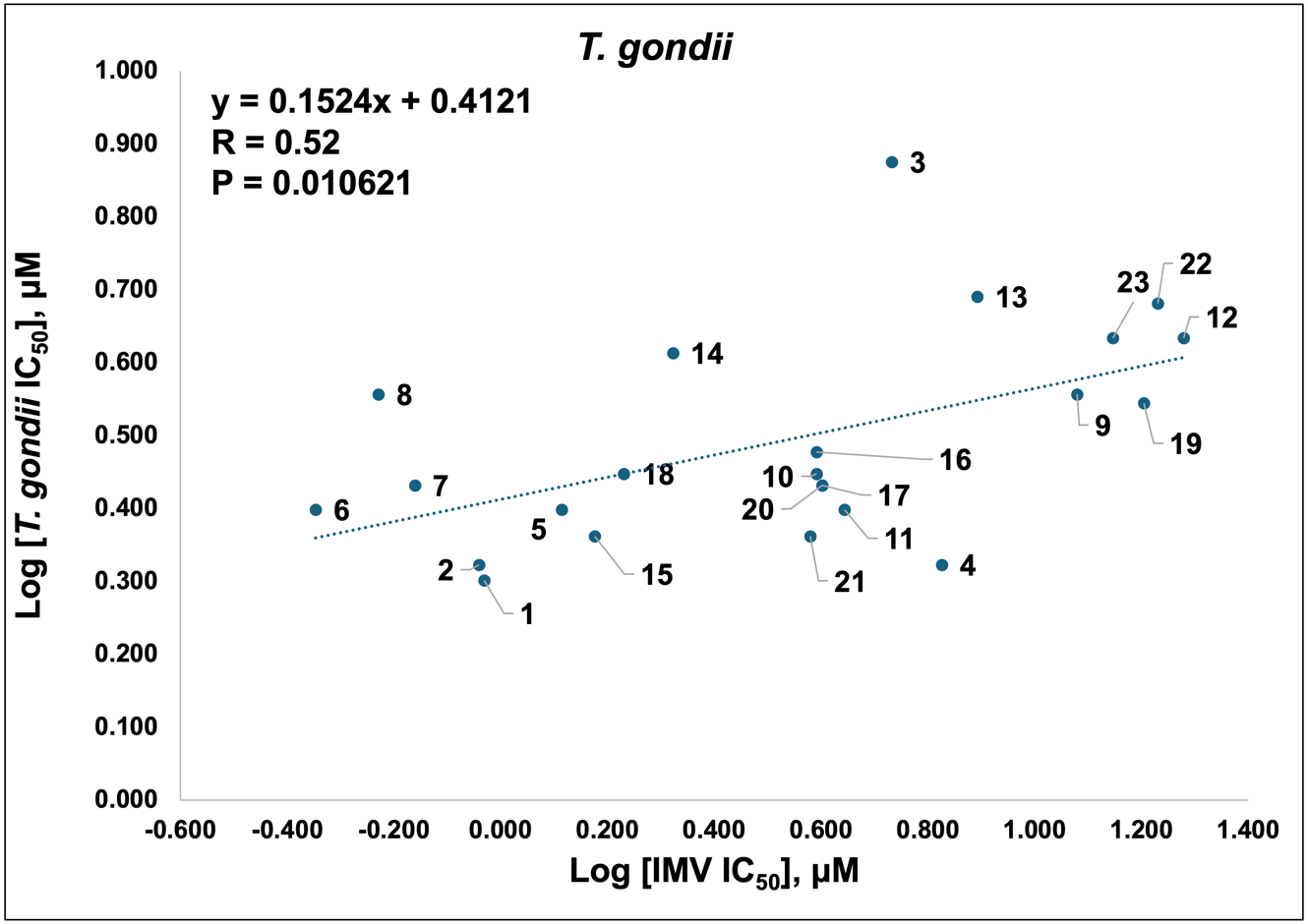

a)

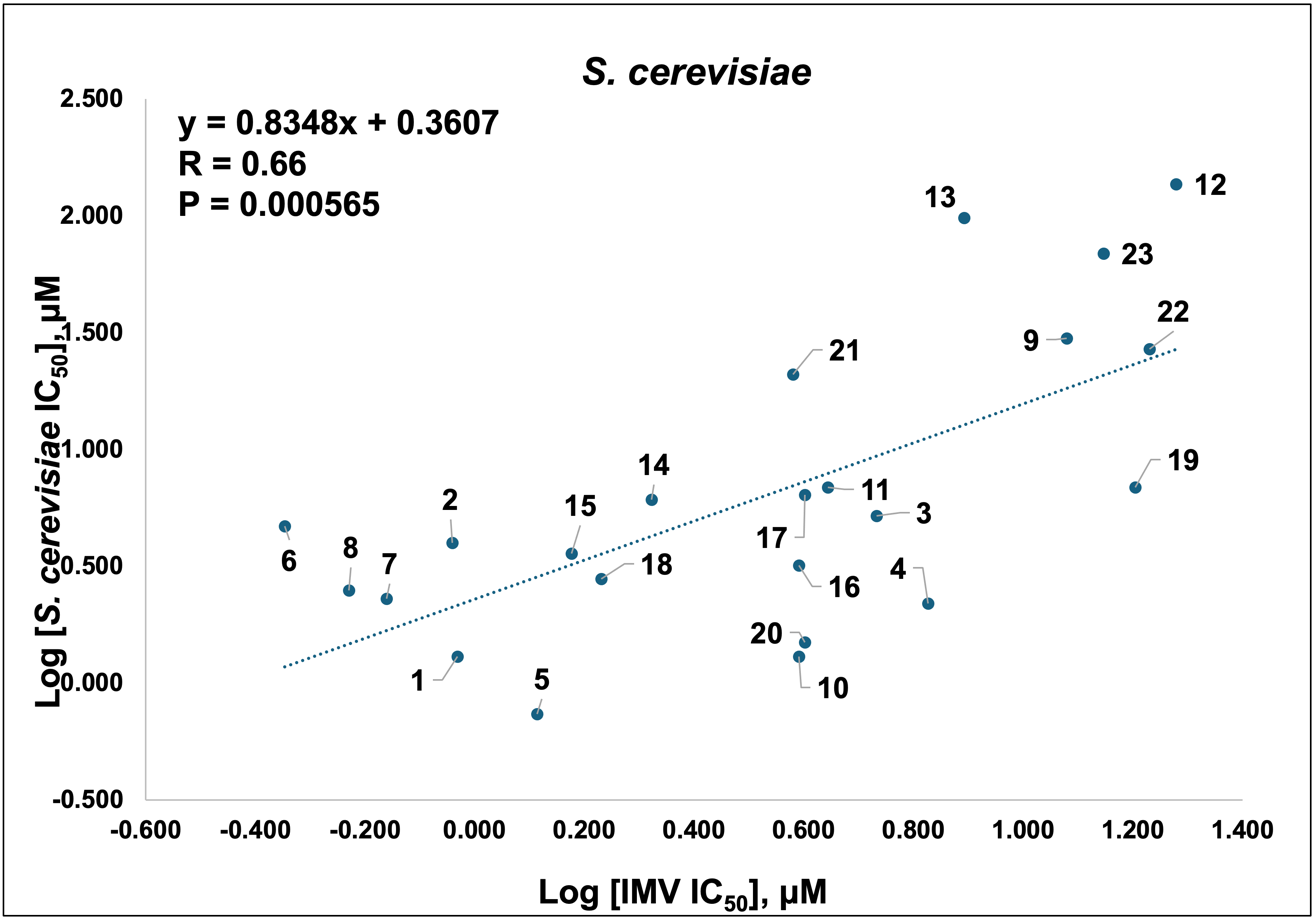

b)

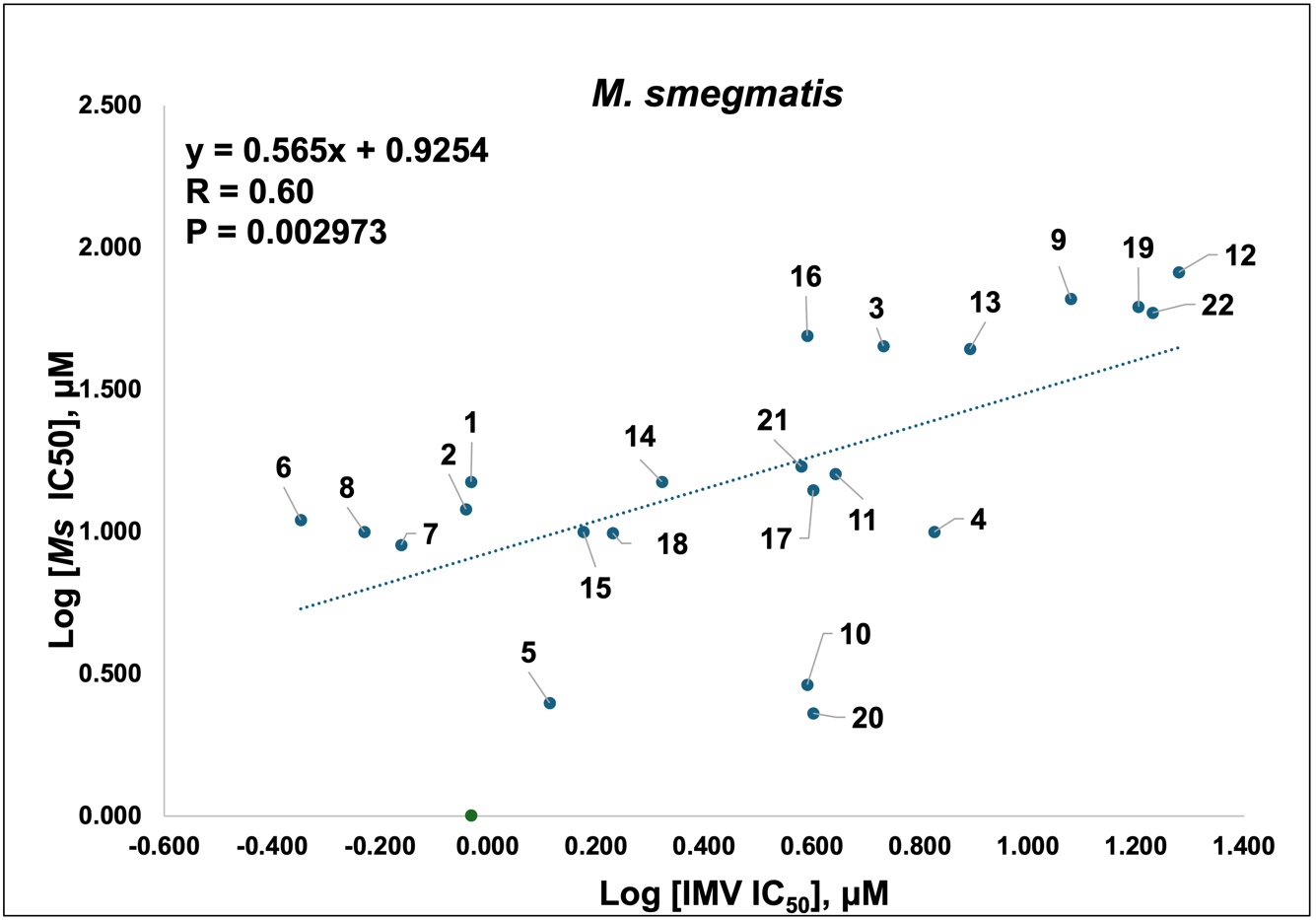

c)

**Figure S1.** Correlations between IMV logIC_50_ and cell growth inhibition logIC_50_ values for all 23 compounds. a) *T. gondii.* b) *S. cerevisiae*. c) *M. smegmatis*.

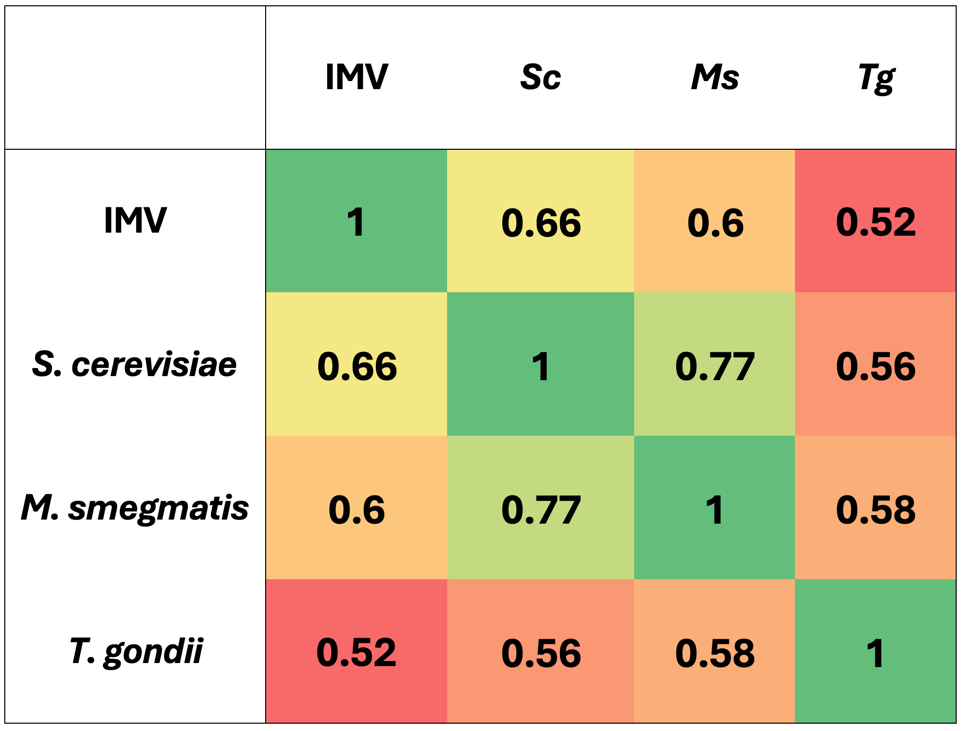

a)

b)

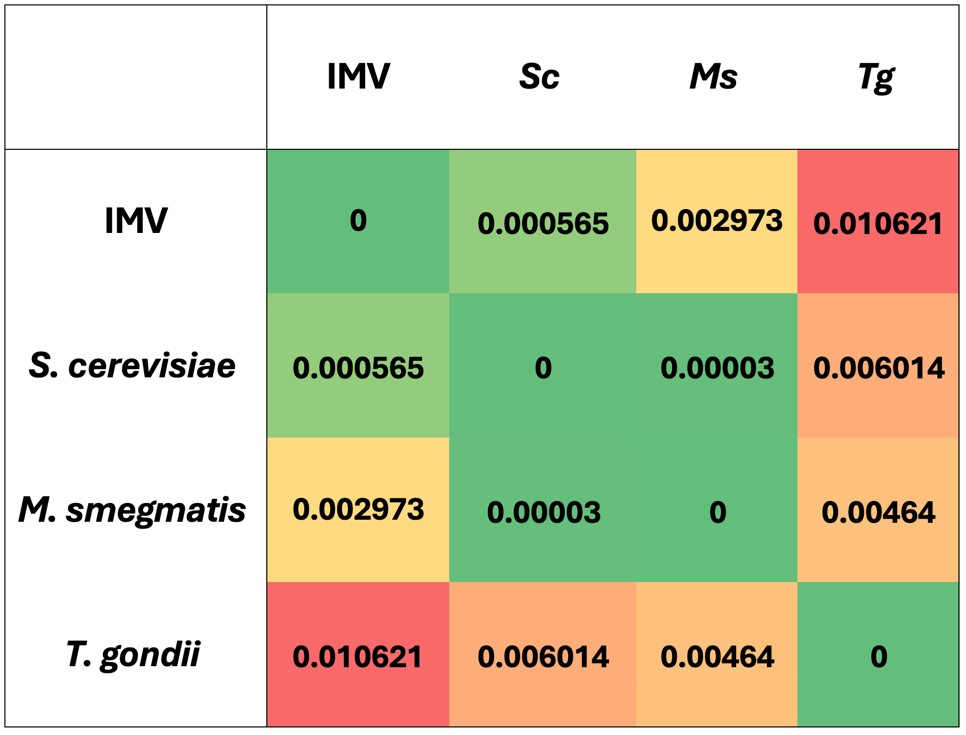

**Figure S2.** Correlation matrix between R and p-values for IMV, *T. gondii, S. cerevisiae*, *and M. smegmatis* cell growth inhibition logIC_50_ (μM) results based on data for all compounds. a) R-values. b) p-values.

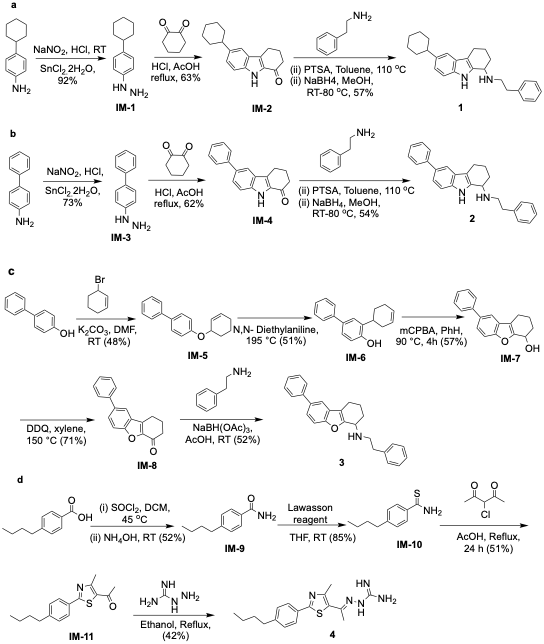

**Figure S3.** Synthesis of carbazoles and aminoguanidine analogs **1**−**4**.

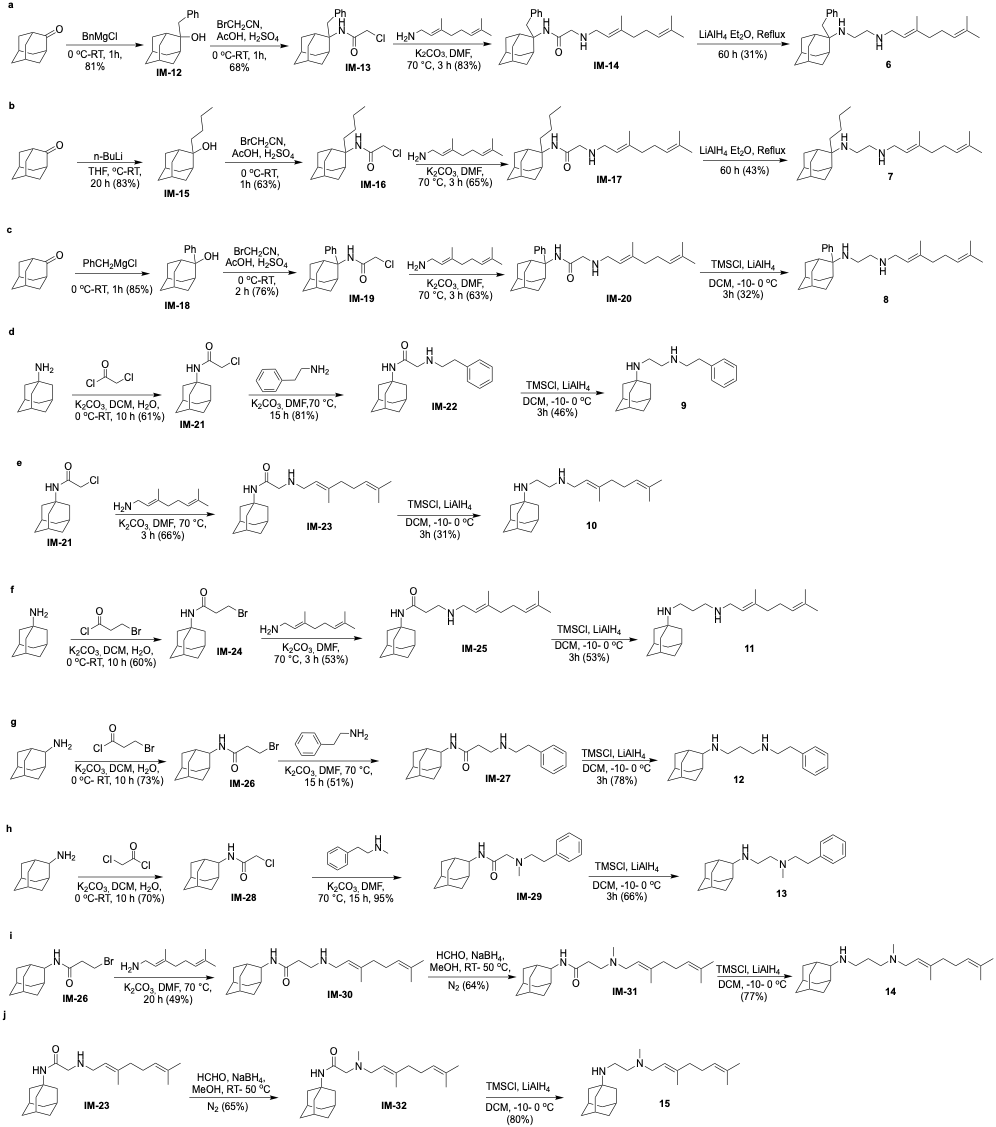

**Figure S4.** Synthesis of SQ109 analogs **6-15**

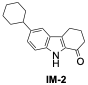

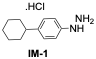

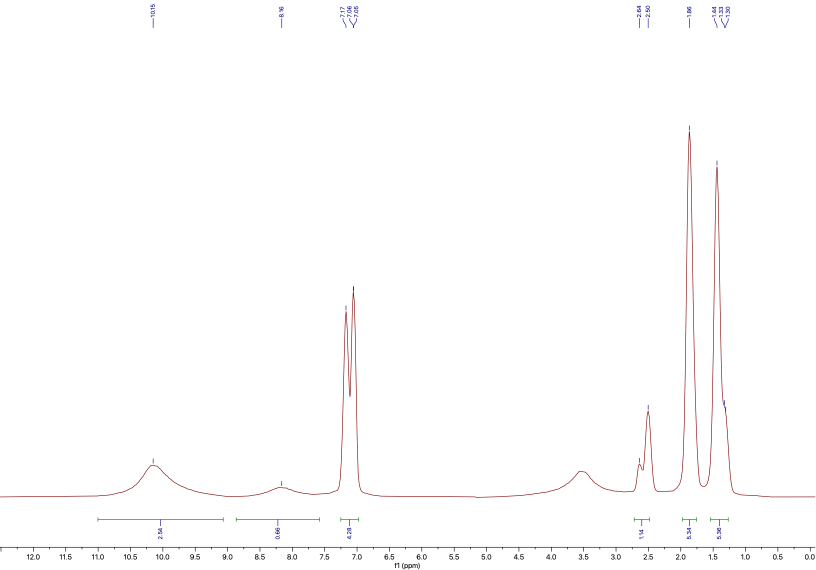

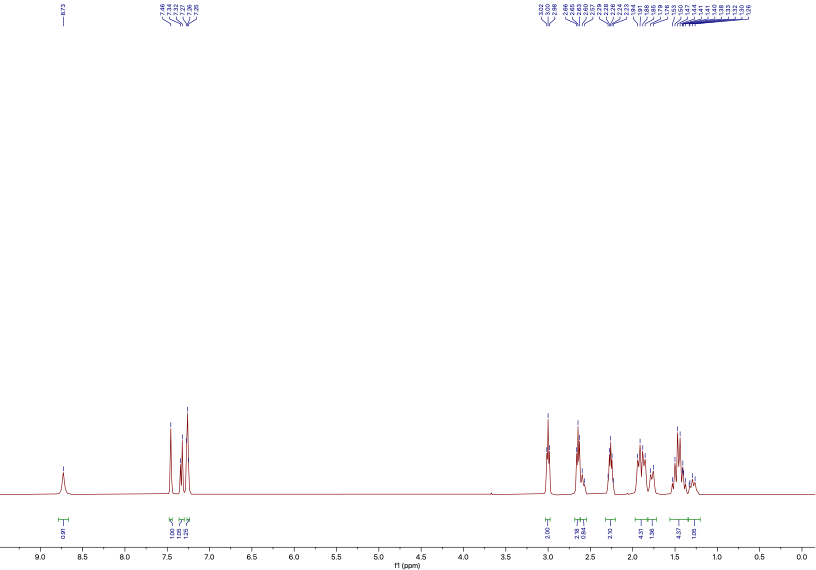

**Spectral Data: ^1^H NMR and ^13^C NMR spectra**

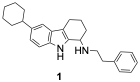

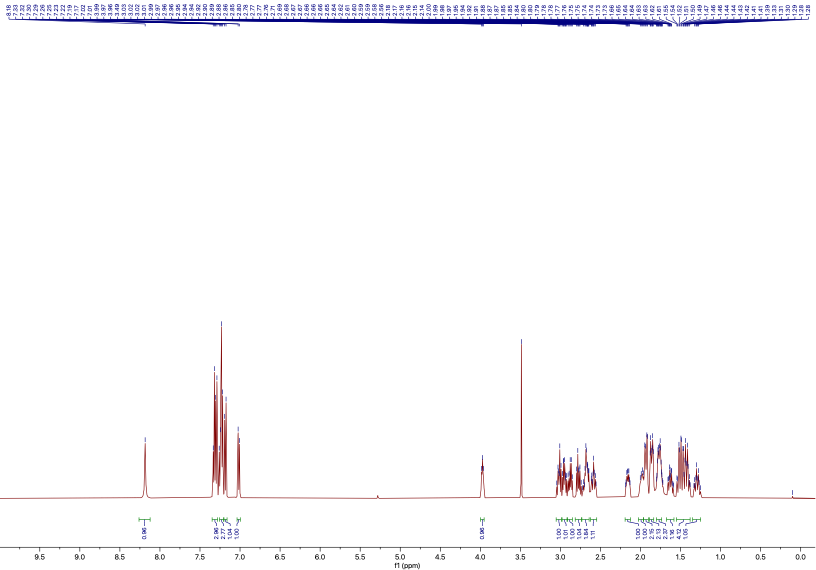

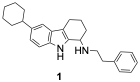

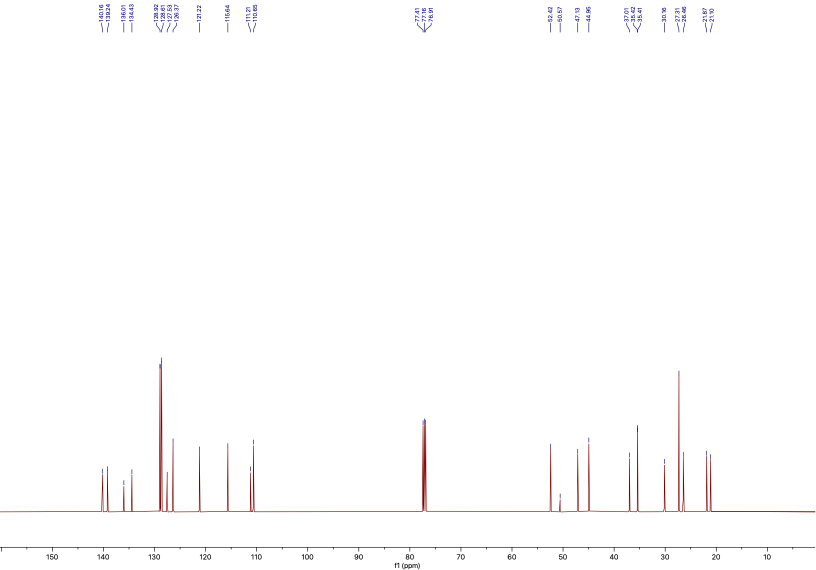

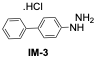

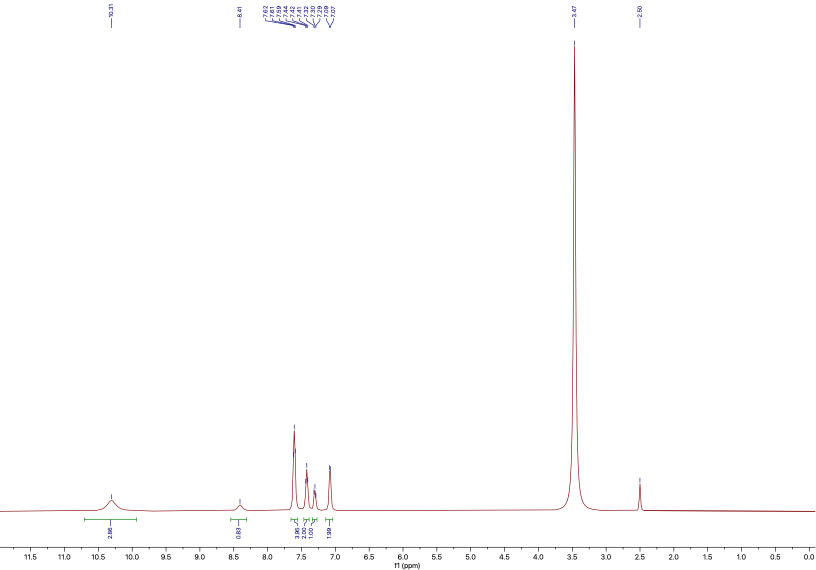

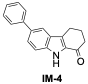

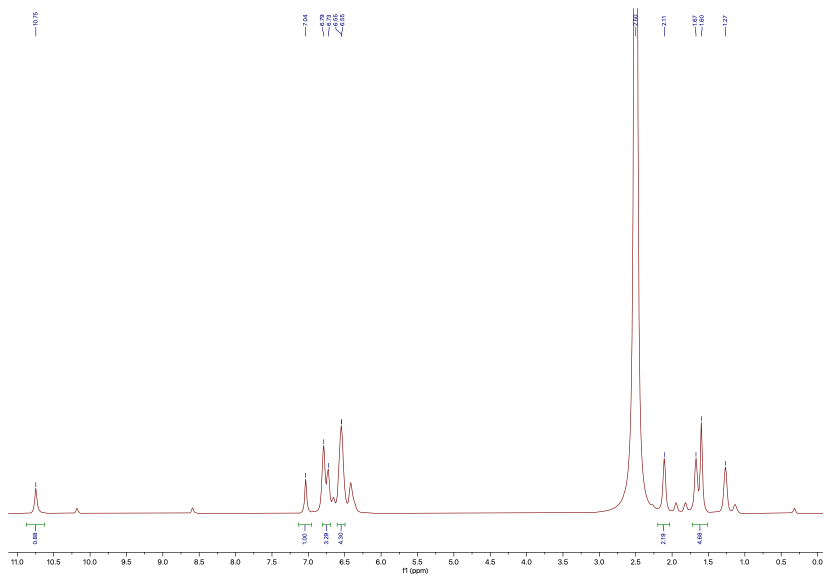

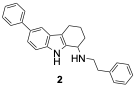

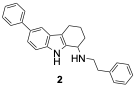

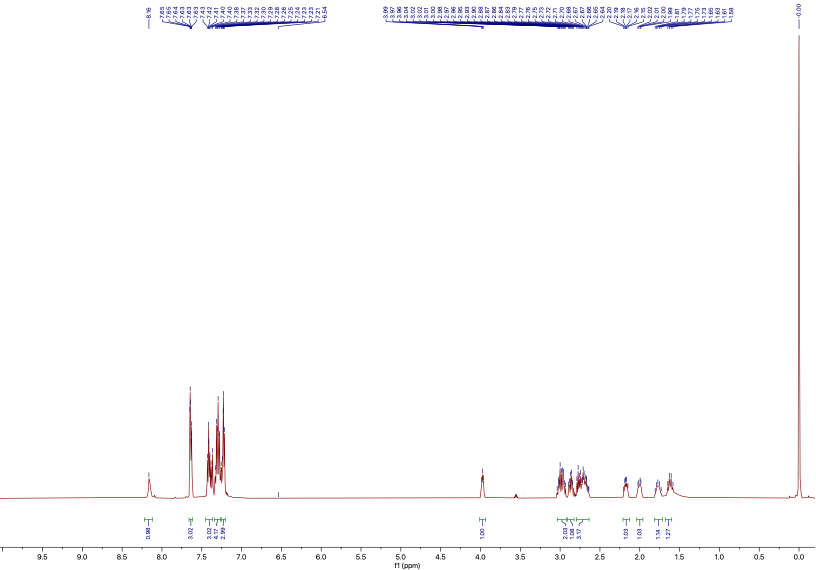

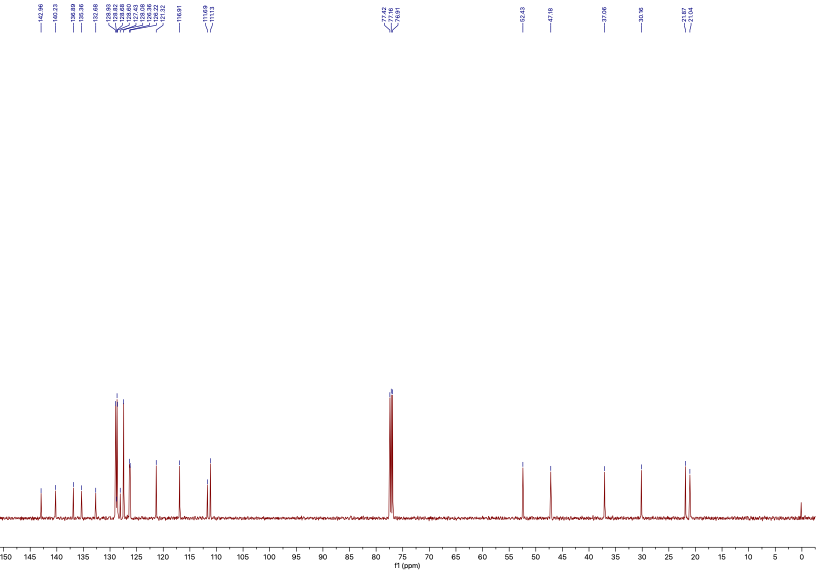

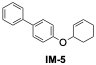

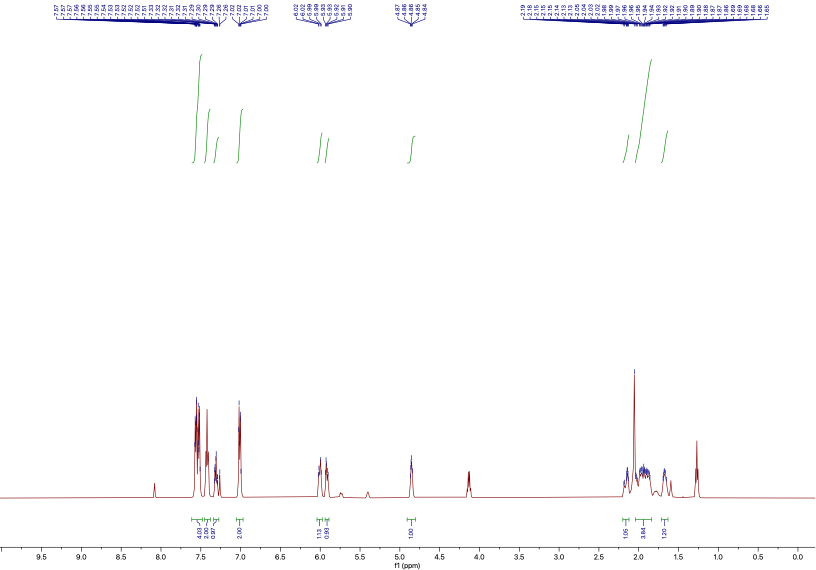

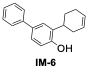

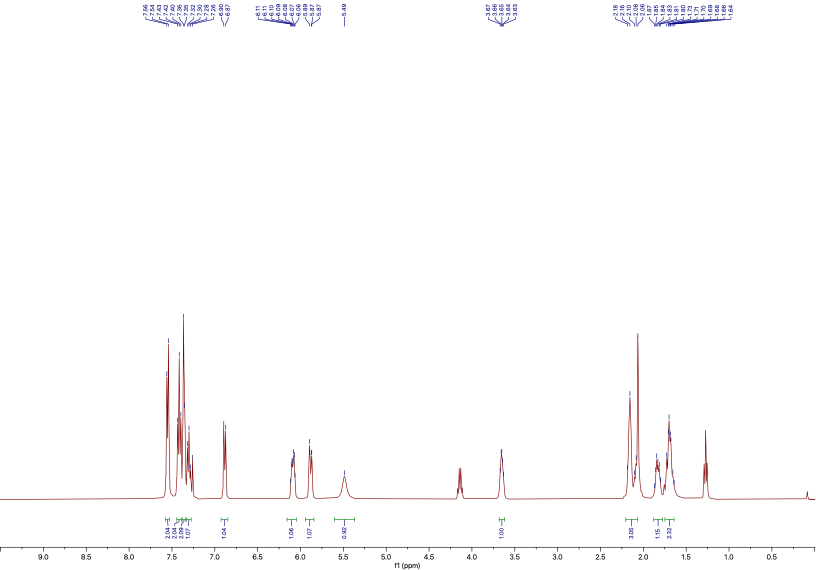

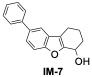

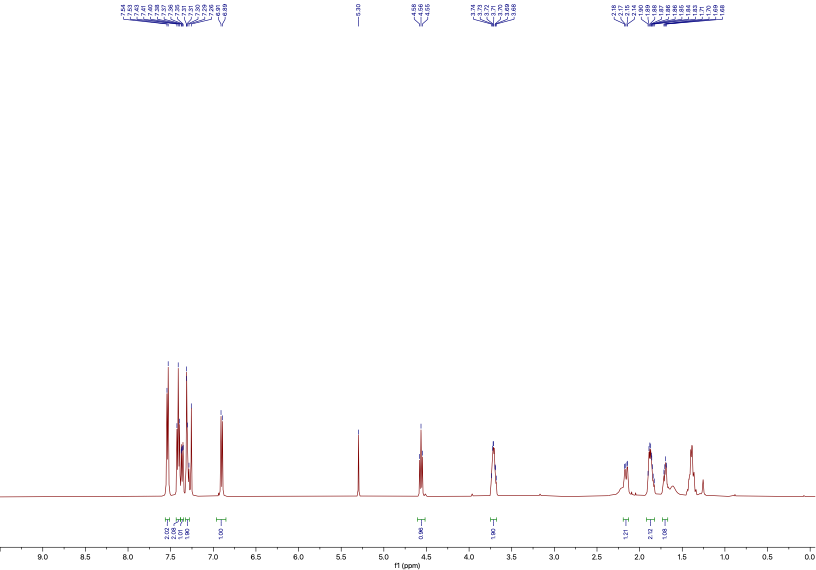

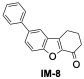

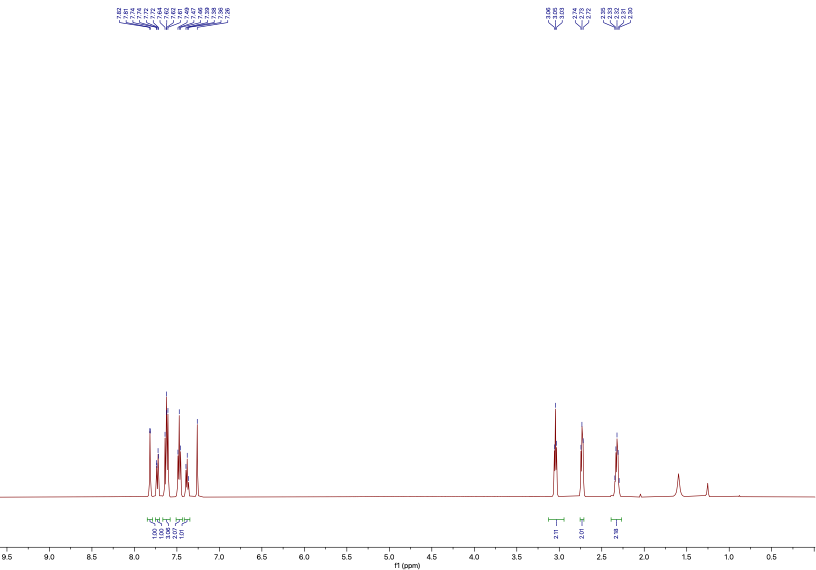

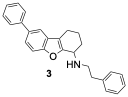

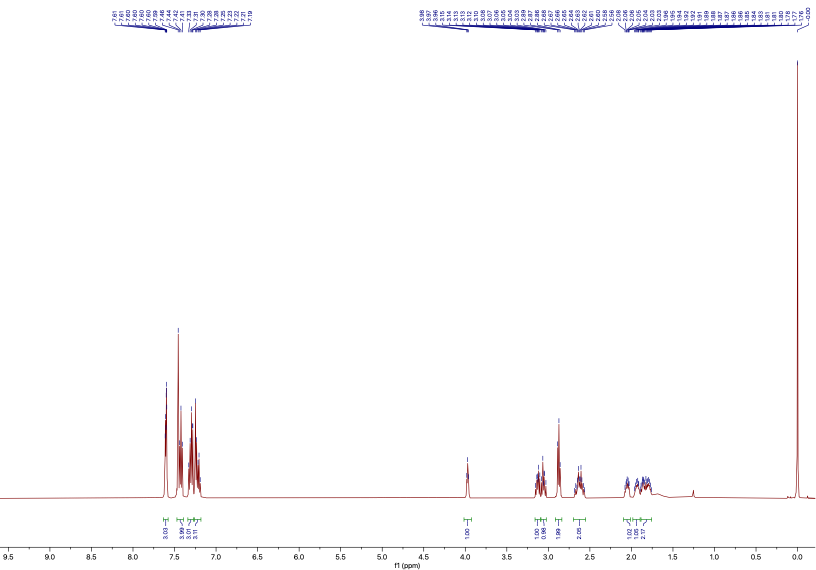

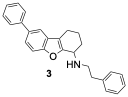

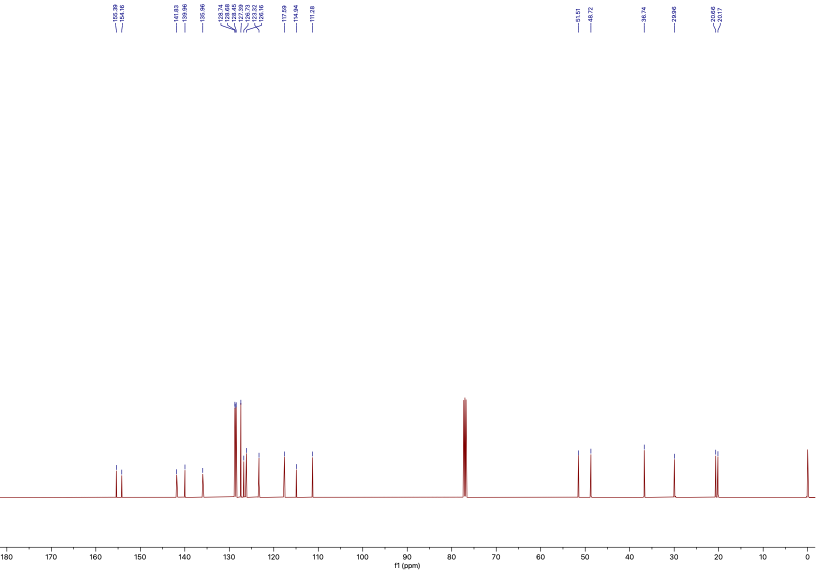

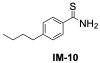

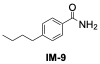

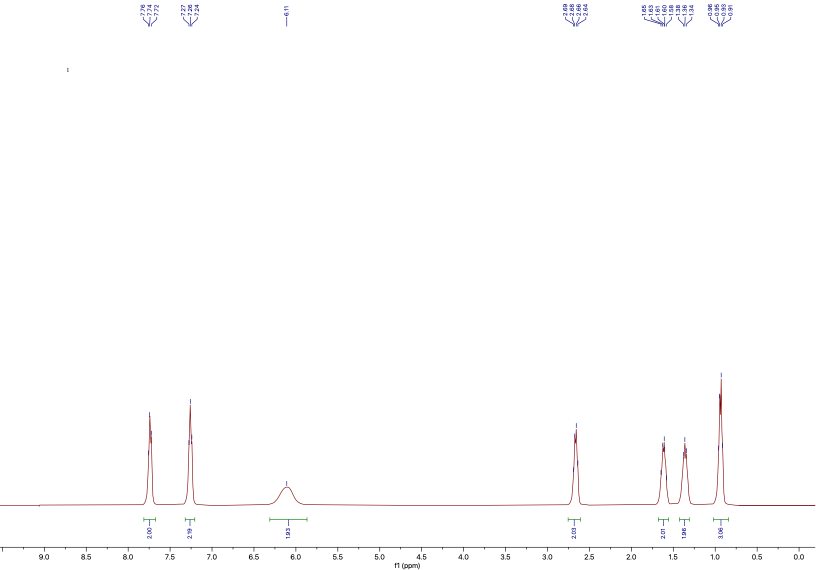

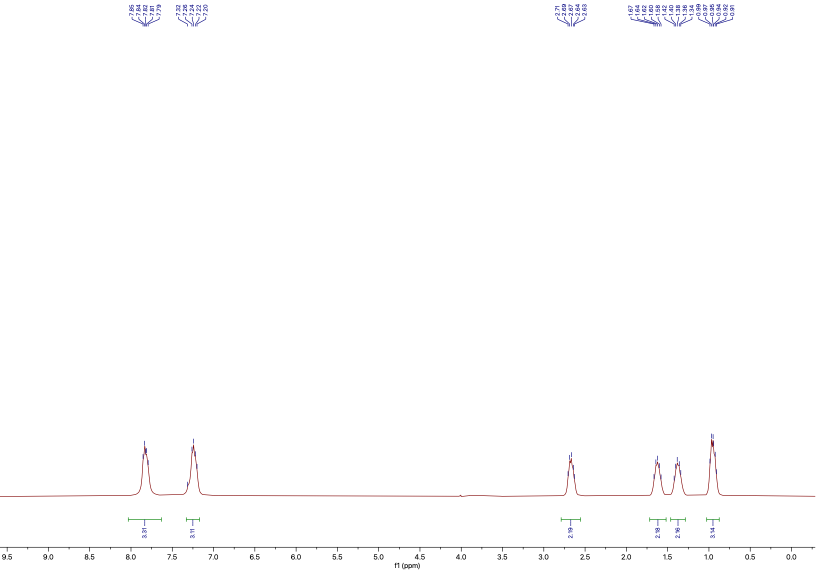

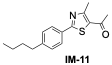

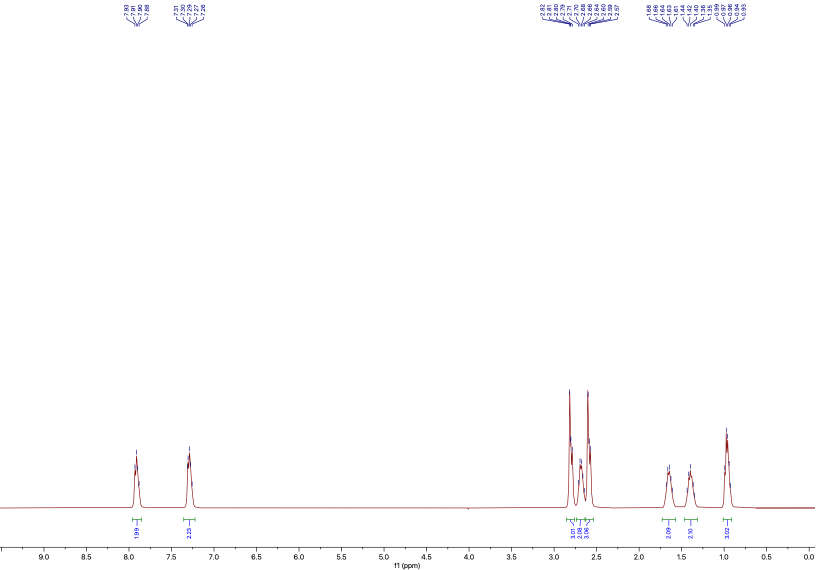

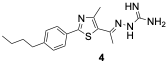

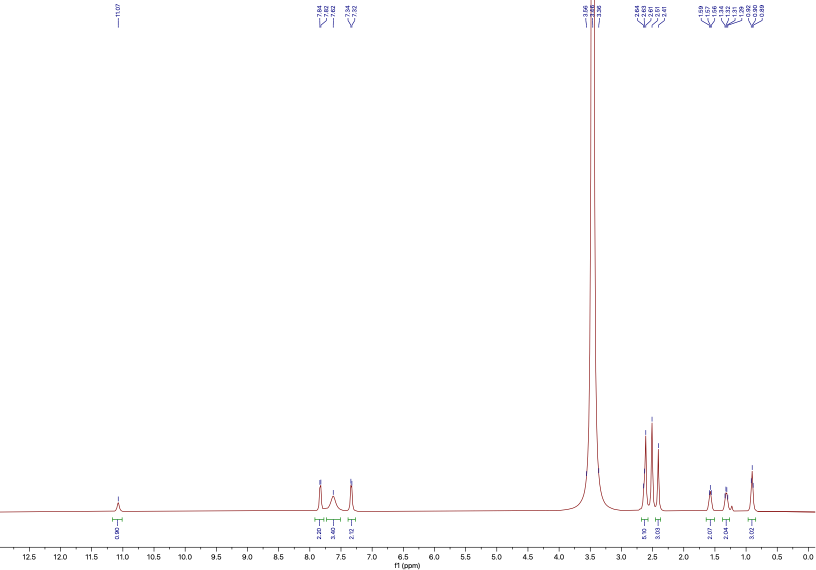

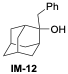

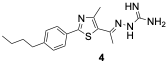

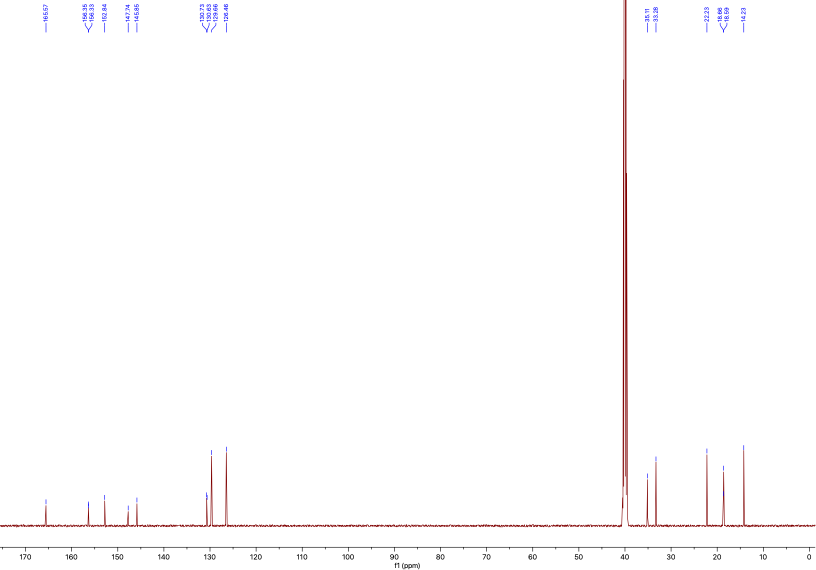

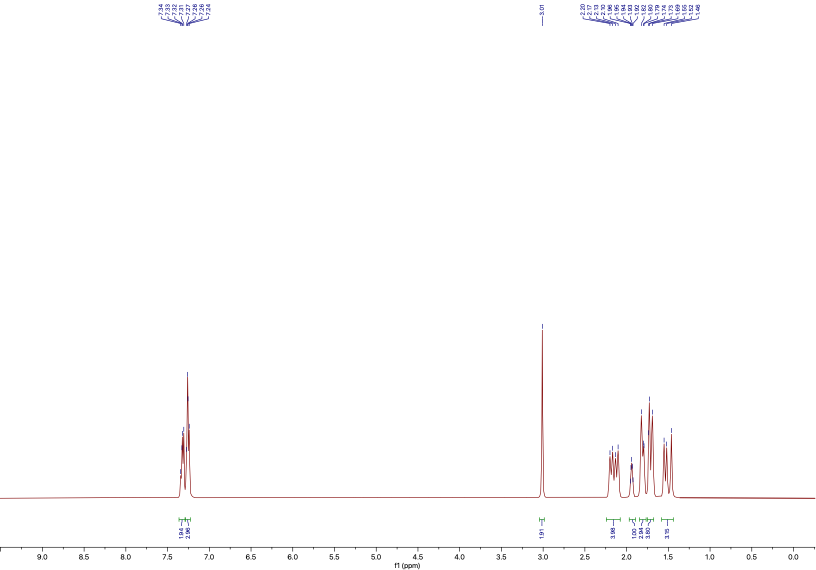

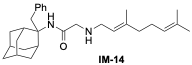

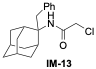

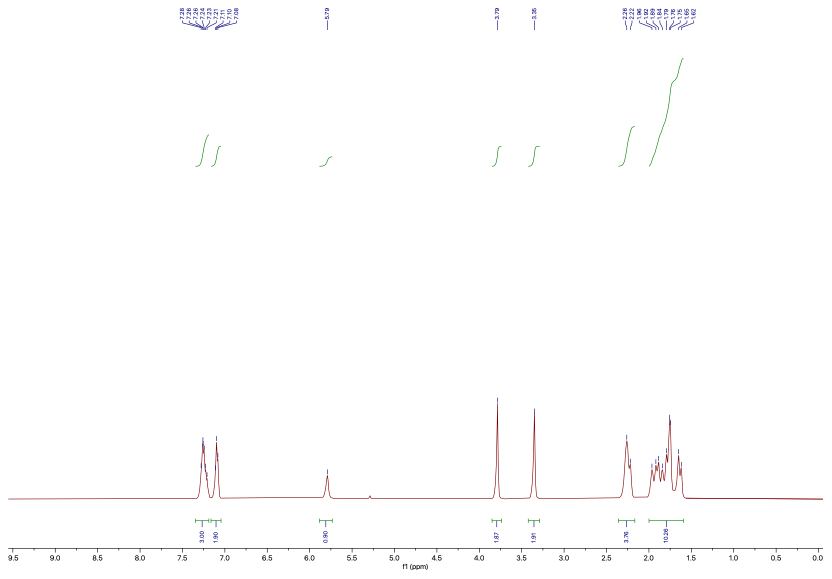

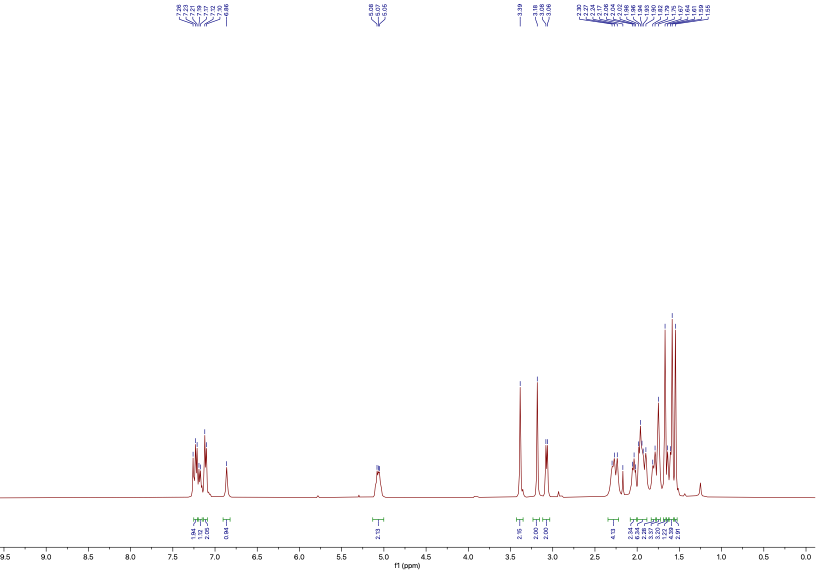

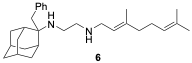

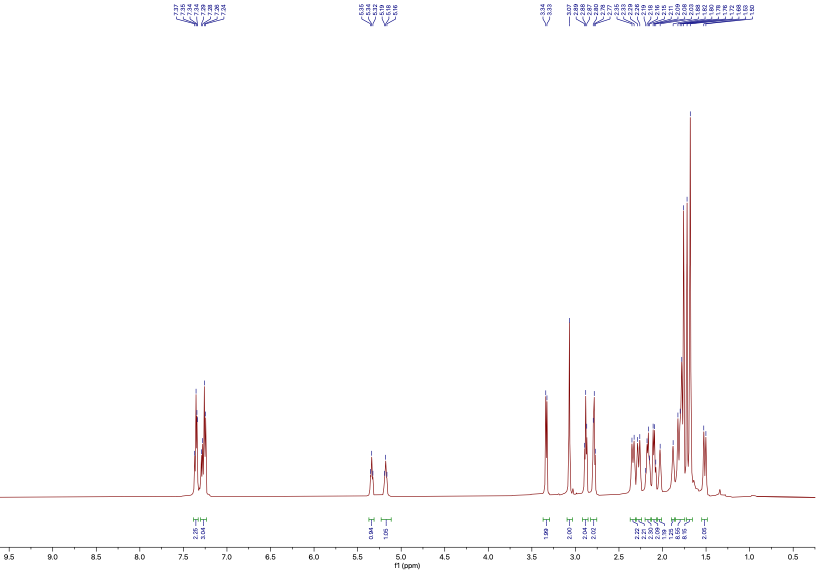

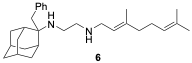

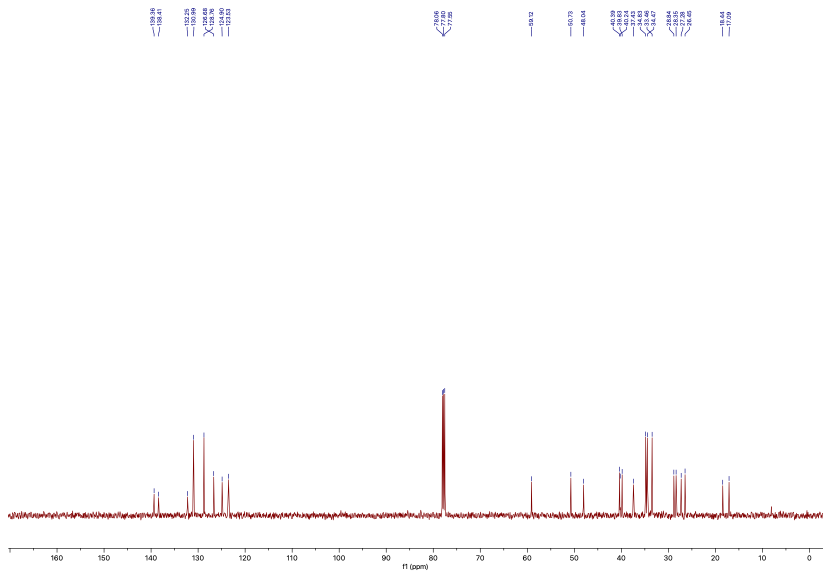

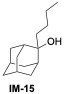

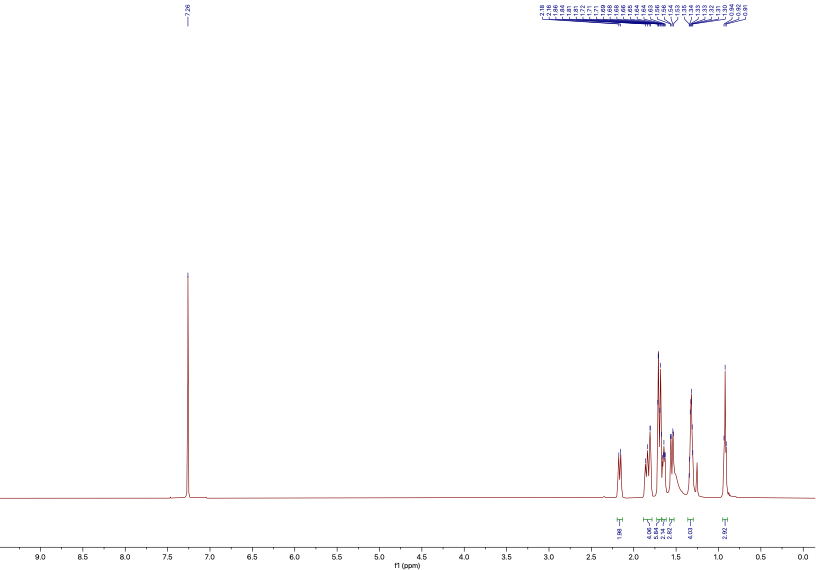

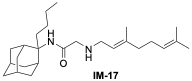

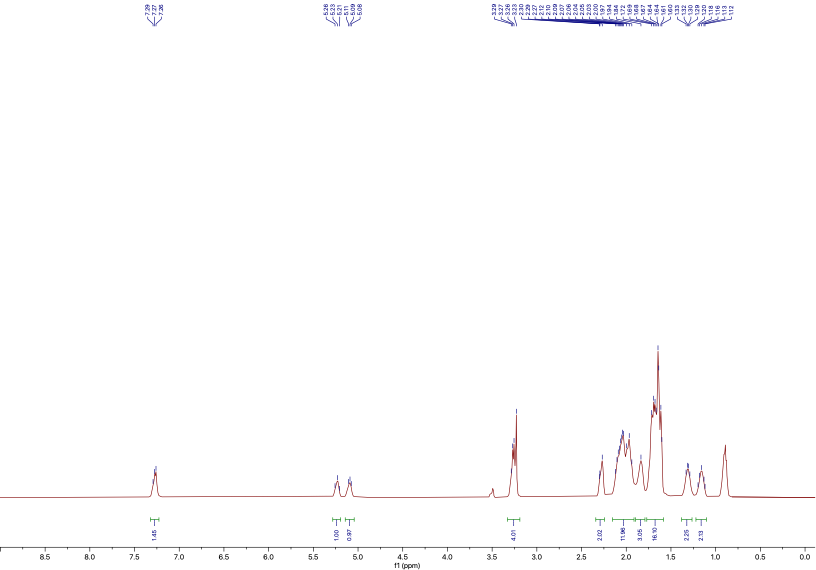

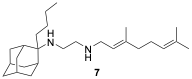

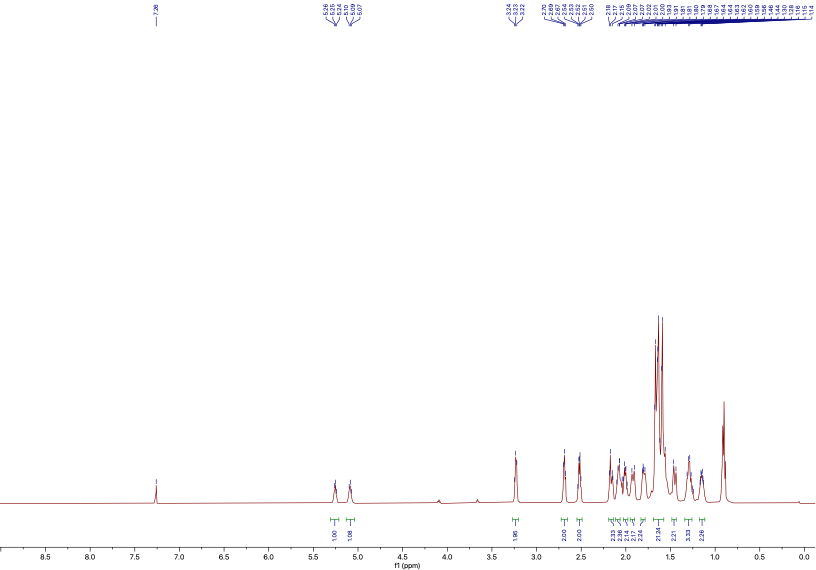

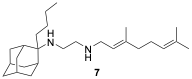

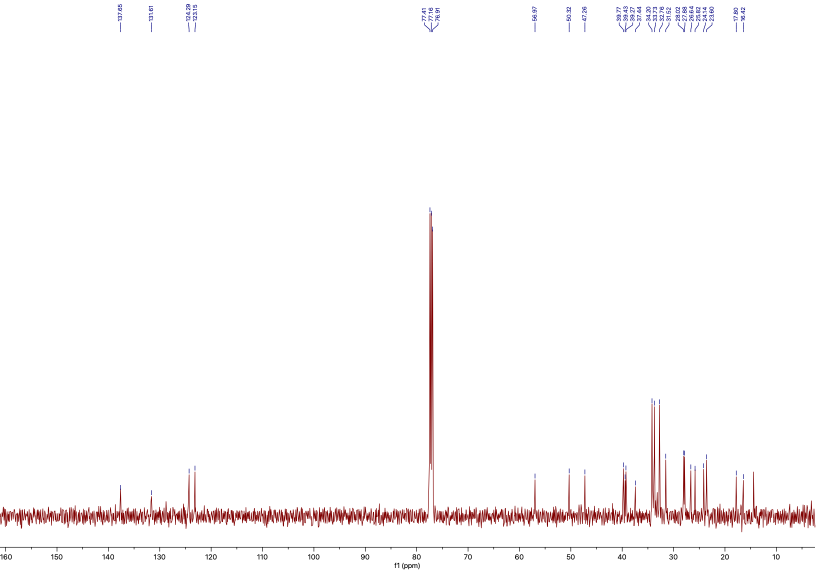

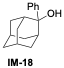

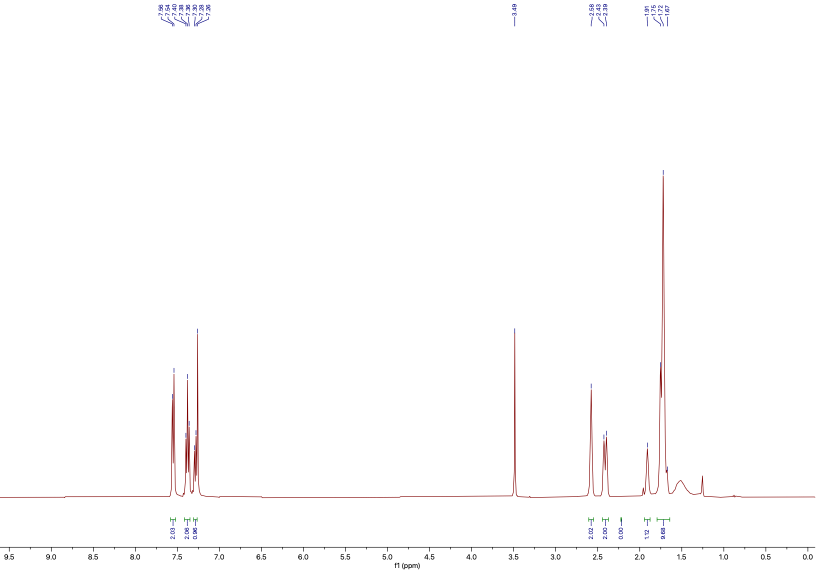

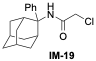

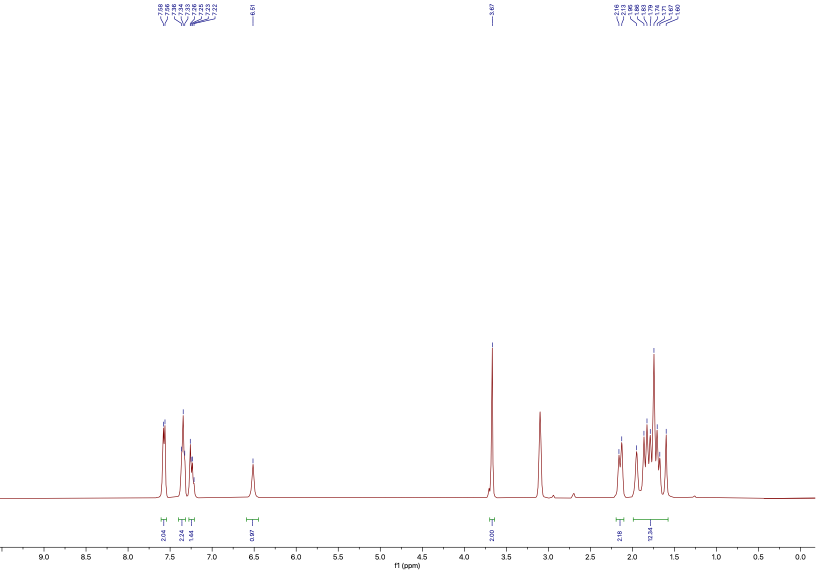

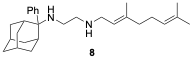

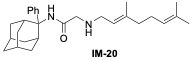

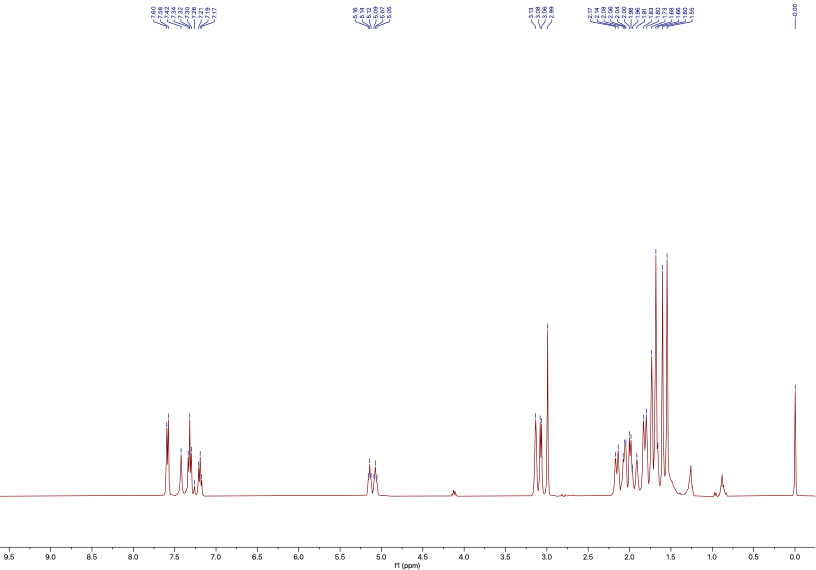

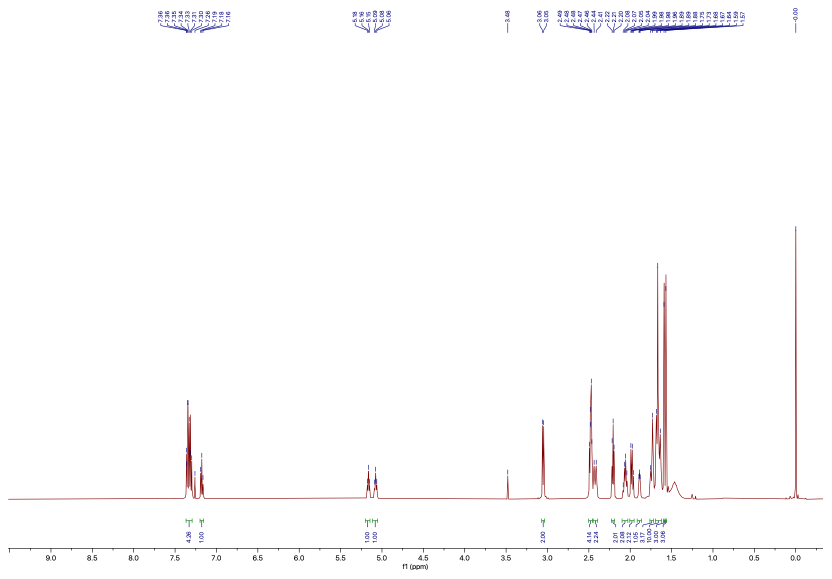

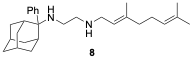

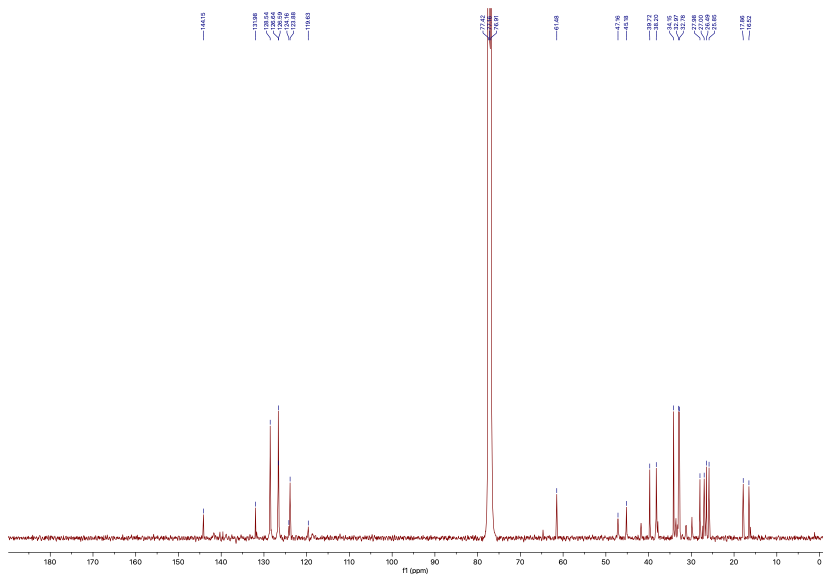

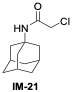

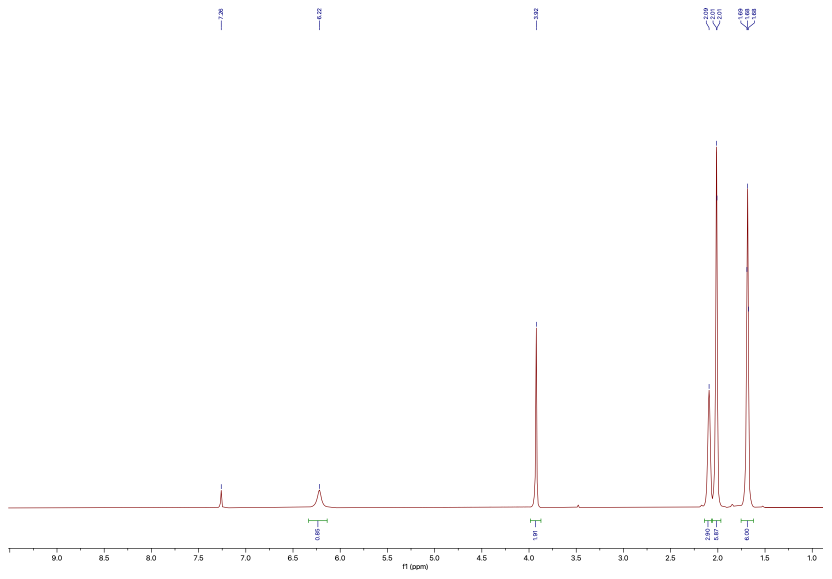

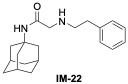

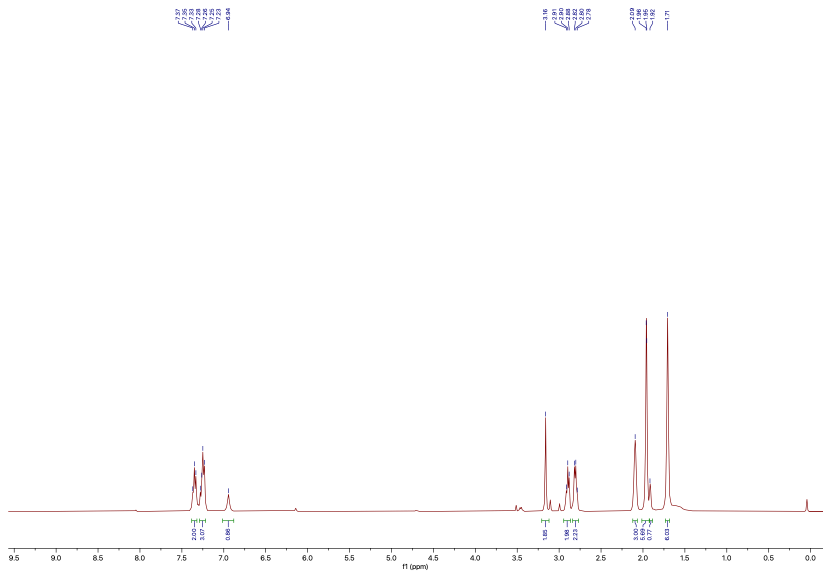

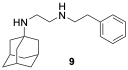

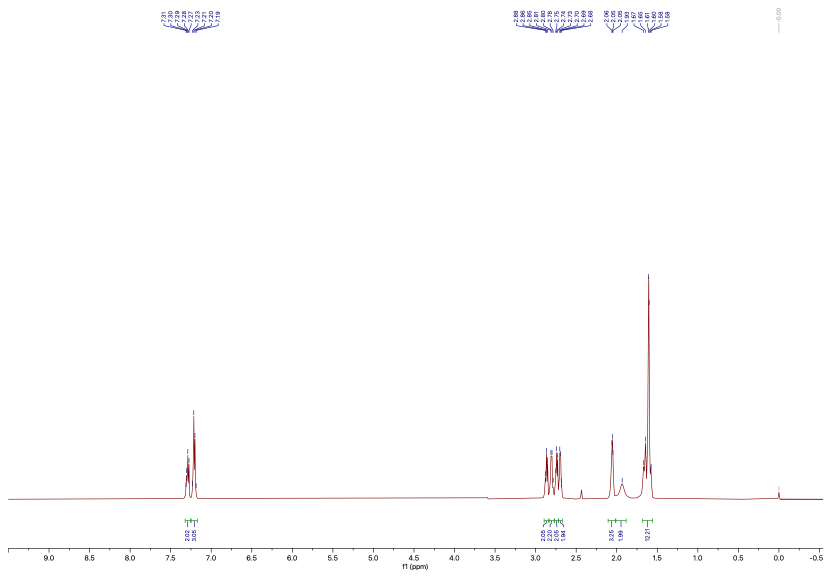

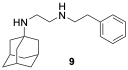

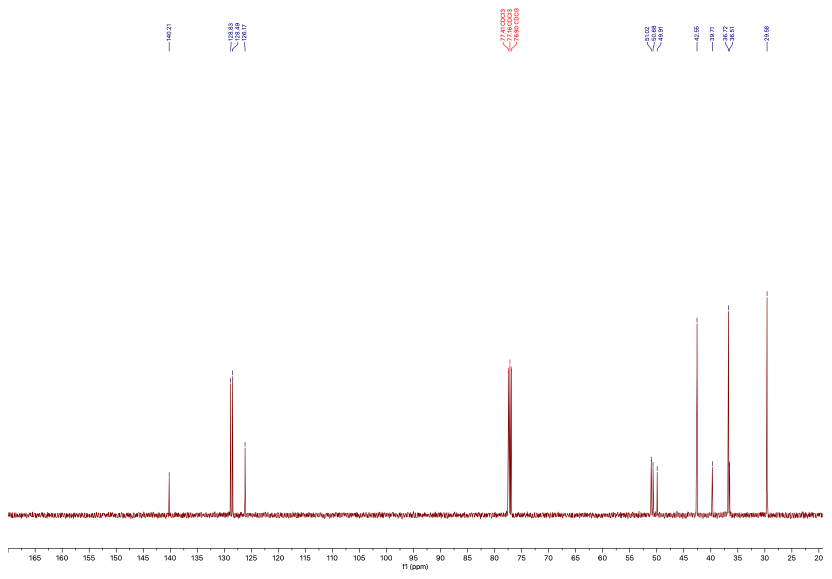

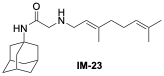

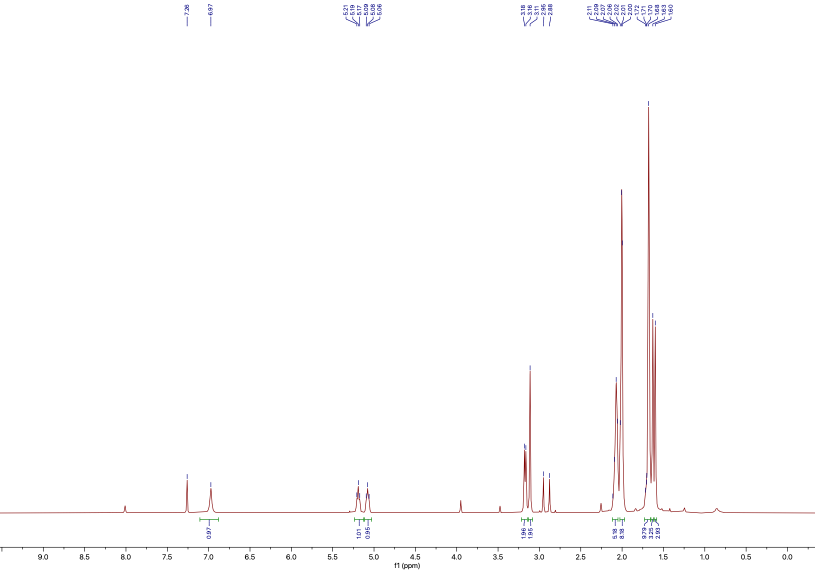

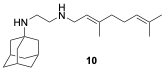

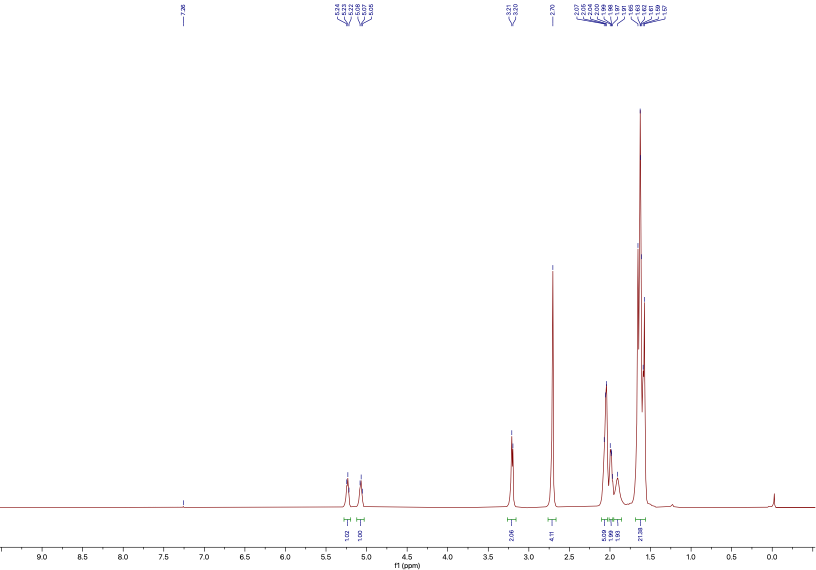

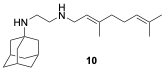

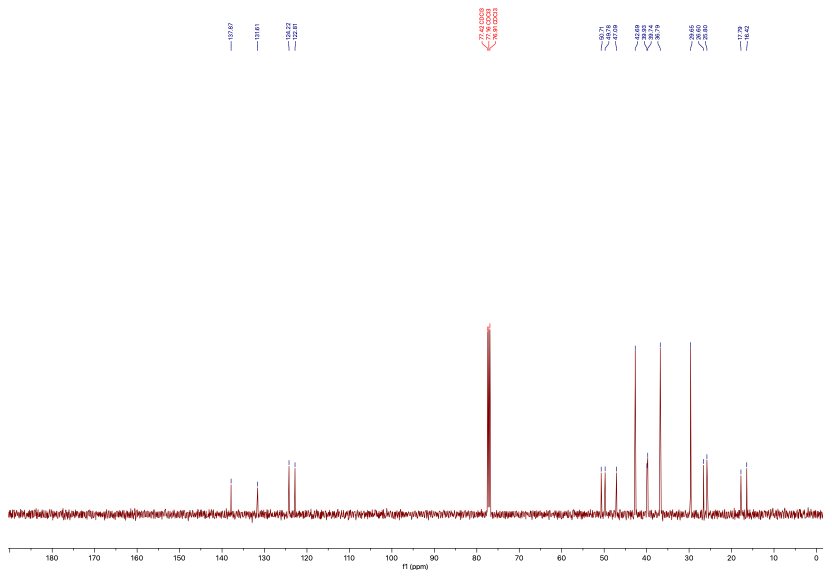

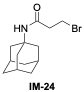

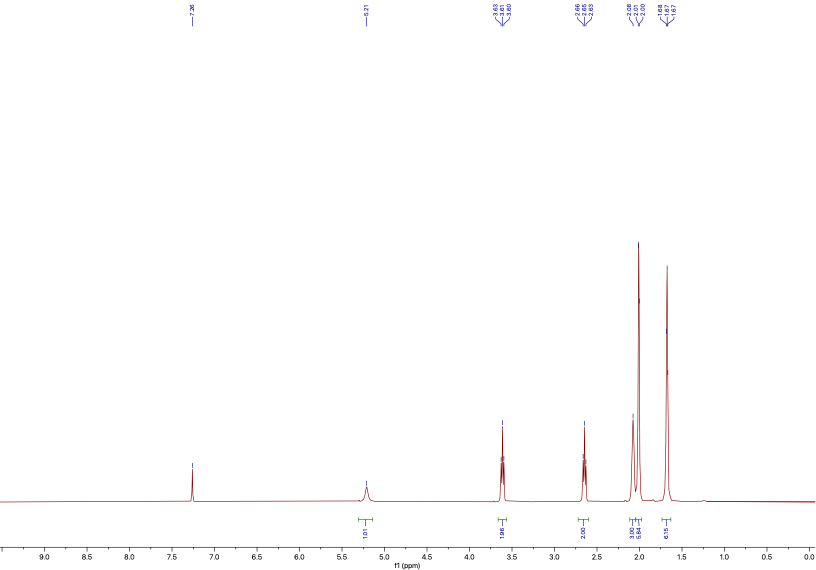

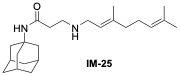

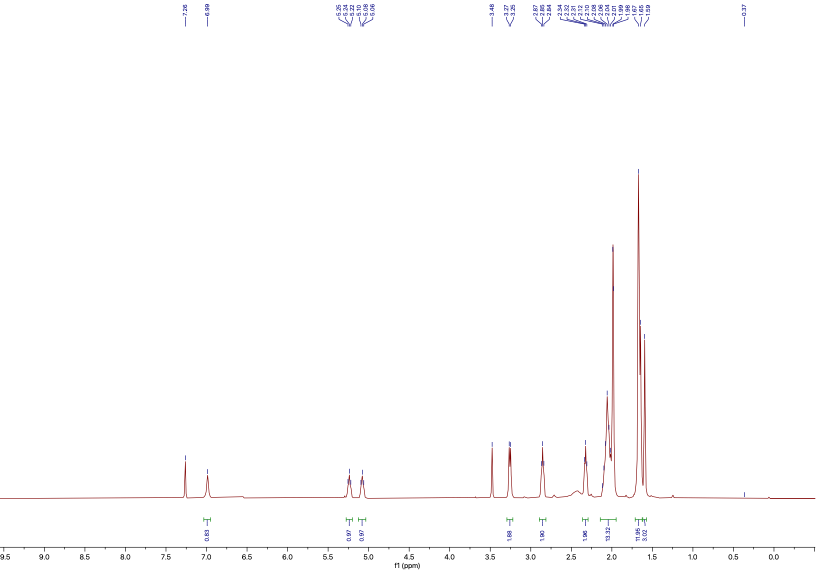

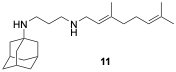

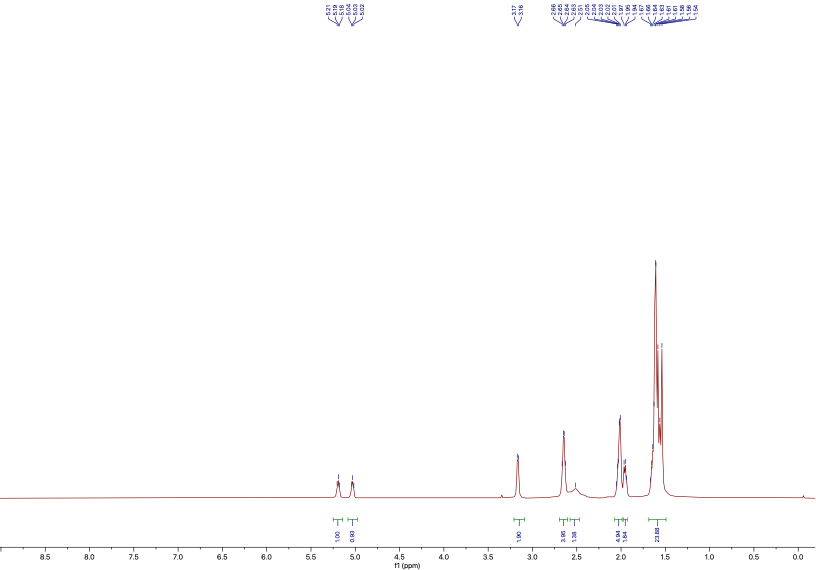

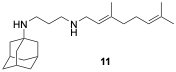

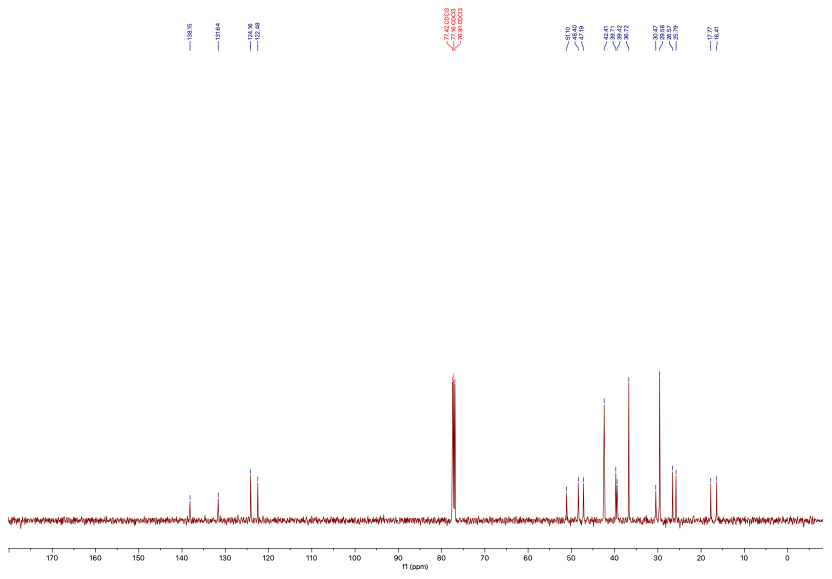

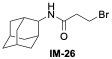

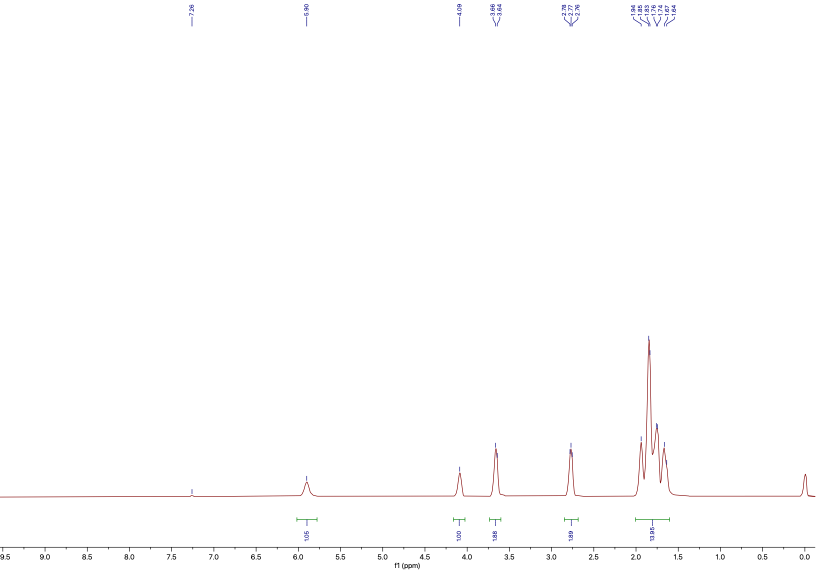

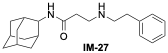

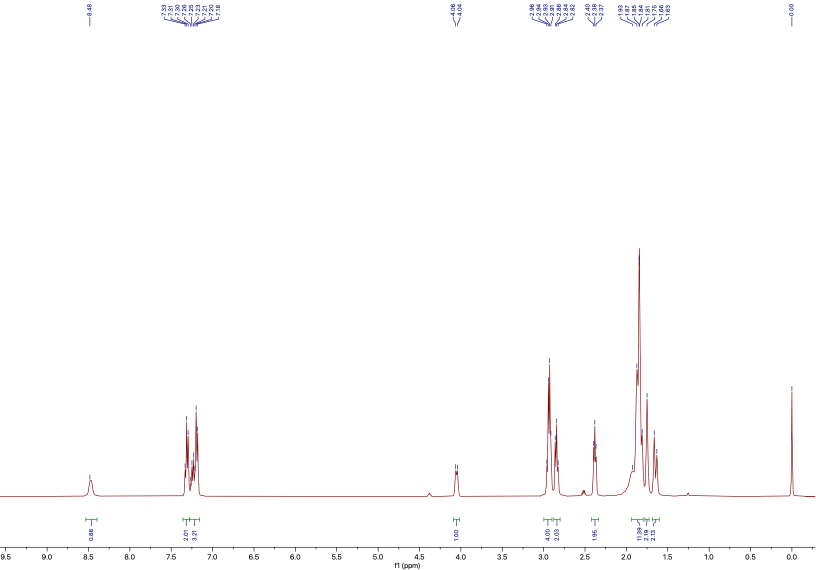

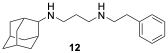

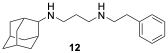

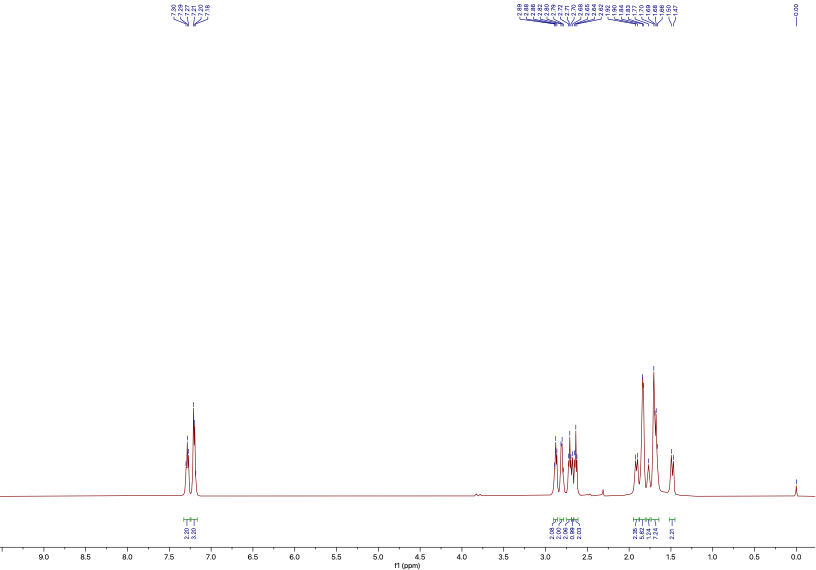

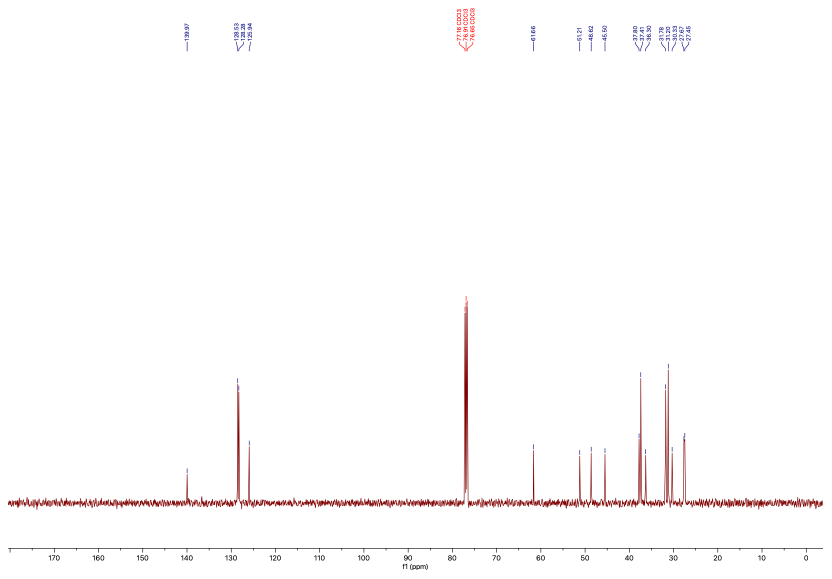

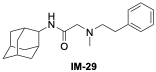

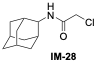

**qNMR spectra**

**Table S1.** SMILES of the compounds

| **Compound** | **Structure** | **SMILES** |
| --- | --- | --- |
| **1** |  | C1(CCCCC1)C2=CC3=C(C=C2)NC4=C3CCCC4NCCC5=CC=CC=C5 |
| **2** |  | C1(C2=CC3=C(C=C2)NC4=C3CCCC4NCCC5=CC=CC=C5)=CC=CC=C1 |
| **3** |  | C1(C2=CC3=C(C=C2)OC4=C3CCCC4NCCC5=CC=CC=C5)=CC=CC=C1 |
| **4** |  | CC1=C(/C(C)=N/NC(N)=N)SC(C2=CC=C(CCCC)C=C2)=N1 |
| **5** |  | C/C(C)=C/CC/C(C)=C/CNCCNC1[C@H]2C[C@@H]3C[C@@H](C[C@H]1C3)C2 |
| **6** |  | C/C(C)=C/CC/C(C)=C/CNCCNC1(CC2=CC=CC=C2)[C@H]3C[C@@H]4C[C@@H](C[C@H]1C4)C3 |
| **7** |  | C/C(C)=C/CC/C(C)=C/CNCCNC1(CCCC)[C@H]2C[C@@H]3C[C@@H](C[C@H]1C3)C2 |
| **8** |  | C/C(C)=C/CC/C(C)=C/CNCCNC1(C2=CC=CC=C2)[C@@H]3C[C@H](C[C@H]1C4)C[C@H]4C3 |
| **9** |  | C/C(CC/C=C(C)/C)=C\CNCCNC12C[C@@H]3C[C@H](C1)C[C@H](C2)C3 |
| **10** |  | C/C(CC/C=C(C)/C)=C\CNCCNC12C[C@@H]3C[C@H](C1)C[C@H](C2)C3 |
| **11** |  | C/C(CC/C=C(C)/C)=C\CNCCCNC12C[C@@H]3C[C@H](C1)C[C@H](C2)C3 |
| **12** |  | [C@@H]1(C[C@@H]2C3NCCCNCCC4=CC=CC=C4)C[C@H](C2)C[C@H]3C1 |
| **13** |  | CN(CCC1=CC=CC=C1)CCNC2[C@@H]3C[C@H](C[C@H]2C4)C[C@H]4C3 |
| **14** |  | C/C(CC/C=C(C)/C)=C\CN(C)CCCNC1[C@@H]2C[C@H](C[C@H]1C3)C[C@H]3C2 |
| **15** |  | C/C(CC/C=C(C)/C)=C\CN(C)CCNC12C[C@@H]3C[C@H](C1)C[C@H](C2)C3 |
| **16** |  | C/C(CC/C=C(C)/C)=C\CNCCNC1CC2CCC1C2 |
| **17** |  | C/C(CC/C=C(C)/C)=C\CNCCCNC1[C@@H]2C[C@H](C[C@H]1C3)C[C@H]3C2 |
| **18** |  | C/C(C)=C/CC/C(C)=C/CN(C)CCNC1[C@@H]2C[C@H](C[C@H]1C3)C[C@H]3C2 |
| **19** |  | [C@@H]1(C[C@@H]2C3NCCNCCC4=CC=CC=C4)C[C@H](C2)C[C@H]3C1 |
| **20** |  | C1(CNCCNC2[C@@H]3C[C@H](C[C@H]2C4)C[C@H]4C3)=CC=CC(OC5=CC=CC=C5)=C1 |
| **21** |  | [C@@H]1(C[C@@H]2C3NCCNCCC4CCCCC4)C[C@H](C2)C[C@H]3C1 |
| **22** |  | C1(CNCCNC2[C@@H]3C[C@H](C[C@H]2C4)C[C@H]4C3)=CC=CC=C1 |
| **23** |  | C/C(C)=C/CNCCNC1[C@@H]2C[C@H](C[C@H]1C3)C[C@H]3C2 |
